# Direct interaction of ribosomes with postsynaptic proteins gives rise to a privileged local synaptic translatome

**DOI:** 10.64898/2026.02.27.708433

**Authors:** Ashley M. Bourke, Marianna Massari, Georgi Tushev, Mei Wu, Kristina Desch, Sara Guerreiro Mota, Anja Staab, Elena Ciirdaeva, Julian D. Langer, Fan Liu, Erin M. Schuman

## Abstract

Ribosomes and thousands of mRNAs are localized near synapses to support local protein synthesis. Little is known, however, about how ribosomes are positioned and maintained in dendritic spines– the primary postsynaptic sites of excitatory neurotransmission. Here, using proximity labeling-mass spectrometry, we mapped the interactome of postsynaptic ribosomes, and discovered an unexpected interaction with AMPA receptor complex proteins. Co-immunoprecipitation and crosslinking mass spectrometry using rat cortical synaptosomes showed a direct, mRNA-independent interaction between postsynaptic proteins and intact ribosomes. Immunoprecipitation-ribosome profiling (IP-Ribo-seq) revealed not only the complete synaptic translatome but also that the AMPA receptor-associated subpopulation of synaptic ribosomes preferentially translates mRNAs encoding proteins related to post-synaptic density (PSD) scaffolding and cytoskeletal dynamics. Translation of one of the mRNA targets, *Camk2a*, was reduced in spines following ER sequestration of endogenous GluA1. Together, these results reveal a role for PSD proteins in positioning ribosomes near the postsynaptic membrane, providing a mechanism to couple synaptic activity with the local production of proteins needed for structural remodeling.

## Main Text

The large volume and elaborate morphology of a neuron’s dendritic and axonal arbor pose challenges for meeting proteostatic demands. It is estimated that the single neuron possesses more than 30 billion proteins (*1*) while ∼ 400,000 protein molecules reside in a single dendritic spine (*1*). Neuronal proteins have typical half-lives in the range of 5-10 days (*2*, *3*) resulting in protein turnover rates of ∼ 500,000 proteins per minute in an individual cell. To help meet this proteostasis challenge, mRNAs and ribosomes are trafficked to dendrites and axons where mRNAs can be translated locally (*4–6*). Many studies have identified a large and functionally diverse population of mRNAs present in dendrites and axons (*7–11*) and even growth cones (*12*). Similarly, early electron microscopy studies observed ribosomes near synapses in primate motoneurons (*13*) or within the synapse itself, close to the postsynaptic density (*14–16*). More recently, super-resolution microscopy methods have quantified the abundance of ribosomes at synapses, detecting an average of two translating ribosomes per synapse (*17*) and have estimated that synaptic ribosome abundance is sufficient to maintain the synaptic proteome (*1*). Not only are mRNAs and ribosomes present near synapses, their abundance is regulated by plasticity. For example, early studies demonstrated the activity-dependent localization of the immediate early gene *Arc* in activated synaptic laminae of the dentate gyrus (*18*) and more recent work has demonstrated the recruitment of mRNAs to spines following stimulation or plasticity (*19*). Following long-term potentiation (*16*, *20*, *21*) or behavioral learning (*22*), more polyribosomes were detected in dendritic spines.

As speculated in the early EM studies (*13*, *14*) the presence of ribosomes near synapses potentially allows them to respond to local signals. Indeed, during development, ribosomes associate with transmembrane adhesion and signaling receptors in spinal commissural axons (*23*) and retinal growth cones (*24*); stimulation with the appropriate ligand leads to a dissociation of the ribosome and mRNA translation. It is unknown, however, if there are mechanisms to position the ribosome in mature postsynaptic compartments to give it local access to synaptic information. Here we exploit advances in microscopy, proximity labelling, ribosome profiling, mass spectrometry (MS) and crosslinking MS to study the proteins that interact with ribosomes at mature synapses and the consequences of these interactions. We found that, within dendritic spines, assembled ribosomes associate with the receptors for the excitatory neurotransmitter glutamate and associated molecules in the postsynaptic density. This association leads to the enriched translation of a subset of mRNAs that encode for important synaptic complex members. The mis-localization of the glutamate receptor leads to a reduced translation of these target mRNAs. These data indicate that there is precision to the spatial targeting of synaptic translation machinery, making it well-poised to rapidly respond to synaptic signals and adjust translational output according to the needs of the synapse.

### The synaptic ribosome protein interactome

To identify the interaction partners of ribosomes at synapses we developed a pipeline that includes subcellular APEX proximity labelling, ribosome enrichment and downstream mass spectrometry (‘PL-MS’; Fig. 1A). We created 3 different APEX constructs targeted to the nuclear membrane (APEX-NUC; SENP2), the cytosol (APEX-CYTO), or to synapses (APEX-SYN, Homer1c) (Fig. 1A,B; fig. S1). Each APEX construct, introduced via an adeno-associated virus (AAV) vector, primarily localized to the intended compartment (nucleus, cytosol or synapse) and following a brief (1 min) H_2_O_2_ pulse gave rise to biotinylated protein that was also concentrated in the intended APEX-targeted region (Fig. 1B). To capture ribosomal and ribosome-associated proteins within each compartment, we first enriched ribosomes by sucrose cushion ultracentrifugation (*25*) before performing streptavidin pulldown of biotinylated proteins (Fig. 1A,C, fig. S3) followed by DIA mass spectrometry. As expected, the ribosome-enriched input fractions were very similar across conditions (fig. S2A-C), with compartment-specific differences revealed only in the eluate fractions (fig. S2D). In the eluates, we identified 1609, 1851, and 1869 high-confidence ribosome-interacting proteins in the ‘nuclear’, ‘cytosolic’ and ‘synaptic’ fractions, respectively (fig. S2F), defined by their enrichment over no-APEX controls (fig. S2E; table S1). The majority (64%-75%) of the ribosome interacting proteins were shared across compartments; nevertheless, there were subsets of compartment-specific enriched interactors. For example, neuronal ribosomes near the nucleus were enriched for interactors that included splicing factors, nuclear pore complex components and nuclear-enriched RNA-binding proteins (fig. S2G; table S2). Neuronal cytoplasmic ribosomes, by contrast, were enriched for several RNA-binding proteins associated with dendritic ribonucleoprotein granules and their transport (fig. S2G; table S2).

**Fig. 1.**
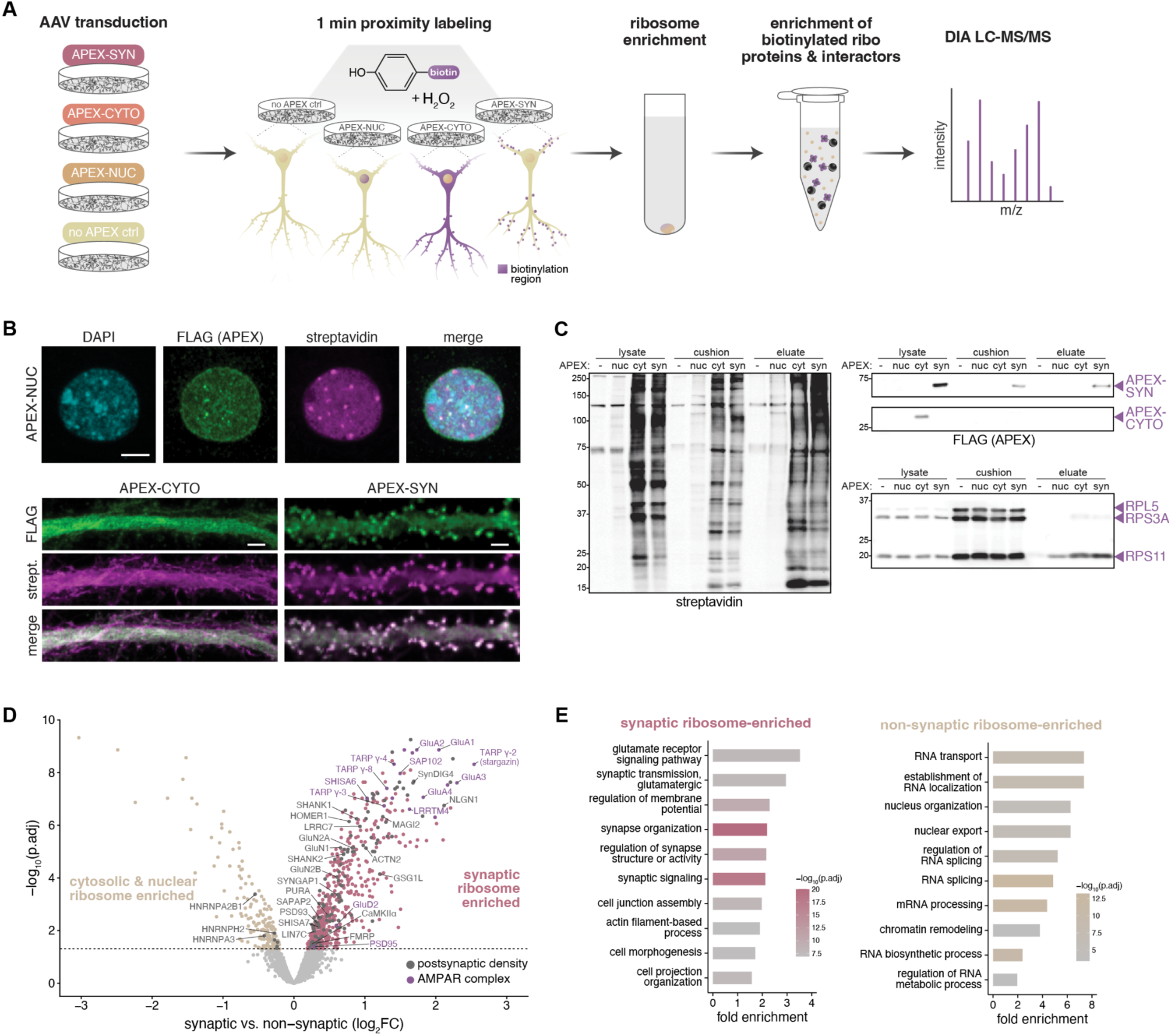
The synaptic ribosome interactome. (**A**) Schematic of proximity labeling-mass spectrometry (PL-MS) workflow. (**B**) Representative confocal images of the localization and biotinylation patterns of APEX-NUC (top), APEX-CYTO (bottom left), and APEX-SYN (bottom right) after 1 minute of labeling in primary rat hippocampal neurons. Scale bars, 4 µm. (**C**) Western blot images from one of five experimental replicates. (**D**) Volcano plot of differentially expressed proteins in the synaptic (n=564) and non-synaptic (combined cytosolic and nuclear, n=163) ribosome interactomes (FDR<0.05). Significantly enriched proteins associated with the GO terms “postsynaptic density” and “AMPA glutamate receptor complex” are colored in dark gray and plum, respectively. For proteins with multiple isoforms enriched, only the canonical isoform is labeled. Only selected “postsynaptic density” proteins are labeled. (**E**) GO overrepresentation analysis of enriched synaptic or non-synaptic ribosome-associated proteins. Shown are the top 10 enriched GOBP terms.

To resolve with stringency the protein interactors characteristic of ribosomes at synapses, we compared the combined cytosolic and nuclear ribosome interactomes with the synaptic ribosome interactome (Fig. 1D; table S3). We identified 564 significantly enriched synaptic ribosome interacting proteins. Strikingly, amongst the most enriched synaptic ribosome–enriched proteins were members of the major excitatory neurotransmitter receptor complex, including all four core subunits of the AMPA receptor as well as seven auxiliary subunits (Fig. 1D; fig. S2H). Gene ontology (GO) enrichment analysis revealed that synaptic ribosome interactors were strongly enriched for ‘glutamate receptor signaling pathway’, ‘synapse organization’, ‘synaptic signaling’, and ‘regulation of synapse structure or activity’, whereas non-synaptic ribosome interactors were enriched for terms more broadly associated with RNA localization and processing (Fig. 1E; fig. S2I; table S3). We note that fewer than 10% of the ribosome interactors exhibited N-terminal detection bias (mean peptide position <0.3; fig. S2J; see Methods), indicating that the majority – including AMPA receptor complex proteins – are bona fide ribosome-associated rather than nascent chains.

### Synaptic ribosomes interact with AMPA receptors

To independently determine the AMPA receptor-ribosome interaction, we developed a reciprocal co-immunoprecipitation strategy targeting endogenous ribosomes and AMPA receptor associated ribosomes from rat cortical synaptosomes, coupled to mass spectrometry (‘IP-MS’; Fig. 2A; fig. S4A-B). For immunoprecipitation we used an antibody against either the large ribosomal subunit protein RPLP0 (associated with mature ribosomes; (*26*)) or the AMPA receptor subunit GluA1 (*27*). MS analysis of the synaptic RPLP0 and GluA1 IP eluates yielded ∼694-1055 and 2350-3427 unique proteins per replicate, respectively, whereas the corresponding isotype control eluates contained many fewer proteins (Fig. 2B; fig. S4C,D). Of the 564 proteins (529 unique gene identifiers) enriched in the APEX PL-MS synaptic ribosome interactome, 97 were also found in both the RPLP0 IP and GluA1 IP interactomes (Fig. 2C; table S4), including AMPA receptor subunits GluA1-3, the auxiliary subunits stargazin and TARP γ-8, and the AMPA receptor-associated scaffold protein SAP102 (Fig. 2D,E). GO analysis identified ‘ionotropic glutamate receptor binding’ among the top 10 enriched terms for the RPLP0 IP and ‘structural constituent of the ribosome’ for the GluA1 IP (Fig. 2E,F; table S4). Together, these findings demonstrate that two complementary biochemical strategies (PL-MS and IP-MS) capture AMPA receptors as interactors of synaptic ribosomes in both primary cultured neurons and cortical tissue.

**Fig. 2.**
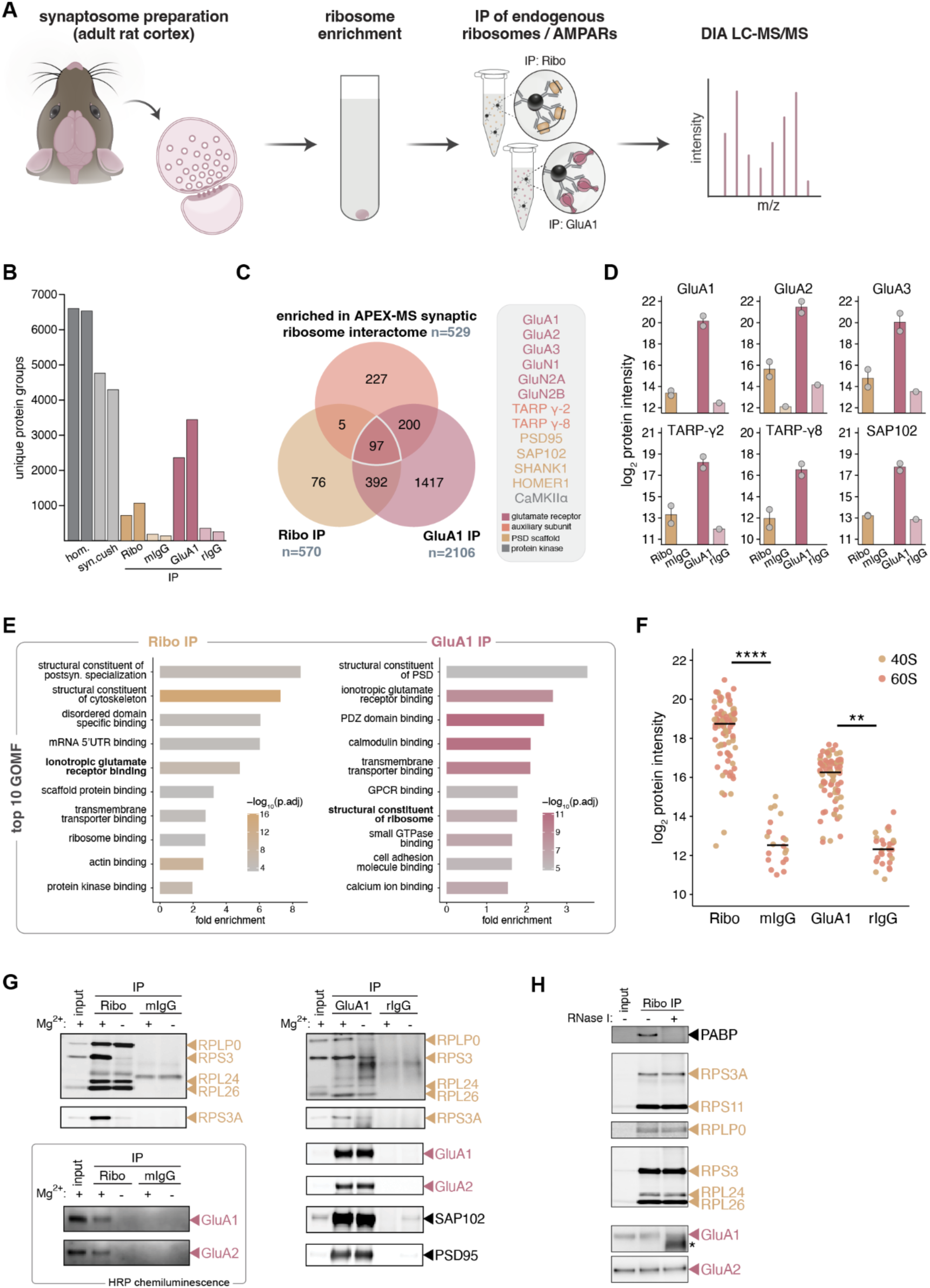
Synaptic ribosomes interact with AMPA receptor complex proteins. (**A**) Schematic of immunoprecipitation-mass spectrometry (IP-MS) workflow. (**B**) Number of protein groups quantified per replicate from the tissue homogenate, synaptic fraction cushion and IP eluate fractions. (**C**) Distinct biochemical strategies (PL-MS and IP-MS) capture AMPA receptors as interactors of synaptic ribosomes. IP proteins were filtered based on uniqueness to the IP or >3-fold enrichment compared to the corresponding isotype control. Protein groups were collapsed to unique gene identifiers prior to overlap analysis. Values correspond to the number of unique gene identifiers. *Right:* Proteins found in the union of all three interactomes that are associated with the GO terms “AMPA glutamate receptor complex” or “Glutamate receptor signaling pathway”. (**D**) Relative intensities of six selected proteins that are enriched in the ribosome IP and GluA1 IP eluate fractions compared to the corresponding isotype control and are associated with the GO term “AMPA glutamate receptor complex”. Error bars show mean ± SEM. (**E**) GO overrepresentation analysis for proteins in the ribosome IP and GluA1 IP eluate fractions. Shown are the top 10 enriched GOMF terms. (**F**) Average log_2_ protein intensities of ribosomal proteins (RPs) in the ribosome IP and GluA1 IP eluate fractions compared to the corresponding isotype control. Horizontal bars represent the median; **p<0.01; ****p<0.0001, Kruskal-Wallis test, Dunn’s multiple comparisons test. See table S2. (**G**) Western blot analysis of ribosomal proteins, AMPA receptor subunits and known AMPA receptor interactors after IP of ribosomes or GluA1, with on-bead washes performed with or without magnesium. Blots are from one of three experimental replicates. Input: 0.25 ug of synaptic cushion (0.5% of IP input), IP: 50% of IP eluate. (**H**) Western blot analysis of PABP, ribosomal proteins and AMPA receptor subunits after IP of ribosomes or GluA1 with and without RNase I treatment. Blots are from one of three experimental replicates. Input: 0.5 ug of synaptic cushion (1% of IP input), IP: 50% of IP eluate. Asterisk indicates a possible glycosylated or partially proteolyzed form of GluA1. Abbreviations: hom., homogenate; syn.cush, synaptic fraction cushion; GOMF, Gene Ontology Molecular Function; IP, immunoprecipitation; mIgG, mouse IgG; rIgG, rabbit IgG; PABP, poly(A)-binding protein.

To determine whether the AMPA receptor interactions, observed in both the IP and proximity-labelling data, require an intact assembled (80S) ribosome, we repeated the experiment with Mg^2+^-free washes, promoting dissociation of ribosomes into subunits after capture (*28*), and analyzed the eluates by Western blot (Fig. 2G, fig. S4F). As expected, RPLP0 recovery of 40S, but not 60S, proteins was markedly reduced following Mg^2+^-free washes. AMPA receptor complexes were recovered with RPLP0 only under standard (Mg^2+^-containing) conditions, and recovery of proteins from both ribosomal subunits in the GluA1 IP was likewise diminished upon ribosome disassembly. Detection of GluA2, SAP102, and PSD95 in the GluA1 IP was unaffected by Mg^2+^ removal, confirming that AMPA receptor complexes remained intact. Overall, these results suggest that AMPA receptors associate specifically with assembled ribosomes. Previous work in developing *Xenopus* axons showed that ribosomes associate with guidance cue receptors in an mRNA-dependent manner (*24*). To assess whether the AMPA receptor-ribosome interaction is similarly mediated by mRNA, we treated the synaptic fraction cushion with RNase I prior to RPLP0 IP (Fig. 2H). RNase I efficiently degraded mRNA while preserving rRNA integrity, as indicated by loss of poly(A)-binding protein (PABP) and stable recovery of ribosomal proteins in the RPLP0 IP. RNase I treatment did not, however, affect the ribosome-mediated IP of GluA1 or GluA2, suggesting that AMPA receptor-ribosome interactions are not dependent on mRNA.

### A subpopulation of postsynaptic ribosomes clusters near surface AMPA receptors

Can the AMPA receptor-ribosome interaction be visualized *in situ*? Ribosomes have been readily detected in dendrites and spines using light microscopy (*1*, *17*, *29–31*) (Fig. 3A,B). To resolve their nanoscale organization relative to surface AMPA receptors, we used stimulated emission depletion (STED) super-resolution imaging. We live-labeled endogenous surface GluA1 and co-stained for ribosomal protein RPS11 (which labels 80S ribosomes; (*17*)) in primary neurons expressing mVenus to delineate the dendritic spines, sites of excitatory synapses. For each spine, we quantified the nearest-neighbor distances between centers of segmented surface GluA1 and RPS11 puncta (Fig. 3C; fig. S5A,B) and normalized them to the density of the target population (GluA1-to-RPS11 by RPS11 density; RPS11-to-GluA1 by GluA1 density) to account for variability across spines. Distribution analyses revealed that surface GluA1 puncta were closer to ribosomes than expected under complete spatial randomness (CSR) (Fig. 3D,E). As expected, however, the reverse relationship was not observed for ribosomes relative to surface GluA1. This asymmetry is consistent with their known localization: surface AMPA receptors form nanoclusters in the spine head plasma membrane (*32–34*), whereas ribosomes are thought to be more broadly distributed throughout the cytoplasm of the head and neck, and can relocalize from the dendritic shaft to spines during long-term potentiation (*14*, *16*, *20*, *21*, *35*). In addition, the observed subset of spine-localized ribosomes that associate with surface AMPA receptors is in line with our observation that the ribosomal protein signal was ∼5.6-fold higher in the RPLP0 IP than in the GluA1 IP (Fig. 2F). To quantify the distance between surface GluA1 and the nearest ribosome we used 2D sum-projections of three-plane stacks and calculated a median separation of 125 nm (IQR: 70.7-175 nm; mean ± SD: 133.9 ± 98.2 nm; fig. S5B), a projection-limited lower bound on the true 3D distance. To test whether this proximity exceeded chance, we compared per-spine median distances with a simulated spatial null in which surface GluA1 coordinates were held fixed and RPS11 positions were randomized within each spine ROI (Fig. 3F,G). The measured surface GluA1 to RPS11 medians were significantly smaller than those simulated. Finally, we note that 81% of spines contained at least one ribosome within 100 nm of surface GluA1 (mean ± SD across cells: 0.81 ± 0.13; fig. S5C). Together, biochemical association and nanoscale imaging show that ribosomes are preferentially positioned near surface AMPA receptors in most spines, providing a spatial framework for localized translation at synaptic sites.

**Fig. 3.**
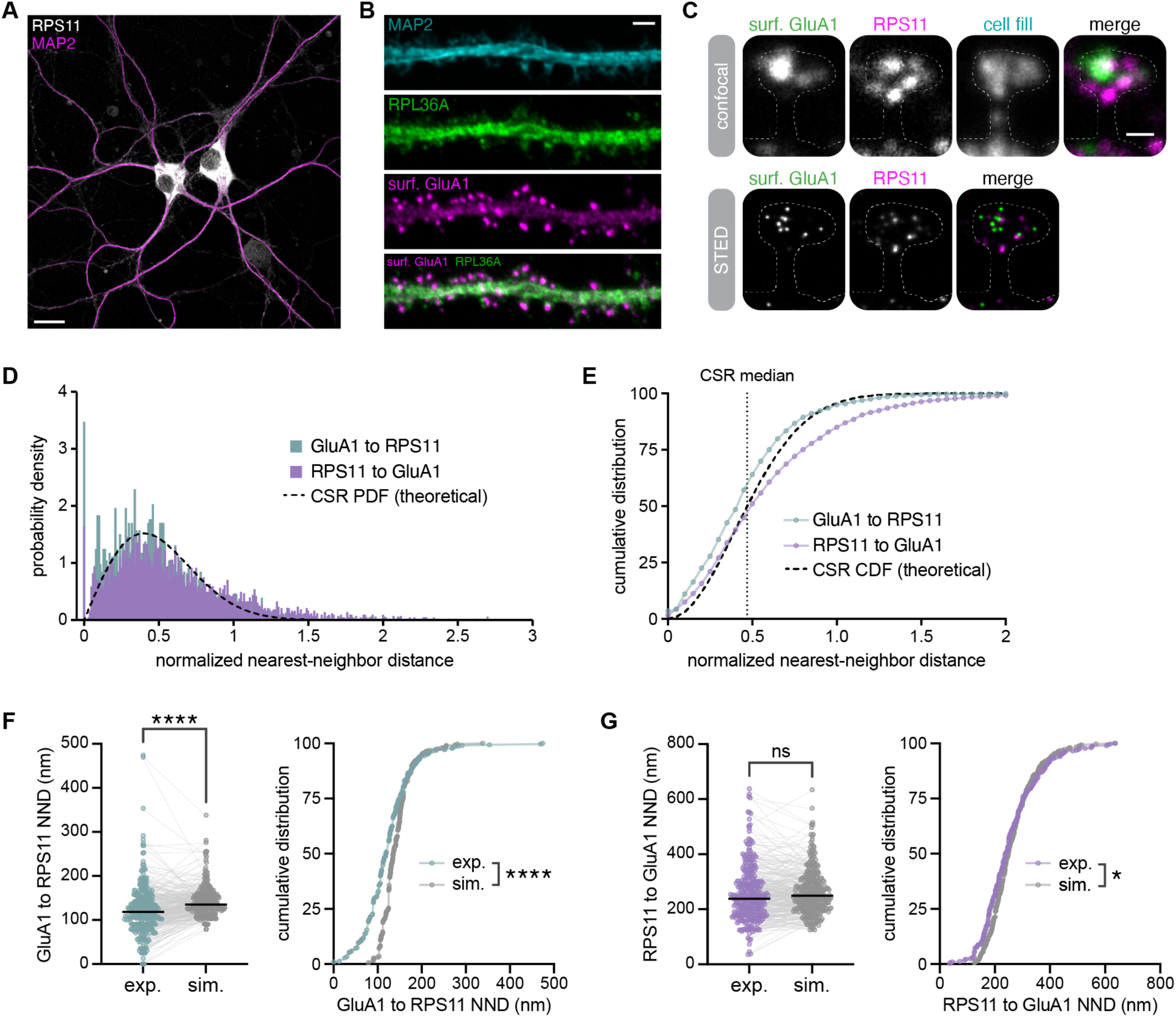
A subpopulation of synaptic ribosomes clusters near surface AMPA receptors. (**A**) Confocal image of primary hippocampal neurons immunostained for MAP2 (magenta) and RPS11 (white). Scale bar, 20 µm. (**B**) Confocal images of dendrites immunolabeled for surface GluA1 (magenta), MAP2 (cyan) and RPL36A (green). Scale bar, 2 µm. (**C**) Overview of STED imaging analysis. Scale bar, 0.5 µm. (**D**) Probability density (area = 1) of target-normalized nearest-neighbor distances. Teal: surface GluA1 to RPS11 (normalized by RPS11 density). Purple: RPS11 to surface GluA1 (normalized by surface GluA1 density). Black line: CSR PDF. Excess density at small distances indicates closer-than-random proximity. (GluA1 to RPS11: n=1524 distances (puncta); RPS11 to GluA1: n=4254 distances (puncta). (**E**) Empirical cumulative distributions of the same normalized distances. Black line: CSR CDF. Curves left/above CSR indicate closer-than-random proximity. Dashed vertical line marks the CSR median (0.469). (**F, G**) Median NND per spine compared with simulation values in which surface GluA1 object coordinates were held fixed and RPS11 objects were randomly repositioned within the spine ROI (100 randomizations per spine). For each spine, the simulated value equals the median across its 100 simulated medians. (**F**) Left: paired dot plot of per-spine median surface GluA1 to RPS11 NNDs (exp. vs sim.); lines connect the same spine. Horizontal bars represent the group median across spines. ****p<0.0001, two-tailed Wilcoxon matched-pairs signed rank test. Right: ECDF of per-spine medians (exp. vs sim.). A left/upward shift of the experimental curve indicates shorter medians than the simulated null. ****p<0.0001, two-tailed Kolmogorov-Smirnov test. (**G**) Left: paired dot plot of per-spine median RPS11 to surface GluA1 NNDs (exp. vs sim.); lines connect the same spine. Horizontal bars represent the group median across spines. p=0.1263, two-tailed Wilcoxon matched-pairs signed rank test. Right: ECDF of per-spine medians (exp. vs sim.). ns=not significant (p=0.0141), two-tailed Kolmogorov-Smirnov test; n=323 spines from 100 dendritic segments from 23 cells from three independent cultures. Abbreviations: CSR, complete spatial randomness; PDF, probability density function; CDF, cumulative distribution function; NNDs, nearest-neighbor distances; exp., experimental; sim., simulated.

### Identification of the synaptic and the GluA1-subsynaptic translatomes

Although synaptic ribosome-associated mRNAs have been identified (*24*, *36*, *37*), the full complement of mRNAs that are actively translated by ribosomes within cortical excitatory synapses has not yet been elucidated. We first focused on the global synaptic translatome, unambiguously identifying translating mRNAs with codon resolution by coupling our IP strategy to ribosome profiling (*38*) (Fig. 4A). Read mapping statistics and quality metrics confirmed the specificity and integrity of the ribosome footprints (fig. S6, S7), and proteomic analysis showed that the synaptic fraction was not enriched for proteins of contaminating compartments (fig.S8A). We detected 12,543 transcripts with counts per million (CPM) ≥ 1 in three of four synapse Ribo-seq replicates in either IP (RPLP0 or GluA1). Integration with RNA-seq data derived from the same synaptic fractions (fig. S8B,C) defined a core synaptic translatome of 12,283 mRNAs that were both detectable at the RNA level and actively translated (Fig. 4B). GO enrichment analysis revealed ‘regulation of synapse organization’, ‘regulation of postsynaptic membrane neurotransmitter receptor levels’, and ‘regulation of synapse plasticity’ among the functional classes of synaptic mRNAs undergoing active translation (fig. S8D).

**Fig. 4.**
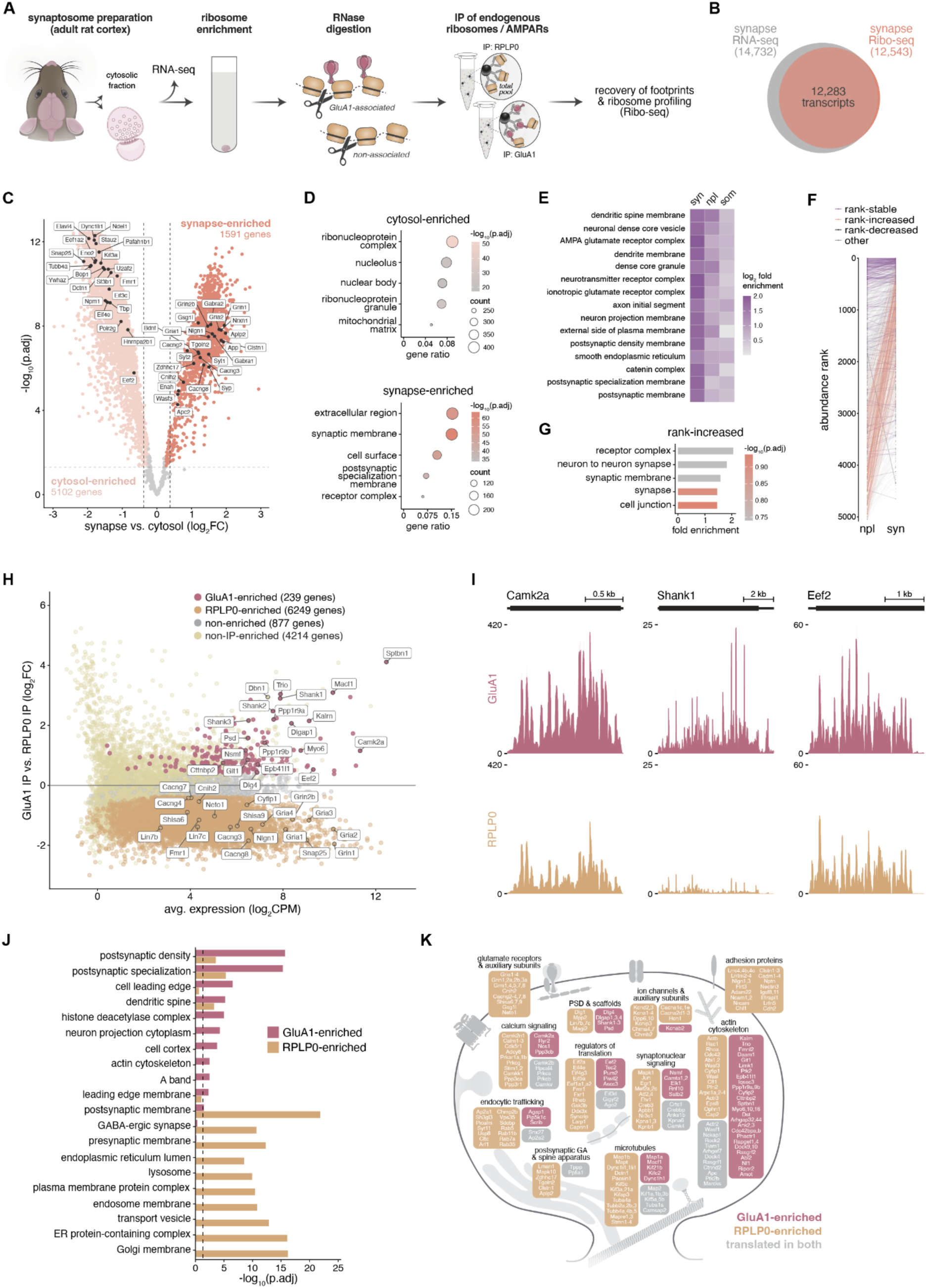
GluA1-associated ribosomes actively translate mRNAs encoding PSD scaffolding proteins and actin-scaffold adapters. (**A**) Immunoprecipitation-ribosome profiling (IP-Ribo-Seq) experimental workflow. (**B**) Overlap of detected transcripts in synapse RNA-seq and synapse Ribo-seq datasets. (**C**) Volcano plot comparing translational levels of 6,889 transcripts between synaptic and cytosolic compartments (RPLP0 IP from synaptic fraction vs. RPLP0 IP from cytosolic fraction). Significantly enriched transcripts are highlighted (FDR ≤ 0.05 and |log_2_FC| ≥ log_2_(1.3)). (**D**) Functional segregation of transcripts differentially translated between synaptic and cytosolic compartments. Shown are the top 5 enriched GOCC terms for synapse- and cytosol-enriched transcripts. (**E**) Functional comparison of synapse-enriched transcripts with neuropil- and somata-enriched transcripts from (*10*). Top 15 GOCC terms are shown, ranked by enrichment score (mean log_2_ fold enrichment). (**F**) Comparison of relative abundance ranks between synaptic and neuropil compartments. Paired dot plot showing footprint abundance ranks of synapse-enriched transcripts in the synaptic and neuropil translatomes. Transcripts with ≤5% rank change were classified as rank-stable. Transcripts with >40% upward or downward shifts from the neuropil to synaptic compartments were classified as rank-increased and rank-decreased, respectively. (**G**) Top 5 GOCC terms for rank-increased transcripts. (**H**) MA plot comparing the translational level of 7,365 transcripts between GluA1-associated ribosomes and the total synaptic ribosome pool. (**I**) Coverage profiles representing the average GluA1 IP (top) or Ribo IP (bottom) footprint coverage for candidate GluA1-associated ribosome (*Camk2a*, *Shank1*, *Eef2*) transcripts. The y-axis indicates the number of normalized reads. (**J**) Top 10 enriched GOCC terms for GluA1-vs RPLP0-enriched transcripts. The dashed line indicates -log10(FDR). (**K**) Schematic of a dendritic spine highlighting some of the transcripts preferentially translated by GluA1-associated ribosomes (rose) or the total synaptic ribosome pool (tan). Transcripts that are not enriched in either (i.e., translated in both) are labeled in gray.

To determine transcripts preferentially translated at synapses, we first compared the translatomes of the synaptic and cytosolic fractions obtained in our synaptosome preparation (see fig. S4A; Fig.4C, D; table S5). After applying a bioinformatic filter to focus on excitatory neuronal transcripts (*9*, *10*), we identified 1,591 transcripts that exhibited increased translation at synapses compared with the non-synaptic cytosol. Prominent among synapse-enriched transcripts were those encoding synaptic vesicle proteins (e.g., *Syp*, *Syt1*), AMPA receptor auxiliary subunits (e.g., *Cnih2*, *Cacng2*, *Cacng3*, *Cacng8*, *Gsg1l*), pre- and postsynaptic adhesion molecules (e.g., *Nrxn1*, *Nlgn1)*, inhibitory and excitatory neurotransmitter receptor subunits (e.g., *Gria1*, *Gria2*, *Grin1*, *Grin2b*, *Gabra1*, *Gabra2*), actin regulators involved in synaptic structural remodeling (e.g., *Enah*, *Apc2*, *Wasf3*), and components of the postsynaptic Golgi apparatus and spine apparatus (e.g., *Aplp2*, *Clstn1*, *Tgoln2*, *Zdhhc17*; Fig. 4C). In contrast, among the 5,102 transcripts with increased translation in the non-synaptic cytosol were those encoding RNA-binding proteins and translation factors (e.g., *Fmr1*, *Elavl4*, *Stau2*, *Hnrnpa2b1*, *Eef2, Eif3c*, *Eif4e, Eef1a2*), spliceosomal and nuclear pre-mRNA processing components (e.g., *Sf3b1*, *U2af2*), transcriptional machinery (e.g., *Polr2g*, *Tbp*), ribosome biogenesis factors (e.g., *Npm1*, *Bop1*), and microtubule-based transport machinery (e.g., *Kif3a*, *Dctn1*, *Dync1li1*, *Pafah1b1*, *Ndel1*; Fig. 4C). GO terms associated with the synaptic membrane were strongly enriched among synapse-enriched transcripts, whereas cytosol-enriched transcripts were associated with nuclear and post-transcriptional RNA regulatory processes (Fig. 4D; table S5). Comparing our synaptic translatome with published CA1 somata and neuropil (comprising axons, dendrites and synapses) translatomes (*10*) revealed a functional partitioning of translation across neuronal subcompartments (Fig. 4E-G). We identified GOCC terms shared across synapse-, neuropil-, and somata-enriched transcripts and ranked them by enrichment score (mean log_2_ fold enrichment) to visualize spatial patterns of subcellular translation (table S6). The top 15 terms were all maximally enriched at synapses, with the neuropil typically showing greater enrichment than somata, indicating a closer functional alignment between synaptic and neuropil translation (Fig. 4E). Are there transcripts that exhibit differences in relative translation at synapses compared to the broader neuropil? To approach this question, we compared the rank-order abundance of translated mRNAs between compartments, focusing on the synapse-enriched transcripts identified in the synapse versus cytosol comparison. A subset of synapse-enriched transcripts exhibited pronounced upward shifts in rank from neuropil to synapse (‘rank-increased transcripts’; Fig. 4F), suggesting candidates for enhanced localized translation within synaptic compartments, including *Bdnf* (*39*), *Cacng3*, and *Slc17a6* (*40*). GO terms associated with rank-increased transcripts were ‘neuron to neuron synapse’, ‘synaptic membrane’, and ‘receptor complex’ among others (Fig. 4G; table S6).

While the above data identify the population of mRNAs that are translated within synapses, the observation that only a subset of synaptic ribosomes interact with the glutamate receptor suggests that there could be spatially segregated channels that coordinate the translation of distinct mRNAs within individual spines. To determine whether GluA1-associated ribosomes translate a distinct subset of synaptic mRNAs, we compared the two translatomes (RPLP0-associated vs. GluA1-associated; table S6). After background subtraction (fig. S7G; table S7), we found that 239 transcripts exhibited significantly enhanced translation by GluA1-associated ribosomes relative to the total synaptic ribosome pool (Fig. 4H; table S7). These GluA1-ribosome preferring transcripts included, for example, *Shank1*, *Camk2a*, and *Eef2* (Fig. 4I). On the other hand, 6,249 transcripts were more translated by the total synaptic ribosome pool than the GluA1-associated ribosomes. A GO analysis revealed that the GluA1-associated enhanced translation mRNAs were enriched for functions associated with the postsynaptic density, synapses, and the actin cytoskeleton (Fig. 4J,K; fig. S8E,F; table S7). The total (RPLP0-associated) synaptic translatome was also enriched for synaptic transcripts but also included mRNAs encoding vesicular and membrane proteins. Thus, GluA1-associated ribosomes may supply PSD scaffolding proteins (e.g., PSD95, GKAP, Shank1-3) and actin-scaffold adapters (e.g., Neurabin, Spinophilin, Protein 4.1N, Cortactin-binding protein 2) necessary for synaptic maintenance and remodeling during synaptic plasticity.

### CaMKIIα supports ribosome-PSD complexes

Our proximity ligation-mass spectrometry experiments (Fig. 1), co-IP (Fig. 2) and imaging experiments (Fig. 3) identified GluA1 as a synaptic ribosome interactor, but do not address whether the interactions are direct or mediated by intermediary protein factors. To obtain better resolution on the protein-protein interactions (PPIs) that govern these interactions we conducted cross-linking mass spectrometry (XL-MS; see Methods) on ribosome-enriched samples prepared from cortical synaptosomes (Fig. 5A). Overall, we identified 13,419 inter-protein residue pairs (crosslinks) and 2,241 PPIs among 836 proteins, including 73 ribosomal proteins (fig. S9A; table S8). After manually resolving crosslink assignments involving shared peptides (see Methods), analysis of the core synaptic ribosome interactome revealed 27 first-tier interacting partners, forming a subnetwork with 782 inter-protein crosslinks and 214 PPIs among 100 total proteins (Fig. 5B; table S8). As expected, the majority (66%) of inter-protein crosslinks mapped within the ribosomal complex. We focused on the postsynaptic density proteins that had crosslinks with ribosomal proteins. As observed previously (*41*) and also in this study, synaptic GluA1 crosslinks were largely confined to other members of the AMPA receptor complex and auxiliary subunits. In addition, we detected crosslinks from proteins that are closely associated with the postsynaptic density, including LIN7B and CaMKIIα. LIN7B crosslinked to RPL4 (Fig. 5B). CaMKIIα crosslinked to RPL35, RPL19 and the ribosome-associated elongation factor EEF1A2, which itself crosslinked to RPS24, RPL9 and RPL12 (Fig. 5B). We note that CaMKIIα was also identified as a synaptic ribosome-associated protein in our PL-MS and IP-MS experiments (Fig. 1D, 2C). The identified CaMKIIα crosslinks to RPL35 and RPL19 mapped to solvent-exposed sites on the 80S ribosome (Movie S1). Site-specific crosslinking data revealed multiple interaction sites within the kinase domain and near the regulatory segment of CaMKIIα – regions that mediate key interactions with most other postsynaptic partners (*42*) (Fig. 5C; fig. S9B). To depict the CaMKIIα-ribosome interaction visually, we performed crosslink-guided PPI docking using RPL35 or RPL19 with the CaMKIIα holoenzyme in its most extended (activation-competent) conformation (Protein Data Bank (PDB) 5U6Y). The 80S ribosome structure was superposed onto the predicted docking models of CaMKIIα bound to RPL35 or RPL19. The RPL35-based model positioned CaMKIIα at the ribosomal surface whereas the RPL19-based model exhibited apparent steric clashes with the 80S ribosome (Fig. 5D; fig. S9C), suggesting that the interaction is more structurally compatible with the RPL35 interface. The Cα-Cα distances of these crosslinks were evaluated on the proposed model. Among all 7 crosslinks between CaMKIIα and RPL35, five fell within the maximum 35 Å span of the crosslinker (Fig. 5D). The remaining 2 crosslinks were also consistent with the distance constraint when the flexibility of a disordered region in CaMKIIα is considered, which corresponds to approximately 30 Å between its most compact and most extended conformations (*43*). CaMKIIα and the AMPA receptor are known to be tightly associated within the PSD (*44*, *45*) where CaMKIIα phosphorylation of GluA1 regulates its conductance (Ser831; (*46*)). In our XL-MS network, we identified crosslinks between CaMKIIα and PSD95. PSD95, in turn, crosslinked to AMPA receptor subunit GluA3 and auxiliary subunits stargazin and TARP γ-3. Taken together, these data suggest the synaptic CaMKIIα-ribosome interaction positions the ribosome near the AMPA receptor (Fig. 1D).

**Fig. 5.**
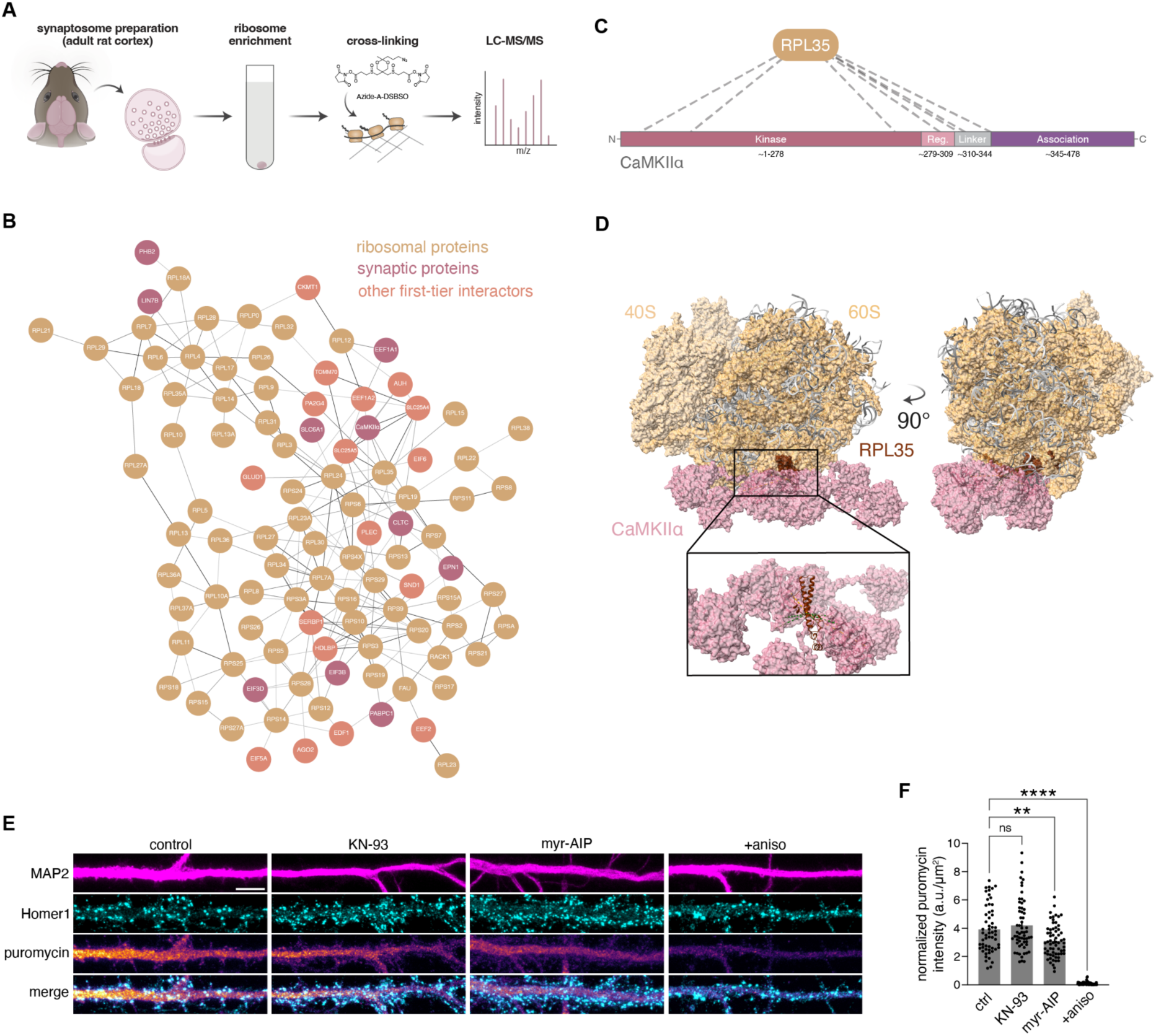
Synaptic ribosomes interact with CaMKIIα. (**A**) Schematic of XL-MS workflow. (**B**) XL-based interactome of synaptic ribosomes. Synaptic annotation is based on SynGO (dataset version 20231201). (**C**) Protein domain-level crosslink map between RPL35 and CaMKIIα. Domain classification based on (*72*). (**D**) Structural model of CaMKIIα binding to the ribosome. Most CaMKIIα–RPL35 crosslinks satisfy distance constraints (green), with remaining links accommodated by CaMKIIα linker flexibility (yellow). (**E**) Detection of newly-synthesized proteins in dendrites and synapses from control (untreated) neurons or neurons treated with 10 μM KN-93 or myr-AIP for 30 min. Nascent proteins were labeled with puromycin (5 min) in the absence or presence of the protein synthesis inhibitor anisomycin (see Methods). Scale bar, 5 µm. (**F**) Quantification of puromycin signal in postsynaptic regions of the apical dendritic arbor from control and treated neurons. Error bars show mean ± SEM. **p<0.01; ****p<0.0001, Brown-Forsythe and Welch’s ANOVA, Dunnett’s multiple comparisons test; n=58-64 neurons from 3 independent cultures.

We next tested the contribution of this “CaMKIIα-ribosome translation channel” to the total nascent protein synthesis detected within synapses. To do so, we metabolically labelled nascent proteins using a brief (5 min) pulse of puromycin (*47*) followed by detection with an anti-puromycin antibody (Fig. 5E,F). We compared nascent protein levels detected at synapses under control conditions or following treatment with two mechanistically distinct CaMKII inhibitors: the calmodulin-competitive inhibitor KN-93 and the substrate-binding site inhibitor myr-AIP (*48*, *49*) (see also (*50*)). Protein synthesis in postsynaptic compartments was selectively impaired by myr-AIP, but not by KN-93, suggesting that autonomously active CaMKIIα –which can be stabilized at the PSD (*51*, *52*) and is resistant to inhibition by KN-93 (*53*)– is sufficient to support postsynaptic protein synthesis (Fig. 5F). Together, these findings support a model in which both the structural interactions and enzymatic activity of CaMKIIα contribute to ribosome positioning and protein synthesis at synapses, potentially enabling CaMKIIα to function as a molecular scaffold linking the synaptic activity transmitted by GluAs to signaling and local translation.

### ER sequestration of endogenous GluA1 impairs synaptic *Camk2a* translation

To assess the importance of this GluA1-CaMKIIα synaptic complex in driving the translation of synaptic targets, we took advantage of an intrabody-based tool (*54*) that sequesters GluA1 at the endoplasmic reticulum (Fig. 6A; GluA1-KDEL) and thus prevents the coupling of GluA1 to synaptic ribosomes. We transfected primary hippocampal neurons with the GluA1-KDEL intrabody and, as expected, observed a depletion of surface GluA1 in dendrites; the control intrabody (an ER-retained GFP nanobody) had no effect on surface GluA1 expression (Fig. 6B,C). We assessed the effect of disrupting the synaptic GluA1-coupled ribosome on the translation of a GluA1-ribosome mRNA target (see Fig.4; *Camk2a*) and, as a control, a non-enriched mRNA target (*Dbn1*). We used Puro-PLA (*55*) to detect nascent CaMKIIα or Drebrin. We found that the GluA1-KDEL expressing dendrites exhibited significantly less nascent CaMKIIα (Fig. 6D,E) whereas Drebrin translation was unaffected (fig. S10B,C). Pretreatment with anisomycin significantly reduced the nascent signal for both proteins (Fig. 6F, fig. S10B,C). Taken together, these data suggest that proper synaptic localization of the AMPA receptor is important for the translation of select synaptic mRNAs.

**Fig. 6.**
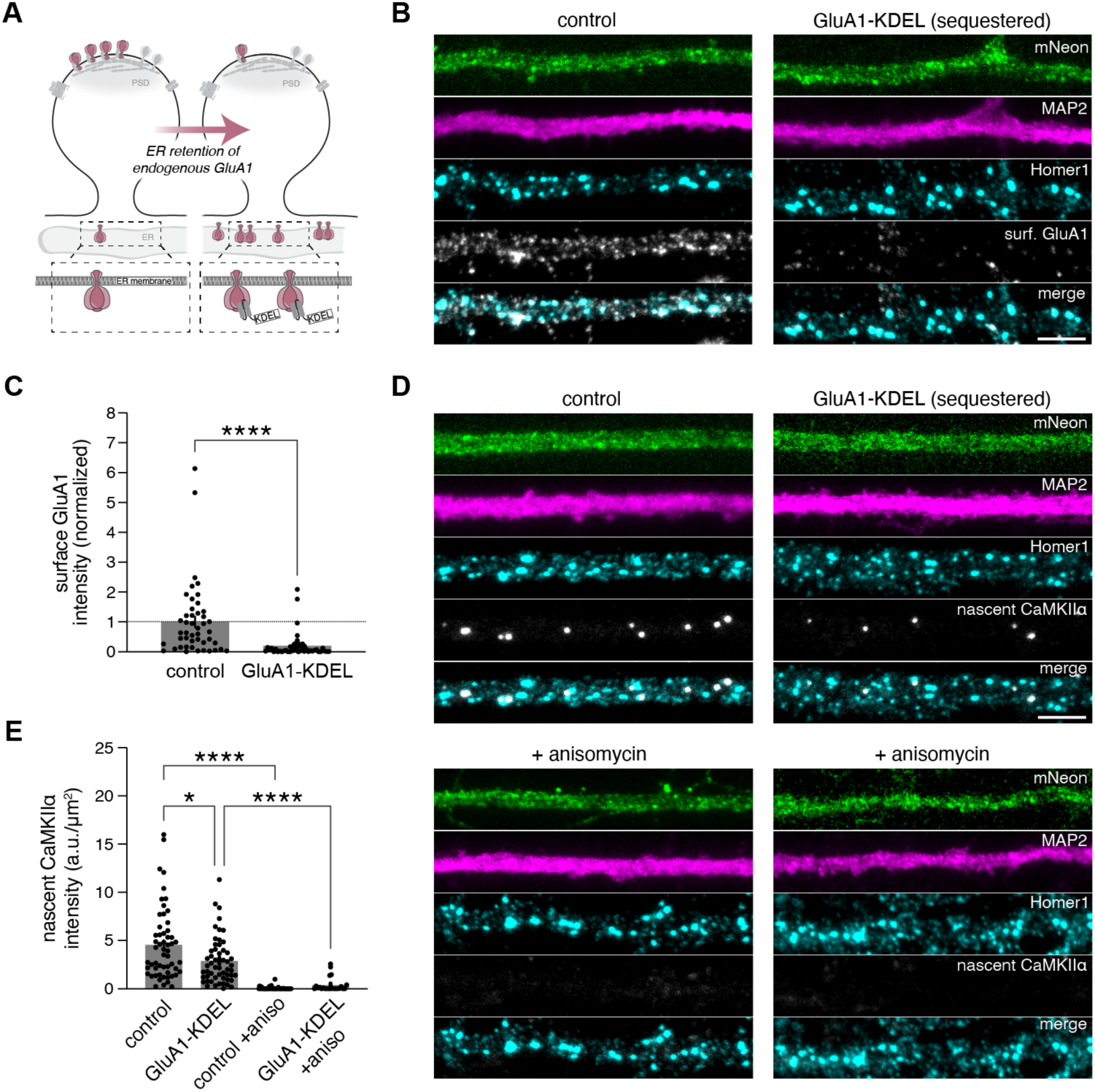
Decreased surface expression of endogenous GluA1 reduces translation of *Camk2a* mRNA within dendrites and synapses. (**A**) Approach for manipulating endogenous GluA1 surface levels (as developed by Kareemo et al. (*54*)). GluA1 surface trafficking was prevented by expressing, for 3.5 days, an ER-retained intrabody (via a C-terminal KDEL motif) that binds GluA1 and sequesters it in the ER. (**B**) Images of primary hippocampal dendrites expressing mNeon-tagged, ER-retained versions of either the GluA1 intrabody (GluA1-KDEL) or a control nanobody directed against a GFP epitope, and immunolabeled for surface GluA1, Homer1, and MAP2. Scale bar, 10 µm. (**C**) Quantification of GluA1 surface signal in control and GluA1-KDEL-expressing neurons. Error bars show mean ± SEM. ****p<0.0001, two-tailed unpaired t-test; n=45-46 neurons from 2 independent experiments. (**D**) Detection of nascent CaMKIIα in dendrites from control and GluA1-KDEL expressing neurons. Nascent CaMKIIα was labeled with puromycin (5 min) in the absence or presence of the protein synthesis inhibitor anisomycin (see Methods). Scale bar, 10 µm. (**E**) Quantification of nascent CaMKIIα in synaptic regions from a 50-80 μm dendritic segment from control and GluA1-KDEL expressing neurons. Error bars show mean ± SEM. *p<0.05, ****p<0.0001, Brown-Forsythe and Welch’s ANOVA, Dunnett’s multiple comparisons test; n=23-56 dendritic segments from 12-56 neurons from 2-3 independent cultures.

## Discussion

Using a subcellularly-targeted proximity labelling approach we identified the interactome of cortical synaptic ribosomes and found that a subpopulation of ribosomes is in close proximity to the AMPA-type glutamate receptor. The subcellular complex comprising ribosomes and AMPA receptors was further verified with co-IP experiments from cortical synaptosomes and visualized directly in dendritic spines using super-resolution imaging.

Many studies have identified abundant populations of mRNAs in axons, dendrites and synapses (*7*, *8*, *10–12*, *56*). The question arises as to whether these mRNAs are translated into protein (the “translatome”). In developing axons, the translatome has been identified with metabolic labeling (*57*) or by RNA-seq of mRNAs associated with ribosomes (*58*). Studies using GFP-tagged ribosomes have also identified, via pull-down and subsequent RNA-seq, populations of developing synaptic mRNAs associated with ribosomes in dopaminergic synaptosomes (*59*), and cortical excitatory synapses onto PV^+^ interneurons (*37*). Ribosome profiling of neuropil from microdissected brain slices (*9*, *10*) or proximity-labeled dendrites in culture (*60*) have furthermore identified local “translatomes”. What has been missing, however, is the identity of mRNAs that are specifically translated (with codon level resolution) within synapses themselves. Here we identified the synaptic translatome and detected the active translation of a large majority of the synaptically localized mRNA population. As expected, there was a strong enrichment of synaptic terms, along with membrane-associated categories, including ‘postsynaptic membrane’. A comparison with the neuropil (which includes processes and synapses) and somata translatomes (*10*) revealed the translation of synaptic cytosolic and membrane molecules becomes progressively enriched from the soma to the synapse.

XL-MS identified CaMKIIα as a likely physical mediator of the AMPA receptor-ribosome complex. It is known that, when activated, CaMKIIα translocates to postsynaptic sites (*51*, *61*) in close proximity to AMPA receptors (*45*, *62*) to potentiate synaptic strength through phosphorylation of PSD substrates including GluA1 itself (*46*) as well as an auxiliary subunit (*44*, *63*). Beyond its catalytic function, the structural roles of CaMKIIα are increasingly recognized as critical for synapse function, including the induction and early phase of long-term potentiation (*64*, *65*). Integrating XL-MS constraints with AlphaFold3-guided docking positioned activation-competent CaMKIIα adjacent to the ribosomal large subunit, in proximity to the ribosome exit tunnel. Such placement would position CaMKIIα near emerging nascent polypeptides, consistent with a potential role in co-translational regulation, including co-translational phosphorylation as described for Akt by mTORC2 (*66*). CaMKIIα has been reported to serve as a multivalent docking platform for synaptic regulators, including the proteasome (*67*). Our data indicate that CaMKIIα scaffolds not only degradative machinery, but also translational machinery.

Our data indicate that a subpopulation of ribosomes within dendritic spines are present in a complex with AMPA receptors, optimally positioning the ribosome to respond to synaptic signals, as speculated by early EM studies (*14*). Two previous studies in the developing nervous system identified adhesion molecule receptors (DCC and others) as mRNA-dependent interactors of ribosomes in the axonal growth cone (*23*, *24*); the ribosomes were found to dissociate from these anchoring molecules before translation. Here we show that the interaction of ribosomes with GluA1 is mRNA-independent and associated with active translation. The association of translating ribosomes with the AMPA receptor complex prompted us to ask whether proximity to the postsynaptic density biases translation toward a particular “privileged” subset of localized mRNAs, such as those encoding proteins involved in PSD reorganization. We found that transcripts preferentially translated by GluA1-associated ribosomes (the ‘subsynaptic’ translatome) included multiple PSD scaffolding proteins and actin-scaffold adapters (Fig. 4K). Single-spine two-photon glutamate uncaging studies have demonstrated increases in the number of PSD scaffolding proteins ∼1 hour after induction of structural plasticity (*68*), in a protein-synthesis dependent manner (*69*). Thus, GluA1-associated ribosomes may locally supply proteins required for the rapid expansion and stabilization of the PSD. More broadly, the close association of ribosomes with the AMPA receptor and associated PSD proteins places the translation machinery in an ideal position to decode and respond to synaptic signals. Indeed, the physical arrangement of the ribosome in close proximity to the site of excitatory transmission, arranged in nanodomains (*32–34*, *70*), may constitute a self-regulating-reinforcing loop where neurotransmitter signaling could directly modify the proteomic composition of the receptor signaling complex itself. It indicates that even within the small compartment of the synapse, there are even smaller subcompartments where decoding can occur.

## Acknowledgments

We thank Matthew Kennedy (Department of Pharmacology, University of Colorado Anschutz Medical Campus) for the GluA1 scFv and GFP nanobody mNeon-KDEL constructs; Susanne tom Dieck, Belquis Nassim-Assir and Ina Bartnik for assistance with experiments; Ina Bartnik, Nicole Fürst, Anja Staab and Christina Thum for preparation of primary rat neurons; Stephan Junek and Christine Bohnstaedt of the MPIBR imaging facility for technical support; Naomi Greenberg for help with characterizing APEX constructs; Claudia Fusco, Marcel Jüngling and the rest of the Schuman lab for discussions.

## Funding

This work was funded by the Max Planck Society (EMS), the European Union (ERC Advanced Investigator Awards NeuroRibo 743216 and DiverseSynapse 101054512 to EMS), and an EMBO Postdoctoral Fellowship (ALTF 238-2021, to AMB). Views and opinions expressed are however those of the authors only and do not necessarily reflect those of the European Union or the European Research Council Executive Agency. Neither the European Union nor the granting authority can be held responsible for them.

Max Planck Society (EMS)

European Union Advanced Investigator Award NeuroRibo 743216 (EMS)

European Union Advanced Investigator Award DiverseSynapse 101054512 (EMS)

European Molecular Biology Organization Postdoctoral Fellowship ALTF 238-2021 (AMB)

## Author contributions

Conceptualization: AMB, EMS

Methodology: AMB, GT, MW, KD, JDL, FL, EMS

Investigation: AMB, MM, MW, KD, SGM, AS, EC

Formal Analysis: AMB, MM, GT, MW, KD, SGM

Visualization: AMB, MM, GT, MW, KD

Software: AMB, GT, KD

Funding acquisition: AMB, EMS

Project administration: AMS, EMS

Supervision: JDL, FL, EMS

Writing – original draft: AMB, EMS

Writing – review & editing: AMB, MM, GT, MW, KD, SGM, AS, EC, JDL, FL, EMS

## Data, code, and materials availability

The raw ribosome profiling and RNA sequencing data will be available in the NCBI Sequence Read Archive (SRA) under BioProject PRJNA1381615 upon publication. The PL-MS, IP-MS, and XL-MS proteomics data have been deposited to the ProteomeXchange Consortium via the PRIDE partner repository (*71*) under dataset identifiers PXD072116 (PL-MS and IP-MS) and PXD072664 (XL-MS) and will be released upon publication. All customized scripts will be available via the Edmond Data Repository of the Max Planck Society (doi:10.17617/3.OMEAYP) upon publication.

## Materials and Methods

### Animals

Timed-pregnant Sprague Dawley female rats (typically at gestation day 16) were obtained from Charles River Laboratories and housed under a 12-h light/dark cycle with ad libitum access to food and water until the litter was born. Primary neurons were derived from both male and female neonatal pups, and brains/cortical tissue were obtained from adult female rats (aged 11-16 weeks) for acute slice experiments/synaptosome preparation. The housing and sacrifice procedures involving animal treatment and care were conducted in conformity with the institutional guidelines that are in compliance with national and international laws and policies (Directive 2010/63/EU; German animal welfare law; Federation of European Laboratory Animal Science Associations guidelines). The animals were euthanized according to annex 2 of § 2 Abs. 2 Tierschutz-Versuchstier-Verordnung. Animal numbers were reported to the local authority (Regierungspräsidium Darmstadt).

### Preparation of primary cultured neurons

Primary rat hippocampal and cortical neurons were prepared and maintained as described previously (*73*). Briefly, hippocampi and cortices were dissected from postnatal day 0-2 rat pups of either sex, dissociated with papain (Sigma, #P3125), plated, and maintained up to 22 days in vitro (DIV) at 37°C in 5% CO_2_. Dissociated hippocampal neurons were plated at a density of 1.3 × 10^4^ cells/cm^2^ for STED imaging or 2.0 × 10^4^ cells/cm^2^ for all other imaging experiments onto the 14-mm microwell of 35-mm poly-D-lysine-coated glass-bottom dishes (MatTek, #P35G-1.5-14-C). For biochemical experiments, dissociated cortical neurons were plated at a density of 5.1 × 10^4^ cells/cm^2^ onto poly-D-lysine-coated 10-cm dishes. Glia-conditioned medium was added 3 h after plating and again on DIV 4. Fresh growth medium (Neurobasal-A, Gibco, #10888-022) supplemented with B27 (Gibco, #17504-044) and GlutaMAX (Gibco, #035050-038) was added on DIV 11 and weekly thereafter.

### Molecular cloning

APEX constructs were cloned into a pAAV hSyn1 backbone derived from pAAV hSyn1-mRuby2 (Addgene plasmid #99126, a gift from Viviana Gradinaru) by Gibson Assembly (NEB, E5510). For the APEX-SYN construct (pAAV hSyn1-FLAG-APEX2-Homer1c), FLAG-APEX2 was PCR-amplified from pcDNA3 FLAG-APEX2-NES (Addgene #49386; gift from Alice Ting), Homer1c was PCR-amplified from CRY2-GFP-Homer1c (Addgene #89442; gift from Matthew Kennedy), and a SAGSAG linker was inserted between the two. For the APEX-CYTO construct (pAAV hSyn1-FLAG-APEX2-NES), FLAG-APEX2-NES was PCR-amplified from pcDNA3 FLAG-APEX2-NES. For the APEX-NUC construct (pAAV hSyn1-FLAG-APEX2-SENP2), APEX2-SENP2 was PCR-amplified from V5-APEX2-SENP2 (Addgene #129276; gift from Alice Ting). All constructs were verified by whole-plasmid sequencing (Eurofins).

### AAV production

Recombinant AAV-DJ vectors encoding APEX constructs were packaged in HEK293T cells by calcium phosphate transfection with the AAV expression plasmid (encoding the APEX construct), pHelper, and pDJ (Cell Biolabs, #VPK-410-DJ). Three days post-transfection, cells were harvested, lysed, and viral particles were purified by iodixanol discontinuous gradient ultracentrifugation for 2 hr 25 min at 350,000×g at 4°C using a Type 70 Ti rotor, thickwall polycarbonate bottles (Laborgeräte Beranek, #4415), and adapters (Laborgeräte Beranek, #11035). The 40% iodixanol fraction containing AAV was collected, concentrated, aliquoted, and stored at -80°C until use. The amount of AAV used for each construct was determined empirically to achieve 80-90% transduction of primary cultured neurons at 3 days post-infection.

### APEX proximity labeling in primary rat hippocampal neurons - imaging

For APEX imaging experiments, DIV 18-19 hippocampal neurons were transduced with an AAV encoding either hSyn-FLAG-APEX2-SENP2, hSyn-FLAG-APEX2-NES or hSyn-FLAG-APEX2-Homer1c. After 3 d of expression (DIV 21-22), cells were preincubated for 30 min at 37°C with 500 μM biotin-phenol (Sigma, #SML2135) in conditioned medium. APEX proximity labeling was then initiated by adding H_2_O_2_ to a final concentration of 1 mM and gently agitating for 1 min. The reaction was quenched by adding 2× quench buffer (20 mM sodium azide, 20 mM sodium ascorbate, 10 mM Trolox in aCSF [130 mM NaCl, 5 mM KCl, 10 mM HEPES, 30 mM glucose, 1 mM MgCl_2_, 2 mM CaCl_2_; pH 7.4]) followed immediately by three washes in 1× quench buffer. Quench buffers were pre-warmed to 37°C. Cells were then fixed for 10-12 min with a HEPES-buffered 4% PFA-sucrose (4% PFA, 4% sucrose, 50 mM HEPES in PBS, pH 7.4) and permeabilized with 0.1% Triton X-100 in PBS for 10-12 min at RT. After blocking with 4% goat serum in PBS, pH 7.4 for 1 hr at RT, cells were incubated with primary antibodies in blocking buffer for 2 hr at RT, followed by secondary antibodies in blocking buffer for 1 hr at RT. Cells were mounted in ProLong Gold Antifade Mountant with DAPI (Thermo Scientific) or Aqua-Poly/Mount (Polysciences, #18606) and overlaid with a 13-mm glass coverslip. Three PBS washes were performed between all steps post-fixation, except between blocking and primary antibody incubation. Primary antibodies: mouse IgG1 anti-FLAG (1:200; Sigma, #F1804); mouse IgG2a anti-PSD95 (1:300; GeneTex, #GTX22723); guinea-pig anti-MAP2 (1:1000; Synaptic Systems, #188004). Secondary antibodies and probes: Alexa Fluor 568 goat anti-mouse IgG1 (1:500; Invitrogen, #A21124); Alexa Fluor 647 goat anti-mouse IgG2a (1:500, Invitrogen, #A21241); CF 405S donkey anti-guinea pig (1:500, Sigma, #SAB4600230); Alexa Fluor 488-conjugated streptavidin (1:500; Invitrogen, #S11223). Streptavidin was included in the same incubation as the secondary antibodies. Controls included: (i) non-transduced neurons (no APEX control), and (ii) neurons expressing each APEX construct where either biotin-phenol or H_2_O_2_ was omitted.

### Image acquisition and analysis for APEX proximity labeling

Samples were imaged on a Zeiss LSM880 confocal microscope equipped with a Plan-Apochromat 63×/1.4 NA Oil objective and ZEN 2.3 SP1 FP3 (black edition) software (Zeiss; v14.0.27.201). APEX-positive neurons, identified by anti-FLAG immunostaining, were selected for imaging and quantification if they had a pyramidal-shaped soma, a prominent apical dendrite, visible dendritic spines, and a dendritic arbor that could be unambiguously attributed to a single neuron. The z-stack was set to encompass the entire volume of a neuron, with optical slice thickness set to optimal. Laser power and detector gain were adjusted to avoid pixel saturation, and image acquisition settings were held constant across conditions (i.e., for all APEX constructs). Quantification was performed on maximum-intensity projections of the raw z-stack images in ImageJ2 (*74*). A 10-20 μm dendritic segment was selected as the ROI and linearized. For each ROI, two masks were generated: a ‘synaptic mask’ defined by the PSD95 signal, and a mask encompassing both synaptic and non-synaptic regions derived from the combined MAP2 and PSD95 signal. The background-subtracted mean fluorescence intensity of the streptavidin signal within each mask was quantified. Non-synaptic mean fluorescence intensity was obtained by subtracting the background-subtracted mean fluorescence intensity within the synaptic mask from that within the combined (synaptic + non-synaptic) mask. Images were expanded and smoothed in ImageJ2 for display only.

### APEX proximity labeling in primary rat cortical neurons - biochemistry/proteomics

For APEX biochemistry/proteomics experiments, DIV 14 cortical neurons in 10-cm dishes were transduced with an AAV encoding either hSyn-FLAG-APEX2-SENP2, hSyn-FLAG-APEX2-NES or hSyn-FLAG-APEX2-Homer1c. Nontransduced (no APEX) neurons served as a control. After 3 d of expression (DIV 17), cells were preincubated for 30 min at 37°C with 500 μM biotin-phenol (Sigma, #SML2135) in conditioned medium. APEX proximity labeling was then initiated by adding H_2_O_2_ to a final concentration of 1 mM and gently agitating for 1 min. The reaction was quenched by adding RT 2× quench buffer followed immediately by two washes in ice-cold 1× quench buffer supplemented with 100 μg/mL cycloheximide (CHX; Sigma, #C7698). Cells were scraped in 100 μL of ice-cold 2× ribosome lysis buffer (40 mM Tris pH 7.4, 300 mM NaCl, 10 mM MgCl_2_, 2% Triton X-100, 200 μg/mL CHX, 2 mM dithiothreitol (DTT; Sigma, #43816), 48 U/mL TurboDNase, 400 U/mL RNasin Plus RNase inhibitor, and 2× cOmplete EDTA-free protease inhibitor cocktail). Lysates from 3 10-cm dishes were pooled, manually triturated through a 0.4 × 20 mm needle (15 passes), and centrifuged at 10,000×g for 10 min at 4°C. An aliquot (∼1%) of the resulting supernatant was reserved as the lysate for Western blot and proteomics. A volume corresponding to 1 A260 unit of the remaining sample was layered onto a 1-mL sucrose cushion (34% sucrose, 20 mM Tris pH 7.4, 150 mM NaCl, 5 mM MgCl_2_, 100 μg/mL CHX, 1 mM DTT) in a thickwall polycarbonate tube (Beckman, #349622) and centrifuged for 30 min at 4°C at 55,000 rpm (367,600×g) with a SW55Ti rotor (acceleration: max, deceleration: 7). The ribosome pellet was resuspended in 20 μL of ice-cold RIPA lysis buffer (50 mM Tris pH 8.0, 150 mM NaCl, 1% Triton X-100, 0.5% sodium deoxycholate, 0.1% SDS) supplemented with 1× protease inhibitor cocktail (Sigma, #P8849) and 1 mM PMSF. Aliquots were reserved for Western blot (5%) and proteomics (5%). The remaining cushion sample was incubated overnight at 4°C with 10 μL streptavidin magnetic beads (Pierce, #88817) pre-equilibrated in RIPA lysis buffer, with rotation. Beads were then washed by briefly vortexing in a series of ice-cold buffers (200 μL for each wash) to remove nonspecific binders: twice with RIPA lysis buffer, once with 1 M KCl, once with 0.1 M Na_2_CO_3_, once with 2 M urea in 50 mM Tris-HCl pH 8.0, twice with RIPA lysis buffer, twice with a modified RIPA lysis buffer (50 mM Tris pH 8.0, 150 mM NaCl, 0.2% sodium deoxycholate, 0.1% SDS) and twice with 150 mM NaCl in 50 mM Tris pH 8.0. Biotinylated proteins were eluted from beads by heating at 95°C in 30 μL of 3× NuPAGE LDS sample buffer supplemented with 20 mM DTT and 2 mM biotin for 10 min with shaking at 400 rpm. Aliquots of the eluate were reserved for Western blot (33%) and proteomics (67%). Samples were kept on ice throughout the procedure.

### Cortical tissue collection and synaptosome preparation

To enrich postsynaptic ribosomes, we used a synaptosome preparation with abundant bipartite structures containing intact postsynapses (*37*). Cortices from both brain hemispheres (adult rats, 12-16 weeks) were dissected on ice and immediately homogenized in SynPER reagent (Thermo Scientific, #87793) supplemented with 1 mg/mL heparin (Sigma, #H3393), 100 μg/mL cycloheximide (CHX; Sigma, #C7698), 1 mM DTT (Sigma, #43816), 200 U/mL RNasin Plus RNase inhibitor (Promega, #N2611), and 1× cOmplete EDTA-free protease inhibitor cocktail (Roche) using a glass Dounce homogenizer. The homogenate was centrifuged twice in succession at 1200×g for 10 min at 4°C. The supernatant (S1) was then centrifuged at 15,000×g for 20 min at 4°C. The resulting pellet (P2) containing synaptosomes was resuspended in 1.25 mL of ribosome lysis buffer (10 mM Tris pH 7.4, 100 mM KCl, 5 mM MgCl_2_, 1% NP-40 (BioVision, #2111-100), 100 μg/mL CHX, 1 mM DTT, 24 U/mL TurboDNase, 200 U/mL RNasin Plus RNase inhibitor, and 1× cOmplete EDTA-free protease inhibitor cocktail). The synaptosome suspension was designated as the synaptic fraction, and the S2 supernatant as the cytosolic fraction (see fig. S4A for schematic). Synaptosomes were prepared from one rat, except for two Ribo-seq experiments in which the synaptosome suspension from two rats were pooled. Samples were kept on ice throughout the procedure.

### Enrichment of synaptic ribosomes by sucrose cushion

Synaptosome membranes were lysed by manual trituration through a 0.4 × 20 mm needle (20 passes) and then 0.3 A260 units of sample was layered onto a 1-mL sucrose cushion (34% sucrose, 20 mM Tris pH 7.4, 25 mM KCl, 5 mM MgCl_2_, 100 μg/mL CHX, 1 mM DTT) in a thickwall polycarbonate tube (Beckman, #349622) and centrifuged for 30 min at 4°C at 55,000 rpm (367,600×g) with a SW55Ti rotor (acceleration: max, deceleration: 7). The ribosome pellet was resuspended in 100 μL of ice-cold ribosome lysis buffer (10 mM Tris pH 7.4, 100 mM KCl, 5 mM MgCl_2_, 1% NP-40, 100 μg/mL CHX, 1 mM DTT, 24 U/mL TurboDNase, 200 U/mL RNasin Plus RNase inhibitor, and 1× cOmplete EDTA-free protease inhibitor cocktail) and manually triturated through a 0.4 × 20 mm needle (10 passes). Samples were kept on ice throughout the procedure. For XL-MS samples, buffers contained 20 mM HEPES pH 7.5 instead of Tris, and the ribosome pellet was briefly washed in HEPES-based lysis buffer prior to resuspension in 50 μL.

### Endogenous co-immunoprecipitation

For each IP, 50 μg of input sample was pre-cleared with 5 μL Dynabeads Protein G (Invitrogen) for 1 h at 4°C with rotation. The pre-cleared sample was then incubated for 2 h at 4°C with 5 μL Dynabeads Protein G that had been pre-coupled overnight at 4°C to 1 μg of the following antibodies: mouse IgG2aκ anti-RPLP0 [9D5] (MBL, #RN004M), rabbit anti-GluA1 (C-terminal; Synaptic Systems, #182003), mouse IgG2a κ isotype control [6H3] (MBL, #M076-3), or rabbit IgG isotype control (Merck, #12-370). Note that the anti-GluA1 antibody recognizes the C-terminus of GluA1 (amino acids 829-907). Beads were washed four times in ice-cold high-salt buffer (20 mM Tris pH 7.4, 200 mM KCl, 10 mM MgCl_2_, 0.01% NP-40, 100 μg/mL CHX, 1 mM DTT) and bound proteins were eluted in 30 μL of 2× NuPAGE LDS sample buffer supplemented with NuPAGE reducing agent by heating at 95°C for 10 min with shaking at 400 rpm. In experiments designed to induce ribosome subunit dissociation, bead washes were performed in high-salt buffer without MgCl₂. For ribosome profiling experiments, pre-cleared samples were digested with RNase I (0.1 U/μg RNA, LGC, #N6901K) for 45 min at RT with rotation, and then immediately cooled and treated with 10 μL SUPERase·In RNase inhibitor (Thermo Scientific, #AM2694) prior to IP. Samples were eluted in 30 μL RLT Plus buffer (RNAeasy Plus Micro Kit, Qiagen) supplemented with 1% β-mercaptoethanol, vortexed for 30 sec, incubated at RT for 5 min, and the supernatant was stored at -80°C until RNA extraction.

### SDS-PAGE and Western blot analysis

Denatured protein extracts (prepared as described above) were separated by SDS-PAGE using Novex 4-12% Bis-Tris precast gels (Invitrogen) for 2 h at 120 V in NuPAGE MOPS SDS running buffer (Invitrogen, #NP0001) and then transferred onto PVDF membranes (Immobilon, #IPFL0001) for 1 h 35 min at 135 V with cooling (ice pack) in Novex Tris-Glycine Buffer (Invitrogen, #LC3675) with 10% methanol. For APEX experiments, 33% of the eluate, 5% of the cushion and ∼2-2.5 ug of lysate were loaded per gel lane. For IP experiments, 50% of the eluate and ∼0.25-1 ug of input (corresponding to 0.5–2% of the material used for the IP) was loaded per gel lane. Membranes were stained using Revert 700 Total Protein Stain (LI-COR, #926-110156) and destained before blocking in Intercept TBS Blocking Buffer (LI-COR, #927-60001) for 1 h at RT. Membranes were incubated overnight at 4°C with primary antibodies in 0.5× Intercept TBS Blocking Buffer supplemented with 0.2% Tween-20. For APEX experiments, membranes were blocked overnight at 4°C and incubated at RT with primary antibodies. Following three 5-min washes in TBS supplemented with 0.1% Tween-20 (0.1% TBS-T), membranes were incubated for 1 hr at 4°C with either near-infrared fluorescent secondary antibodies diluted 1:5000 in 0.5× Intercept TBS Blocking Buffer supplemented with 0.2% Tween-20 and 0.01% SDS, or HRP-conjugated secondary antibodies diluted 1:3000 in 5% non-fat dry milk (VWR, #84615.0500) in TBS. For fluorescence detection, membranes were washed three times (5 min each) in 0.1% TBS-T and kept in TBS before imaging using a LI-COR Odyssey 9210 imaging system and Image Studio Lite software (LI-COR, v.3.0.30). For chemiluminescence detection, membranes were washed five times (15-20 min each) in 0.5% TBS-T and three times in TBS before developing with SuperSignal West Femto Maximum Sensitivity Substrate (Thermo Scientific, #34094) and imaging using an Azure 280 imaging system. Primary antibodies: rabbit anti-CaMKII [EP1829Y] (1:1000; Abcam, #ab52476); mouse anti-CaMKII [6G9] (1:1000; Cell Signaling, #50049S); mouse anti-FLAG (1:1000; Sigma, #F3165); mouse anti-GAPDH [6C5] (1:2500; Abcam, #ab8245); rabbit anti-GluA1 (1:1000; MilliporeSigma, #ABN241); mouse anti-GluA2 [6C4] (1:1000; MilliporeSigma, #MAB397); rabbit anti-GluN2A (1:1000; MilliporeSigma, #AB1555P); mouse anti-GluN2B [S59] (1:1000; Abcam, #ab93610); rabbit anti-PABP (1:1000; Abcam, #ab21060); mouse anti-PSD95 [7E3-1B8] (1:1000; Thermo, #MA1-046); mouse IgG2a κ anti-RPLP0 [9D5] (1:1000; MBL, #RN004M); rabbit anti-RPL24 (1:1000; Proteintech, #17082-1-AP); rabbit anti-RPL26 (1:1000; Sigma, #R0655); rabbit anti-RPS3 (1:1000; Bethyl, #A303-840A); rabbit anti-RPS3A (1:1000; Bethyl, #A305-001A); rabbit anti-RPS11 (1:1000; Bethyl, #A303-936A); rabbit anti-SAP102 (1:1000; Synaptic Systems, #124213). Secondary antibodies and probes: IRDye 800CW streptavidin (1:5000; LI-COR, #926-32230); IRDye 680RD goat anti-rabbit IgG (1:5000; LI-COR, #926-68071); IRDye 800CW goat anti-mouse IgG (1:5000; LI-COR, #926-33210); HRP-conjugated goat anti-rabbit IgG (1:3000; Dako, #P0448); HRP-conjugated horse anti-mouse IgG (1:3000; Cell Signaling, #7076).

### MS sample preparation (PL-MS, IP-MS)

Proteins were prepared for bottom-up proteomics using a suspension trapping protocol as previously reported (*75*). For APEX-MS experiments, 67% of the eluate, 5% of the cushion and ∼5 µg of the lysate was digested (4.44 +/- 1.23 µg, mean +/- SD). For IP-MS, 50% of the eluate and ∼10 µg of lysate or cushion was prepared. In brief, the samples were mixed with lysis buffer (10% SDS, 100 mM Tris, pH 7.55, with H_3_PO_4_) in a 1:1 ratio, reduced using 20 mM DTT for 10 min at RT, and alkylated with 50 mM iodoacetamide (IAA) for 30 min at RT in the dark. Subsequently, the samples were acidified using phosphoric acid in a final concentration of ∼1.2%. Binding/wash buffer (BW buffer; 90% methanol, 50 mM Tris, pH 7.1 with H_3_PO_4_) was added in a 1:7 ratio. The protein suspension was loaded onto the S-trap filter (size: micro; ProtiFi) by centrifugation for 20 sec at 4000×g. Trapped proteins were washed with 150 μl of BW buffer four times. Trypsin (1 μg; Promega) was added in 60 μL of 40 mM ammonium bicarbonate (ABC) buffer. Digestion was performed overnight (∼18 hours) at RT in a humidified chamber. Peptides were collected by washing in three consecutive steps by centrifugation at 4000×g for 40 sec starting with digestion buffer and two washes of 0.2% formic acid in MS-grade water. After digestion, peptides were desalted using an adapted C18 StageTip protocol (*76*). StageTips were prepared by placing of two disks of C18 material (Empore 3M) into a 200 μL pipette tip (Eppendorf). Disks were conditioned using 100 μL pure methanol and centrifuged at 2000×g for ∼3 min. This was followed by a wash with 100 μL 50% acetonitrile (ACN) with 0.5% acetic acid and equilibration with two washes of 100 μL 0.5% acetic acid. Peptides were loaded and reloaded on the disks by centrifugation at 1500×g for ∼5 min (or until all solvent had passed) and desalted by washing with 100 μL 0.5% acetic acid twice at 2000×g for 3 min. Purified peptides were eluted in two steps, using 75 μL 50% ACN with 0.5% acetic acid and centrifugation at 2000×g for ∼3 min. Peptides were dried *in vacuo* at 45°C.

### LC-MS/MS data acquisition (PL-MS, IP-MS)

Dried peptides were reconstituted in 95% MS-grade H_2_O, 5% ACN, and 0.1% FA. For PL-MS experiments, peptides were loaded onto a C18-PepMap 100 trapping column (particle size 3 µm, L = 20 mm, ThermoFisher Scientific) and separated on a C18 analytical column with an integrated emitter (particle size = 1.7 µm, ID = 75 µm, L = 50 cm, CoAnn Technologies) using a nano-HPLC (Dionex U3000 RSLCnano) coupled to a nanoFlex source (2000 V, ThermoFisher Scientific). The temperature of the column oven (SonationAnalytics) was maintained at 55°C. Trapping was carried out for 6 min with a flow rate of 6 μL/min using a loading buffer (100% H_2_O, 2% ACN with 0.05% trifluoroacetic acid). Peptides were separated by a gradient of water (buffer A: 100% H2O and 0.1% FA) and acetonitrile (buffer B: 80% ACN, 20% H_2_O, and 0.1% FA) with a constant flow rate of 250 nL/min. In 155 min runs, peptides were eluted by a non-linear gradient with 120 min active gradient time, as selected and reported for the respective MS method by Muntel et al. (*77*). Analysis was carried out on a Fusion Lumos mass spectrometer (ThermoFisher Scientific) operated in positive polarity and data-independent acquisition (DIA) mode. In brief, the 40-window DIA method had the following settings: full scan: orbitrap resolution = 120k, AGC target = 125%, mass range = 350-1650 m/z, and maximum injection time = 100 ms. DIA scan: activation type: HCD, HCD collision energy = 27%, orbitrap resolution = 30k, AGC target = 2000%, maximum injection time = dynamic. For IP-MS experiments, peptides were analyzed using a nanoElute 1 nano-HPLC coupled to a timsTOF Pro II mass spectrometer via a captive spray ion source (1600 V, Bruker Daltonics). Peptides were loaded directly onto the analytical column (15 cm × 150 µm column with 1.5 μm C18-beads, PepSep), maintained at 60°C and connected to a 20 µm ZDV sprayer (Bruker Daltonics). In 25 min runs, peptides were separated in a linear gradient of water (buffer A: 100% Hs_2_O and 0.1% FA) and acetonitrile (buffer B: 100% ACN and 0.1% FA), ramping from 2% to 38% buffer B in active gradient time of 21 min with a constant flow rate of 800 nL/min. For data acquisition in DIA mode, the ‘short-gradient’ DIA-PASEF method was employed. In brief, 21 dia-PASEF windows were distributed to a TIMS scan each and designed to cover an m/z range from 475 to 1000 m/z in 25 Da windows, leading to an estimated method cycle time of 0.95 sec. The ion mobility range was set from 1.30 to 0.85 Vs/cm². Further details on the method parameters are embedded in the uploaded raw data.

### Analysis of DIA LC-MS/MS data (PL-MS, IP-MS)

DIA raw files were processed with the open-source software DIA-NN (IP-PL: v1.8.2 beta 27; IP-MS: v1.9, (*78*)) using a library-free approach. A spectral library was predicted using the *in silico* FASTA digest (Trypsin/P) option with the UniProtKB database (Proteome_ID: UP000002494) for *Rattus norvegicus,* including common MS contaminants. Deep learning-based spectra and RT prediction were enabled. The covered peptide length range was set to 7-35 amino acids and missed cleavages to 2. N-terminal methionine excision and methionine oxidation were set as variable modifications, including cysteine carbamidomethylation as a fixed modification. According to most of DIA-NN’s default settings, scan windows were set to 0; isotopologues and match-between-runs were enabled, while shared spectra were disabled. For orbitrap data, MS1 and MS2 mass accuracies were inferred automatically, while for timsTOF data mass accuracies were fixed at 15 ppm and the unrelated runs option was enabled. Protein inference was performed using genes with the heuristic protein inference option enabled. The neural network classifier was set to single-pass mode, and the quantification strategy was selected as ‘QuantUMS (high precision)’. The cross-run normalization was set to ‘RT-dependent’, the library generation to ‘smart profiling’, and the speed and RAM usage to ‘optimal results’. No normalization was applied (‘--no-norm’).

For IP-MS data, the DIA-NN report was processed using the DIA-NN R package (v1.0.1), filtering for proteotypic peptides and excluding contaminants (keratin). Protein roll-up was performed using the maxlfq-algorithm on the normalized precursor intensities with the following q-value thresholds: q-value <= 0.01, protein group q-value <= 0.01, and global protein group q-value <= 0.01. Protein LFQ intensity was log2-transformed. For PL-MS data, the DIA-NN report was processed in R using the msdap (*79*), QFeatures (*80*), and SummarizedExperiment (*81*) packages. Initial group-wise filtering across labeling conditions (no-APEX control, APEX-NUC, APEX-CYTO, APEX-SYN) was performed using msdap, requiring proteins to be supported by at least two peptides. For downstream analysis, peptides were filtered to retain those quantified in ≥4 replicates in at least one of the APEX baits (APEX-NUC, APEX-CYTO, APEX-SYN). Missing peptide intensities were imputed under a missing-at-random assumption using the msImpute R package (method v2; (*82*)), with the labeling condition specified as the grouping variable. Imputed peptide intensities were normalized using robust quantile normalization and subsequently aggregated to protein-level intensities using QFeatures.

### Differential expression analysis (PL-MS)

Differential protein abundance was modeled using msqrob2 (*83*) using a mixed-effects model including labeling condition as a fixed effect and replicate as a random intercept. Differential abundance was first assessed for each APEX bait versus control across all proteins; proteins significant in any bait-versus-control comparison were retained for subsequent bait-versus-bait contrasts. Multiple testing correction was controlled using the Benjamini-Hochberg procedure, with FDR < 0.05 considered significant.

### GO enrichment analysis (PL-MS)

GO enrichment analyses were performed using clusterProfiler (*84*), with internal significance filtering (pvalueCutoff = 0.01; qvalueCutoff = 0.01) and Benjamini-Hochberg multiple testing correction. Protein identifiers (UniProt accessions) were mapped to Entrez Gene IDs. For each condition-specific enrichment analysis, the background comprised all proteins retained in the differential expression model for the corresponding comparison. Redundant GO terms were removed using rrvgo (*85*), which applies GOSemSim semantic similarity (Rel metric, similarity threshold 0.3; (*86*)). For visualization, the top 10 non-redundant terms (ranked by adjusted p-value) were selected.

### GO enrichment analysis (IP-MS)

GO analysis and redundancy filtering were performed as described above using >3-fold enriched protein groups and gene-level exclusives as input. Proteins were mapped to gene identifiers prior to GO analysis. For GluA1 and Ribo IPs, proteins were filtered based on exclusivity to the IP or >3-fold enrichment relative to matched isotype controls. For each IP, proteins quantified in both replicates and in at least one matched isotype control replicate were included in the fold-enrichment analysis. Proteins quantified in the IP replicates but absent from the matched isotype control replicates were classified as IP-exclusive. All proteins quantified in the experiment were used as background. For visualization, the top 10 non-redundant terms (ranked by adjusted p-value) were selected.

GO analyses were performed for proteins in the synaptic fraction and synaptic fraction cushion (fig. S3C). Synaptic fraction cushion proteins were filtered based on exclusivity to the cushion or >3-fold enrichment relative to the synaptic fraction. Proteins quantified in both replicates of each fraction were included in fold-enrichment analyses. Proteins detected in both cushion replicates but absent from the synaptic fraction replicates were classified as cushion-exclusive. All proteins quantified across the homogenate, synaptic fraction, and synaptic fraction cushion were used as background. GO terms were not filtered for redundancy. For visualization, the top 10 terms (ranked by adjusted p-value) were selected.

Dot sizes reflect a normalized fold enrichment rank computed as 1 - ((rank - 1) / (max(rank) - 1)), enabling visual comparison of relative enrichment between datasets with differing numbers of significant terms.

GO analyses were performed for proteins in the Ribo IPs from the synaptic fraction and synaptic fraction cushion (fig. S3D). Proteins were filtered based on exclusivity to the IP or >3-fold enrichment relative to matched isotype controls, with all proteins quantified in the experiment used as background. GO terms were not filtered for redundancy. For visualization, the top 10 terms (ranked by adjusted p-value) were selected. Dot sizes reflect a normalized fold enrichment rank computed as 1 - ((rank - 1) / (max(rank) - 1)), enabling visual comparison of relative enrichment between datasets with differing numbers of significant terms.

### N-terminal bias analysis

The DIA-NN report was filtered to retain only proteotypic, non-contaminant peptides (as described above) observed in ≥3 eluate replicates per condition. For each leading protein ID, the corresponding full-length amino acid sequence was retrieved from UniProt, and peptides were mapped to their parent sequence. The relative position of each peptide was defined as its start index divided by total protein length. For proteins represented by ≥2 unique peptides, a mean relative position was calculated; proteins with mean values <0.3 were classified as exhibiting N-terminal bias and considered more likely to represent nascent polypeptide chains rather than bona fide ribosome-associated proteins.

### Surface GluA1 immunolabeling and sample preparation for STED

Hippocampal neurons were transduced on DIV 15 with an AAV encoding hSyn1-mVenusQ69M to visualize cellular morphology. On DIV 18, cells were treated with 1 µM TTX for 30 min and GluA1 surface immunolabeling was performed live for 5 min at 37°C using a mouse extracellular GluA1 antibody (1:200; Merck, #MAB2263). After live labeling, cells were washed twice with aCSF (130 mM NaCl, 5 mM KCl, 10 mM HEPES, 30 mM glucose, 1 mM MgCl_2_, 2 mM CaCl_2_, 1 µM TTX; pH 7.4) and fixed for 12 min at RT in 4% PFA-sucrose (4% PFA, 4% sucrose, 1 mM MgCl_2_, 0.1 mM CaCl_2_ in PBS, pH 7.4). Cells were then permeabilized with 0.1% Triton X-100 in PBS for 15 min, blocked with 4% goat serum in PBS for 1 hr at RT, and incubated overnight at 4°C with primary antibodies in blocking buffer. After 1 hr incubation at RT with secondary antibodies, cells were mounted in embedding media (Abberior MOUNT Solid Antifade, #MM-2013-2X15ML) and overlaid with a 12-mm (#1) glass coverslip. Three PBS washes were performed between all steps post-fixation, except between blocking and primary antibody incubation. Primary antibodies: rabbit anti-RPS11 (1:500; Bethyl, #S303-936A); chicken anti-GFP (1:2000; Abcam, #ab13970). Secondary antibodies: Alexa Fluor 488 goat anti-chicken IgY (1:500; Thermo, #A11039); Abberior STAR ORANGE goat anti-rabbit IgG (1:500; Abberior, #STORANGE-1002); Abberior STAR RED goat anti-mouse IgG (1:500; Abberior, #STRED-1001). For confocal imaging of RPL36A and surface GluA1, nontransduced DIV19 neurons were surface labeled for 10 min at 37°C using a rabbit extracellular GluA1 antibody (1:200; Merck, #ABN241) and stained for MAP2 (1:1000; Synaptic Systems, #188004) and RPL36A (1:500; Santa Cruz Biotechnology, #SC-100831). Cells were mounted in ProLong Gold Antifade Mountant with DAPI (Thermo Scientific). Secondary antibodies: Alexa Fluor 488 goat anti-guinea pig IgG (1:500; Thermo, #A11073); Alexa Fluor 568 goat anti-rabbit IgG (1:500; Thermo, #A21069); Alexa Fluor 647 goat anti-mouse IgG (1:500; Thermo, #A21236).

### STED image acquisition, processing and analysis

STED and confocal images were acquired using a commercial Abberior easy3D STED module integrated with an Olympus IX83 microscope equipped with the following: an Olympus UPLSAPO100XS 100×/1.35 N.A. objective, five excitation lasers (405 nm CW, 485 nm pulsed, 518 nm pulsed, 561 nm pulsed, 640 nm pulsed), four single-photon counting APD detectors, and two STED lasers (595 nm pulsed and 775 nm pulsed). The 488, 561, and 640 nm wavelengths were used to excite Alexa Fluor 488, Abberior STAR ORANGE, and Abberior STAR RED, respectively. Two-color 2D STED images were acquired by depleting Abberior STAR ORANGE and STAR RED with the 775 nm pulsed depletion laser with 0.75 ns gating time and dwell time of 15 µs. Overview confocal images of each cell, as well as individual dendritic segments, were acquired in the 488 (cell fill) channel. From the dendritic segment overview image, an ROI encompassing a single spine was defined. An 8-slice z-stack was then acquired at 0.18-µm step size in the 488 confocal channel, followed by a 3-slice z-stack with a 0.18-µm step size in all 5 channels (3 confocal, 2 STED) using the Aberrior LIGHTBOX software (pinhole: 1.0 AU, pixel size: 25 nm). The laser power and dwell time settings for each STED channel was kept constant across all experiments. The 5-channel z-stacks were deconvolved with SVI Huygens Essential v24.04 (Scientific Volume Imaging, The Netherlands, http://svi.nl), using the ‘Automatic’ algorithm, ‘Standard’ deconvolution strategy, and theoretical PSFs. The high-throughput analysis pipeline consisted of object segmentation on sum projections of deconvolved STED images (split-Bregman/MOSAIC suite, ImageJ2) followed by geometric analysis (MATLAB, vR2024b).

For segmentation, the following parameters were used: ‘Subpixel segmentation’, ‘Exclude Z edge’, Local intensity estimation ‘High’, Noise model ‘Poisson’. The binary mask generated by object segmentation was applied to the original intensity image to produce a masked intensity image for subsequent quantification. Objects located outside the spine (determined by a cell fill mask generated from the sum projection of the 488 confocal z-stack) were filtered and excluded from further analysis. Coordinates of regional intensity maxima (identified using the MATLAB function *imregionalmax*) were used to represent the centroids of surface GluA1 and RPS11 objects. Nearest-neighbor distances (NNDs) between centroids in the two channels were computed using the MATLAB function *knnsearch*. Normalized NNDs were calculated as y = r·√λ_target_, where r is the NND (µm) and λ_target_ is the target object density (objects/µm²) within the spine. For example, a GluA1 to RPS11 NND corresponds to the distance from one GluA1 object to its nearest RPS11 object, with λ_target_ defined as the number of RPS11 objects divided by the spine area. For comparison to complete spatial randomness (CSR), the probability density function (PDF) and cumulative distribution function (CDF) of the normalized NNDs were given by f(y) = 2πy*exp(-πy^2^) and F(y)=1−exp(-πy^2^), where y is the normalized NND. For comparison of per-spine median NNDs with a simulated dataset, surface GluA1 objects were held fixed and RPS11 objects were randomly repositioned within the spine ROI (using the MATLAB *randperm* function) for a total of 100 randomizations per spine. For each spine, the simulated value equals the median across its 100 simulated medians.

### RNA-seq library preparation

Total RNA was isolated from cortical synaptosomes across four biological replicates. RNA was extracted from samples using 1 mL TRIzol (Invitrogen, #10296010) and 200 μL chloroform (Sigma, #C2432). The aqueous phase was separated by centrifugation at 13,000×g for 15 min at 4°C. RNA from the aqueous phase was precipitated overnight at - 20°C with 1.5 μL GlycoBlue (Invitrogen, #AM9515), 0.1 volumes of 3 M sodium acetate, pH 5.5 (Invitrogen, #AM9470) and 3 volumes of 100% ethanol, then pelleted by centrifugation at 13,000×g for 40 min at 4°C. Pellets were washed with ice-cold 75% ethanol, air-dried for 10 min at RT, and resuspended in 10 μL RNase-free water (Invitrogen, #AM9937). RNA yield was quantified using the Qubit RNA HS assay (Invitrogen, #Q32855) and integrity was assessed using the Agilent RNA 6000 Pico kit. mRNA-seq libraries were generated from 200-300 ng of total RNA using the NEBNext Poly(A) mRNA Magnetic Isolation Module (New England Biolabs, #E7490) together with the Ultra II Directional RNA Library Prep kit for Illumina (New England Biolabs, #E7760) and NEBNext Multiplex Oligos for Illumina (New England Biolabs, #E6440). Libraries were sequenced on an Illumina NextSeq2000 using paired-end 61 bp reads.

### Genome alignment of RNA libraries and gene-level counting

Raw BCL files were demultiplexed using bcl2fastq2 (Illumina) with lane merging enabled, producing per-sample FASTQ files without lane splitting. Adapter and quality trimming were performed using fastp, providing removal of library-specific adapter sequences (via --adapter_fasta), poly-X trimming, low-complexity filtering (complexity threshold = 30), filtering of reads with mean Phred quality <20, and minimum read-length enforcement (≥21 nt). Cleaned reads were aligned to the rat (rn7) genome using STAR (v2.7.11b) with GTF-based splice junction annotation. Gene-level counts were generated from uniquely-mapped reads using featureCounts (Subread v2.0.8; (*87*)) with NCBI RefSeq rn7 GTF annotation (April 2025). For downstream analyses, gene-level raw counts were filtered to retain nuclear-encoded protein-coding genes.

### Definition of the synaptic transcriptome

Transcripts detected in synaptosome RNA-seq libraries were used to define the synaptic transcriptome. Genes were considered detected if CPM ≥ 0.1 in at least 3 of 4 biological replicates.

### Recovery of ribosome footprints

Ribosome footprints were isolated from IP samples (described above) across four biological replicates. For a subset of IPs (GluA1 and RPLP0 IPs), duplicate IPs were performed using the same input sample, providing technical IP replicates. Footprint extraction and recovery were carried out according to Ferguson et al. (*88*) with modifications. Briefly, RNA was extracted from IP samples using 1 mL TRIzol (Invitrogen, #10296010) and 200 μL chloroform (Sigma, #C2432). The aqueous phase was separated by centrifugation at 13,000×g for 15 min at 4°C. RNA from the aqueous phase was precipitated overnight at - 20°C with 1.5 μL GlycoBlue (Invitrogen, #AM9515) and 600 μL isopropanol (Sigma, #59304), then pelleted by centrifugation at 13,000×g for 15 min at 4°C. Pellets were air-dried for 30-60 min at RT and resuspended in 7 μL RNase-free water (Invitrogen, #AM9937). RNA yield was quantified using the Qubit RNA HS assay (Invitrogen, #Q32855) and a Qubit 3 Fluorometer (average RNA yield: 91.3 ng/uL (RPLP0 IP); 5.85 ng/uL (GluA1 IP); isotype control samples were ‘too low’ for quantification). For footprint size selection by denaturing PAGE, the remaining sample (∼6 μL) and 29-nt and 35-nt ssDNA size markers (IDT) were mixed with 2× Gel Loading Buffer II (Thermo, #AN8546G), denatured for 90 sec at 80°C, and placed on ice. Novex 15% TBE-urea gels (Thermo, #EC68855BOX) were pre-run in 1× TBE for 15 min at 200V. Ten picomoles of each size marker were loaded per lane. Blank wells containing 5 uL Novex TBE-Urea Sample Buffer (Thermo, #LC6876) were loaded in the first and last lanes of each gel, and in two lanes between each sample or size-marker lane. Gels were run for 65 min at 200V, stained for 3 min in SYBRGold (Invitrogen, #S11494) and visualized using a Dual LED blue/white light transilluminator (Invitrogen, #LB0100).

Footprints were excised by cutting the region just above the 35-nt size marker (below the xylene cyanol dye front) and slightly below the 29-nt marker.

29-nt size marker: GGT AAT AGC TTT TCT AGT CAG GTT AGG TC

35-nt size marker: ACT CTT TTA GTA TAA ATA GTA CCG TTA ACT TCC AA

RNA was extracted from the gel slice by crushing each slice into 2-3 pieces and submerging in 400 μL RNA extraction buffer (300 mM sodium acetate, pH 5.5, 1 mM EDTA, 0.25% SDS), snap-freezing at -80°C for 30 min and then incubating overnight at 23°C with shaking at 400 rpm. After centrifugation at 13,000×g for 10 min at RT, RNA was precipitated from the eluate (∼400 μL) at -80°C for 1 hr with 1.5 μL GlycoBlue and 3 volumes of 100% ethanol, then pelleted by centrifugation at 13,000×g for 15 min at 4°C. Pellets were washed with 1 mL ice-cold 70% ethanol, incubated at -20°C for 15 min, centrifuged at 13,000×g for 15 min at 4°C, air-dried for 10 min at RT and resuspended in 8 μL RNase-free water.

### Ribo-seq library preparation

Footprint libraries were generated using the D-Plex Small RNA-seq Kit for Illumina (Diagenode, #C05030001) and D-Plex Single Indexes for Illumina (Diagenode, #C05030010, #C05030011). RNA 3’-end dephosphorylation, RNA tailing, reverse transcription with template switching and PCR amplification (15 cycles) were done according to the manufacturer’s protocol. Libraries were purified using AMPure XP beads (Beckman, #A63881) followed by size selection on a 20% non-denaturing Novex TBE gel (Invitrogen, #EC63155BOX). The 200 bp products were excised and eluted overnight at 4°C in 100 μL Buffer EB (Qiagen). After centrifugation at 13,000×g for 15 min at 4°C, DNA was precipitated from the eluate with 1.5 μL GlycoBlue and 3 volumes of 100% ethanol, then pelleted by centrifugation at 13,000×g for 15 min at 4°C. Pellets were washed with 1 mL ice-cold 70% ethanol, incubated at -20°C for 15 min, centrifuged at 13,000×g for 15 min at 4°C, air dried for 10 min at RT and resuspended in 8 μL Buffer EB. DNA yield was quantified using the Qubit dsDNA HS assay (Invitrogen, #Q32854) and a Qubit 3 Fluorometer. Quality of DNA libraries were assessed using the Agilent High Sensitivity DNA kit (VWR, #AGLS5067-4626). Barcoded libraries were pooled and sequenced on an Illumina NextSeq2000 using single-end 100 bp reads and P4(100) XLEAP reagents.

### Genome and transcriptome alignment of Ribo-seq libraries

Raw BCL files were demultiplexed using bcl2fastq2 (Illumina) with lane merging enabled, producing per-sample FASTQ files without lane splitting. Adapter and quality trimming were performed using fastp, providing removal of library-specific adapter sequences (via --adapter_fasta), poly-X trimming, low-complexity filtering (complexity threshold = 30), and minimum read-length enforcement (≥15 nt). Unique molecular identifiers (UMIs) were extracted directly from read 1 using a fixed 12-nt offset strategy (--umi --umi_loc=read1 --umi_len=12 --umi_skip=4) to preserve molecule-level information prior to alignment.

Cleaned reads were aligned to the rat (rn7) genome using STAR (v2.7.11b) with GTF-based splice junction annotation and transcriptome-guided alignment (--quantMode TranscriptomeSAM GeneCounts). Output included coordinate-sorted BAMs, transcriptome-aligned BAMs, splice-junction tables, and per-gene read counts. NCBI RefSeq Annotation from April 2025 was used for transcriptome alignment. To remove PCR duplicates while preserving true ribosome-protected fragments, UMI-aware deduplication was performed using UMI-tools in array mode. UMIs carried through the read identifier were parsed using --extract-umi-method=read_id and --umi-separator=“:UMI_”, and duplicate groups were collapsed using directional adjacency with mapping-quality filtering (MAPQ = 255).

Deduplicated BAM files together with detailed duplication statistics were produced for all samples. This workflow ensured high-stringency preprocessing, accurate UMI handling, and reproducible quantification of ribosome footprints. Genome alignments were used for differential expression and genomic feature analyses, and visualization of coverage tracks.

Transcriptome alignments were processed with riboWaltz (v2.0; (*89*)) to compute quality control metrics, including read length distributions, frame periodicity and metagene profiles. Footprint reads of length 28-32 nt were used for P-site assignment and all subsequent riboWaltz-based analyses.

### Genomic feature analysis

Genomic coordinates for CDS, 5’ UTR, 3’ UTR and intronic regions were obtained from a RefSeq rn7 annotation in BED12 format from the UCSC Table Browser (*90*). Bedtools (v2.31.1; (*91*)) was used to convert BAM files into split-aware BED12 files. Reads were classified by genomic feature using stranded, split-aware intersections (bedtools intersect -s - split) applied sequentially to CDS, 5’UTR, 3’UTR and intronic regions, with remaining reads labeled as intergenic. Counts and fractions of reads mapping to each feature class were summarized as mean ± SEM across replicates.

### Coverage profile visualization

BAM files were used to compute CPM-normalized footprint coverage across RefSeq-annotated exons. Coverage profiles were collapsed onto linear transcript coordinates using BED12-defined exon structures and summarized as mean ± SEM across replicates.

### Gene-level counting

Counts per gene were calculated from uniquely-mapped, UMI-deduplicated reads aligned to the genome described above using featureCounts (Subread v2.1.0; (*87*)), with reads assigned to annotated exons and summarized at the gene (meta-feature) level. Only a single transcript isoform, with the highest APPRIS principal score (*92*), was considered per gene. For samples with technical IP replicates derived from the same biological material, gene-level raw counts were summed to yield one count profile per biological replicate. For downstream analyses, gene-level raw counts were filtered to retain nuclear-encoded protein-coding genes.

### Definition of the synaptic translatome

Footprints detected in RPLP0 or GluA1 IPs from ribosome-enriched fractions from cortical synaptosomes (i.e., ‘synaptic fraction cushion’) were used to define the synaptic translatome. Genes were considered detected if CPM ≥ 1 in at least 3 of 4 biological replicates in either IP (union of detected genes across IPs).

### Differential expression analysis (ribosome profiling)

Differential expression analysis was performed using the voom-limma pipeline (*93*, *94*) on gene-level raw counts pre-filtered to retain nuclear-encoded protein-coding genes. To retain genuine differences in IP enrichment across samples, no composition-based library-size normalization was applied, resulting in effective library sizes equal to the observed library sizes. This preserves global abundance shifts while avoiding normalization artifacts that arise when the assumption that most genes are unchanged across conditions is violated, as is typical in IP/pulldown experiments. Raw counts were transformed to log_2_-counts per million, and precision weights were estimated using the voom method. Weighted linear models were fitted using *voomLmFit*, followed by empirical Bayes moderation using *eBayes*. Sample-level QC was performed using limma’s multidimensional scaling (*plotMDS*) applied to the voom-transformed expression values (top 1,000 most variable genes).

For the synaptic vs. cytosolic translatome comparison (i.e., RPLP0 IP from the synaptic fraction vs. RPLP0 IP from the cytosolic fraction; Fig.4C), lowly expressed genes were removed by CPM filtering in edgeR (*95*); genes were retained if CPM ≥ 1 in at least 3 of 4 biological replicates in either synaptic or cytosolic RPLP0 IPs. A bioinformatic filter focusing on excitatory neuronal transcripts (*9*, *10*) was additionally applied to the gene-level raw counts matrix to enable direct comparison to neuropil and somata translatomes reported previously (*10*). Synapse-enriched and cytosol-enriched genes were defined by the voom-limma model (FDR ≤ 0.05 and |log_2_FC| ≥ log_2_(1.3)), after excluding genes that were not significantly enriched in either the synaptic RPLP0 IP vs. synaptic isotype control comparison or the cytosolic RPLP0 IP vs. cytosolic isotype control comparison (table S5).

For the GluA1 IP vs. RPLP0 IP comparison (Fig. 4H), lowly expressed genes were removed by CPM filtering in edgeR (*95*); genes were retained if CPM ≥ 1 in at least 3 of 4 biological replicates in either the GluA1 IP or RPLP0 IP. GluA1-enriched and RPLP0-enriched genes were defined using the same statistical thresholds (FDR ≤ 0.05 and |log_2_FC| ≥ log_2_(1.3)), after excluding MitoCarta-annotated mitochondrial genes (‘mitochondrial’) and genes that were not significantly enriched in either the GluA1 IP vs. isotype control comparison or the RPLP0 IP vs. isotype control comparison (‘non-IP-enriched’; table S7).

### Comparative rank-based analysis of synaptic and neuropil translatomes

Gene-level footprint counts from both datasets were filtered to retain nuclear-encoded protein-coding genes, and contaminants were excluded using a previously described neuronal transcript filter (*9*, *10*). Genes were restricted to a shared expression universe defined as those detected in both datasets (CPM ≥ 1 in at least 3 of 4 synaptic RPLP0 IP replicates or in at least 2 of 3 neuropil Ribo-seq replicates). For each compartment, footprint abundance was summarized as the mean log_2_-CPM across replicates, and genes were ranked by decreasing abundance. Downstream analyses focused on synapse-enriched genes identified in the synaptic vs. cytosolic translatome comparison, which were classified based on percentile rank shifts within the rank distribution: genes with rank differences ≤5% were defined as rank-stable, whereas genes with >50% upward or downward shifts in rank from neuropil to synapse were classified as rank-increased and rank-decreased, respectively.

### GO enrichment analysis (ribosome profiling)

All GO enrichment analyses were performed using clusterProfiler (*84*), with internal significance filtering (pvalueCutoff = 0.05; qvalueCutoff = 0.05) and multiple testing controlled by the Benjamini-Hochberg procedure. Where applicable, redundant GO terms were removed using rrvgo (*85*), which applies GOSemSim semantic similarity (Rel metric, similarity threshold 0.3; (*86*)).

GO analysis was performed on the ‘core’ synaptic translatome (fig. S8D), defined as mRNAs detected in both synaptic RNA-seq and Ribo-seq datasets (intersection; n=12,283). The background comprised all transcripts detected in either dataset (union; n=14,992). For visualization, the top 15 non-redundant terms (ranked by adjusted p-value) were selected.

GO analyses were performed for synapse-enriched and cytosol-enriched transcripts (Fig. 4D). The background comprised all genes included in the differential expression model. For visualization, the top 5 non-redundant terms (ranked by adjusted p-value) were selected. GO analyses were performed for synapse-, neuropil-, and somata-enriched transcripts with no internal significance filtering (pvalueCutoff = 1; qvalueCutoff = 1; Fig. 4G), with enrichment tested against the corresponding background gene universe for each comparison. Only GO terms present across all three compartments and supported by ≥10 genes and statistical significance (FDR<0.05) in at least one condition were retained for downstream analysis. GO terms were not filtered for redundancy. GO terms were ranked by mean log_2_ fold enrichment across compartments, which was used as an enrichment score, and the top 15 terms were selected for comparative visualization.

GOCC enrichment was performed for rank-increased transcripts with no internal significance filtering (pvalueCutoff = 1; qvalueCutoff = 1; Fig. 4G). The background comprised all synapse-enriched genes that were also detected in the neuropil (n=976 genes). GO terms were not filtered for redundancy. For visualization, the top 5 terms (ranked by adjusted p-value) were selected.

GO analyses were performed for RPLP0-enriched and GluA1-enriched transcripts with no internal significance filtering (pvalueCutoff = 1; qvalueCutoff = 1; Fig. 4J; fig. S8E,F). The background comprised all genes included in the differential expression model. For visualization, the top 10 non-redundant terms (ranked by adjusted p-value) were selected, and enrichment statistics were extracted from the full enrichGO results (i.e., before rrvgo filtering) to ensure that each selected term could be plotted for both groups.

### Crosslinking of ribosome cushion samples

Ribosome-enriched fractions from cortical synaptosomes were cross-linked with 5 mM azide-A-DSBSO for 20 min at RT with constant mixing. The reaction was quenched by adding Tris-HCl, pH 8.0 to a final concentration of 20 mM and incubated for 30 min at RT. An equal volume of 8% SDS in 50 mM triethylammonium bicarbonate (TEAB) was then added to reach a final SDS concentration of 4%. Samples were reduced with 5 mM DTT at 37°C for 30 min, followed by alkylation with chloroacetamide (CAA) at 37°C for 30 min in the dark. Samples were subsequently cleaned and digested using the SP3 protocol as described previously (*96*). Briefly, a mixture of hydrophobic and hydrophilic SeraMag beads was added to the samples, followed by addition of 100% acetonitrile (ACN) to a final concentration of 50% (v/v). After incubation for 20 min on a shaker, the beads were immobilized using a magnetic rack and the supernatant was removed. The beads were washed three times with ACN. For digestion, beads were resuspended in 50 mM TEAB containing LysC (1:100, enzyme:protein, w/w) and trypsin (1:50, enzyme:protein, w/w), and proteolysis was carried out at 37°C overnight with shaking. Cross-linked peptides were enriched using DBCO beads by incubation at RT for 3 hr. The beads were then incubated with 0.5% SDS at 37°C for 15 min, followed by three consecutive washes with 0.5% SDS, 8 M urea, and Milli-Q water, respectively. Cross-linked peptides were eluted by incubating the beads with 2% trifluoroacetic acid (TFA) at 37°C for 1 hr. Enriched peptides were further cleaned using strong cation exchange (SCX) stage tips as described previously (*97*). Cross-linked peptides were fractionated by high-pH reversed-phase chromatography using a Phenomenex Gemini C18 column on an Agilent 1260 Infinity II UPLC system. A 100 min gradient was applied, generating 192 fractions, which were concatenated into 24 final fractions. Each fraction was subsequently cleaned using SP3 beads prior to LC-MS/MS analysis.

### LC-MS/MS data acquisition and analysis (XL-MS)

High-pH fractions were analyzed by LC-MS/MS on an Orbitrap Exploris 480 coupled to a Vanquish neo UHPLC system (Thermo Fisher Scientific). Peptides were separated on an in-house packed analytical reverse-phase column (Poroshell 120 EC-C18, 2.7 µm, Agilent Technologies) with 180 min gradients at a flow rate of 0.25 µL/min. FAIMS Pro interface (Thermo Fisher Scientific) was installed in front of the ion source, with internal compensation voltage (CV) stepping at –50, –60, –75 V. MS1 scans were acquired in the Orbitrap analyzer with a resolution of 120,000 over an m/z range of 375–1400, using a standard AGC target and a maximum injection time of 50 ms. Precursor ions with charge states between +4 and +6 and an intensity above 2.0 × 10⁴ were selected for MS2. MS2 scans were acquired using stepped higher-energy collisional dissociation (HCD) with normalized collision energies of 18, 31, 32, 33. Fragment ions were analyzed in the Orbitrap at a resolution of 45,000 using a 1.6 m/z isolation window, a maximum injection time of 90 ms, and a normalized AGC target of 200%.

For crosslink identification, a reduced protein database (2,028 sequences) was generated from proteins identified by searching non-cross-linked peptide data against the reviewed *Rattus norvegicus* UniProt database (8,321 sequences). Cross-linked samples were searched against the reduced database using pLink3 (v3.0.17; https://github.com/pFindStudio/pLink3) and Scout (v1.5.1; https://github.com/theliulab/Scout) with the following settings: enzyme specificity, trypsin; fixed medication, cysteine carbamidomethylation; variable modification, methionine oxidation, protein N-term acetylation; the DSBSO mass, 308.039 Da; long arm, 236.018 Da; short arm, 54.011 Da; cross-linking site, lysine side chain and protein N-termini. For pLink3, identifications were filtered at a peptide-pair level false discovery rate (FDR) of 1%. For Scout, a 1% FDR was applied at all levels, including cross-linked spectrum matches (CSMs), residue pairs, and protein–protein interactions (PPIs). pLink3 results were exported at the CSM level and aggregated to residue-pair and PPI levels, whereas Scout results were exported directly at CSM, residue-pair, and PPI levels. Results from both search engines were combined and reported in table S8, with the source of each identification indicated. Shared peptides were grouped, and for annotation at the PPI level, the first protein listed within each ProteinAmbiguityGroup was used (table S8). For each PPI, we provide the supporting biological replicate(s) and search engine(s), as well as the corresponding residue pairs and their detection frequencies. These annotations allow users to independently evaluate the confidence and reproducibility of individual crosslinks. Ambiguous peptide assignments were resolved by manual curation (table S8). For instance, crosslinks originating from peptides shared between RPS27A (Ubiquitin-ribosomal protein eS31 fusion protein, P62982) and UBB (Polyubiquitin-B, P0CG51) were excluded from the RPS27A-specific analysis. Similarly, for crosslinks involving CaMKIIα with peptides shared among RPL35, RPL19 and ITPR1 (Inositol 1,4,5-triphosphate receptor), we assigned the interactions to RPL35 and RPL19. This decision was based on their higher abundance (∼5- to 13-fold greater than ITPR1) in our IP-MS dataset using comparable ribosome-enriched fractions from cortical synaptosomes (fig. S9D), indicating that RPL35 and RPL19 were more likely interaction partners than ITPR1 in the XL-MS dataset. Additional evidence further supports that CaMKIIα interacts with ribosomal proteins rather than ITPR1: a) ITPR1 did not contain any other crosslinks; b) CaMKIIα crosslinked to the ribosome-associated elongation factor EEF1A2, which crosslinked to RPS24, RPL9 and RPL12; and c) structural mapping of the crosslinked peptide on ITPR1 (PDB 6MU1) showed that it resides in a membrane-embedded region of the channel, suggesting that this site would not be accessible for crosslinking to CaMKIIα.

### Modeling of CaMKIIα and the 80S ribosome

HADDOCK 2.4 was used for molecular docking (*98*). Due to input size limitations, only half of the holoenzyme form of CaMKIIα (PDB 5U6Y) was used. Structures of RPL35 and RPL19 were obtained from AlphaFold3 predictions. Disordered regions in RPL35 and RPL19 were modeled as fully flexible segments during HADDOCK docking. Crosslinks from the dataset were incorporated as distance restraints with an upper limit of 35 Å as an input for the docking. The 80S ribosomal structure (PDB 7QGG) was superimposed in ChimeraX using Matchmaker, aligning chain i or chain S of 7QGG to RPL35 or RPL19, respectively. The distances between crosslinked residues were assessed on the proposed model where crosslinks that satisfied the distance restraints were highlighted in green, whereas violations were indicated in yellow.

### Puromycin labeling after CaMKII inhibition

DIV 18-19 hippocampal neurons were treated with either 10 μM KN-93 (Tocris, #5215) or 10 μM myr-AIP (Tocris, #5959) for 30 min at 37°C, or left untreated. Cells were metabolically labeled for 5 min at 37°C with 3 μM puromycin (Sigma, #P8833). Anisomycin control cells were pre-treated with 40 μM anisomycin (Sigma, #A9789) for 30 min at 37°C before puromycin labeling. After labeling, cells were washed twice in the original medium, fixed for 20 min in 4% PFA-sucrose (4% PFA, 4% sucrose, 1 mM MgCl_2_, 0.1 mM CaCl_2_ in PBS, pH 7.4), permeabilized for 15 min with 0.5% Triton X-100 in blocking buffer (4% goat serum in PBS, pH 7.4). After blocking with blocking buffer for 1 hr at RT, cells were incubated with primary antibodies in blocking buffer for 1 hr at RT, followed by secondary antibodies in blocking buffer for 30 min at RT. Three PBS washes were performed between all steps post-fixation, except between blocking and primary antibody incubation. Cells were kept at 4°C in PBS until imaging. Primary antibodies: mouse anti-puromycin (1:3000, Kerafast, #Kf-Ab02366-1.1); guinea pig anti-MAP2 (1:1000; Synaptic Systems, #188004); chicken anti-Homer1 (1:500; Synaptic Systems, #160019). Secondary antibodies: Alexa Fluor 405 goat anti-guinea pig IgG (1:500; Abcam, #ab175678); Alexa Fluor 568 goat anti-mouse IgG (1:500; Thermo, #A11031); Alexa Fluor 647 goat anti-chicken IgY (1:500; Abcam, #ab150175).

### Puro-PLA after intrabody-mediated GluA1 sequestration

DIV 14-15 hippocampal neurons were transfected with pAAV-hSyn-GluA1scFv-mNeon-KDEL (Addgene #230056; gift from Matthew Kennedy) or pAAV-hSyn-GFPnanobody-mNeon-KDEL using Lipofectamine 2000 (Invitrogen) according to the manufacturer’s protocol. Detection of newly synthesized proteins by puromycin labeling and proximity ligation (Puro-PLA) was performed 3.5 d after transfection (at DIV 18-19) as previously described (*55*). Cells were metabolically labeled for 5 min at 37°C with 3 µM puromycin (Sigma, #P8833). Anisomycin control cells were pre-treated with 40 µM anisomycin (Sigma, #A9789) for 30 min at 37°C before puromycin labeling. After labeling, cells were washed twice in the original medium, fixed for 20 min in 4% PFA-sucrose (4% PFA, 4% sucrose, 1 mM MgCl_2_, 0.1 mM CaCl_2_ in PBS, pH 7.4), permeabilized for 15 min with 0.5% Triton X-100 in blocking buffer (4% goat serum in PBS, pH 7.4), and then blocked for 1 hr in blocking buffer. Cells were then incubated for 1.5 hr at RT in blocking buffer containing primary antibodies against puromycin (1:3500; Kerafast, #Kf-Ab02366-1.1) and the candidate protein. After three PBS washes, PLA was performed using the Duolink In Situ PLA kit (Sigma, #DUO92008) according to the manufacturer’s protocol. Briefly, cells were incubated with PLA probes anti-rabbit PLUS (1:10; Sigma, #DUO92002) and anti-mouse MINUS (1:10; Sigma, #DUO92004) in blocking buffer for 1 hr at 37°C, washed five times in Wash Buffer A (0.01 M Tris, 0.15 M NaCl, 0.05% Tween 20) and incubated for 30 min at 37°C with ligation buffer. Cells were then washed three times in Wash Buffer A and incubated for 100 min at 37°C with the amplification reaction. To terminate amplification, cells were washed three times in Wash Buffer B (0.2 M Tris, 0.1 M NaCl, pH 7.5) and then washed three times in PBS. After blocking for 1 hr in blocking buffer, cells were incubated with primary antibodies against MAP2 (1:1000; Synaptic Systems, #188004) to label dendrites and Homer1 (1:500; Synaptic Systems, #160019) to label excitatory postsynapses for 1 hr at RT. Cells were then washed with PBS, incubated with secondary antibodies for 30 min at RT, washed and stored at 4°C in PBS until imaging. Antibodies for PLA of candidate proteins: rabbit anti-Drebrin (1:500; Sigma, #D3816); rabbit anti-CaMKII [EP1829Y] (1:2000; Abcam, #ab52476). Secondary antibodies: Alexa Fluor 405 goat anti-guinea pig IgG (1:500; Abcam, #ab175678); Alexa Fluor 647 goat anti-chicken IgY (1:500; Abcam, #ab150175).

### Image acquisition and analysis for Puro-PLA and puromycin labeling experiments

Samples were imaged within one week of labeling on either a Zeiss LSM780 or LSM880 confocal microscope, and imaging for each experiment (e.g., *Camk2a* Puro-PLA) was performed on the same instrument to ensure consistency. Imaging on the LSM780 microscope was performed using an alpha Plan-Apochromat 63×/1.46 Oil Korr M27 objective and ZEN 2.3 SP1 FP3 (black edition) software (Zeiss; v14.0.29.201). Imaging on the LSM880 microscope was performed using a Plan-Apochromat 63×/1.4 Oil objective and ZEN 2.3 SP1 FP3 (black edition) software (Zeiss; v14.0.29.201). MAP2- or mNeon-positive neurons were selected for imaging and quantification if they had a pyramidal-shaped soma, a prominent apical dendrite, visible dendritic spines, and a dendritic arbor that could be unambiguously attributed to a single neuron. The z-stack was set to encompass the entire dendritic volume (∼17 slices) using an optical slice thickness of ∼0.38 µm. Laser power and detector gain were adjusted to avoid pixel saturation, and image acquisition settings were held constant within each experiment. Quantification was performed on maximum-intensity projections of the raw z-stack images in ImageJ2 with an in-house script. For quantification of Puro-PLA images, a 50-80 µm segment of the apical dendrite situated 20-50 µm from the soma was manually selected as the ROI. For quantification of puromycin labeling images, the entire apical dendritic arbor was manually selected as the ROI. For each ROI, two masks were generated: a ‘synaptic mask’ defined by the Homer1 signal, and a mask encompassing both synaptic and non-synaptic regions derived from the combined MAP2 and Homer1 signal. The background-subtracted integrated fluorescence intensity of the puromycin/nascent protein signal within each mask was quantified and normalized to the corresponding mask area. Non-synaptic fluorescence intensity was obtained by subtracting the background-subtracted integrated fluorescence intensity within the synaptic mask from that within the combined (synaptic + non-synaptic) mask, and normalizing to the non-synaptic area (defined as the area of the combined mask minus the area of the synaptic mask). Images were expanded and smoothed in ImageJ2 for display only.

---

**Fig. S1.**
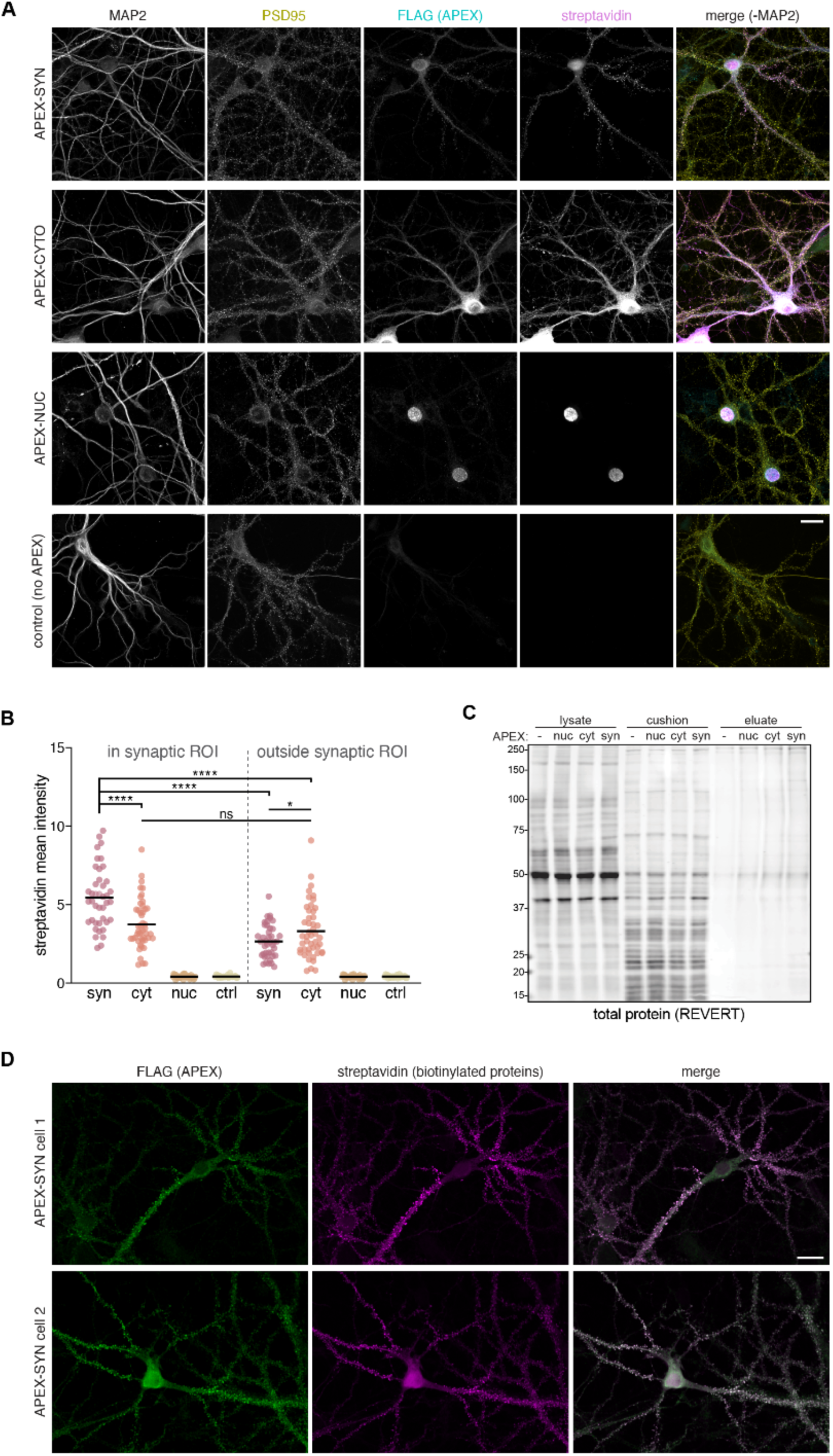
Biotinylation with APEX-SYN is more synaptically enriched than APEX-CYTO. (**A**) Images of the localization and biotinylation patterns of APEX-NUC, APEX-CYTO, APEX-SYN, and control (-APEX) after 1 min of labeling in primary rat hippocampal neurons. The streptavidin and FLAG signals are displayed at the same intensities across conditions. Scale bar, 20 µm. (**B**) Quantification of the biotinylated signal (streptavidin mean intensity) in synaptic regions (overlapping with PSD95) and non-synaptic regions from a 10-20 µm dendritic segment. Horizontal bars represent the mean; *p<0.05; **p<0.01; ***p<0.001; ****p<0.0001, One-way ANOVA, Tukey’s multiple comparisons test; n=28-52 cells from 3 independent cultures. (**C**) Image of total protein stain (REVERT) from one of five experimental replicates (same replicate shown in Fig. 1C). (**D**) Two additional example images of the localization and biotinylation pattern of APEX-SYN after 1 min of labeling. Scale bar, 20 µm.

**Fig. S2.**
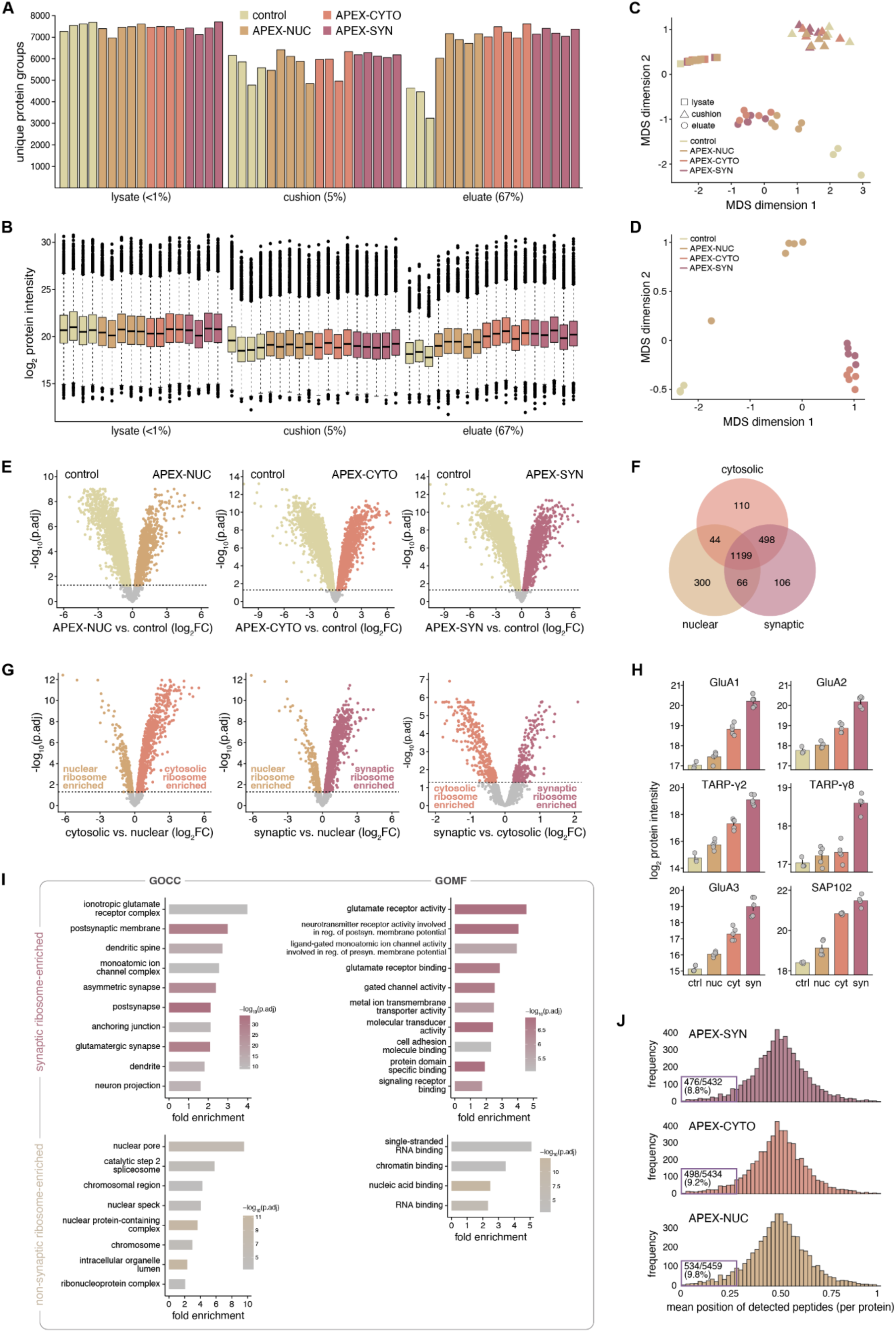
Spatial mapping of the neuronal ribosome protein interactomes. (**A**) Number of unique protein groups quantified per replicate from control (-APEX), APEX-NUC, APEX-CYTO, and APEX-SYN lysates, ribosome cushion pellets, and eluate fractions. (**B**) Distribution of log_2_ protein intensities across all samples. (**C**) Multidimensional scaling (MDS) plot showing sample similarity based on pairwise log_2_ protein intensity differences across the 1,000 most variable proteins. (**D**) MDS plot based on imputed and quantile-normalized protein intensities for eluate samples only. (**E**) Volcano plots showing protein enrichment in APEX-NUC vs. control eluate fractions (left), APEX-CYTO vs. control eluate fractions (center), and APEX-SYN vs. control eluate fractions (right). Proteins that were significantly enriched in the APEX-NUC, APEX-CYTO, and APEX-SYN eluate fractions are labeled in tan, coral, and rose, respectively (FDR<0.05, logFC>0). (**F**) The number of shared and potentially unique interactors of nuclear, cytosolic, and synaptic ribosomes. (**G**) Volcano plots of differentially expressed proteins in the cytosolic vs. nuclear ribosome interactomes (left), synaptic vs. nuclear ribosome interactomes (center) and synaptic vs. cytosolic ribosome interactomes (right). Significantly enriched proteins are highlighted (FDR<0.05). (**H**) Relative intensities of six selected proteins that were enriched in the synaptic ribosome interactome and associated with the GO term “AMPA glutamate receptor complex”. Data are shown as mean ± SEM. (**I**) GO overrepresentation analysis of enriched synaptic or non-synaptic ribosome-associated proteins. Shown are the top 10 enriched GOCC and GOMF terms. (**J**) N-terminal bias assessment to differentiate between bona fide ribosome-associated proteins and nascent polypeptide polypeptide chains.

**Fig. S3.**
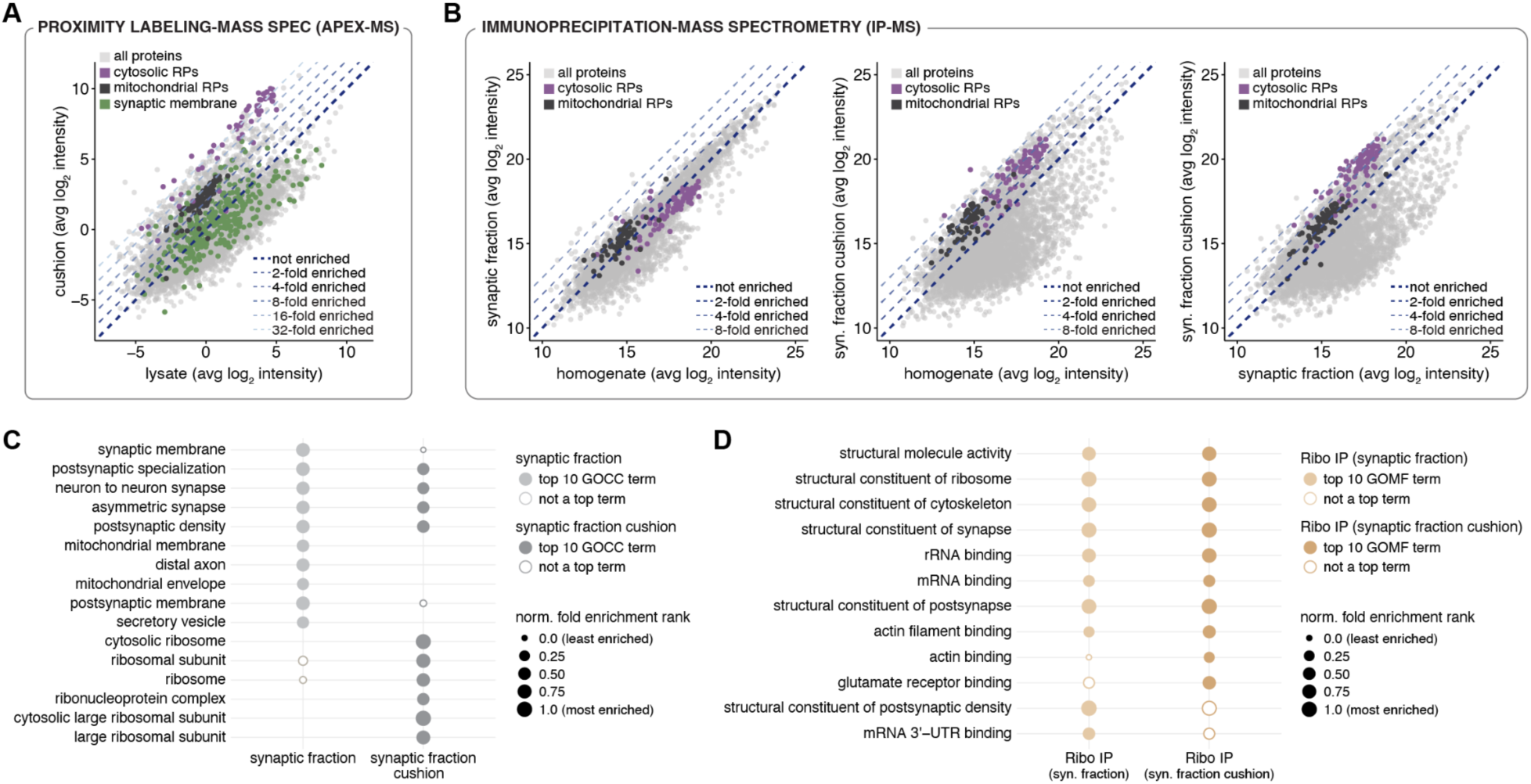
Cytosolic ribosomes, not synaptic membrane proteins, are enriched by sucrose cushion ultracentrifugation. AMPA receptors are enriched independent of ribosome IP input (−/+ cushion). (**A**) Specific enrichment of cytosolic ribosomal proteins (RPs) following sucrose cushion ultracentrifugation of primary rat cortical neuron lysates. Note the relatively modest enrichment of mitochondrial RPs. Sucrose cushion ultracentrifugation does not enrich for synaptic membrane proteins. Proteins associated with the GOCC term ‘synaptic membrane’ are labeled in green. Data points represent the median average of log_2_ protein intensities across replicates. (**B**) Sucrose cushion ultracentrifugation is necessary for enrichment of cytosolic ribosomes in rat cortical synaptosomes. *Left*: Cytosolic ribosomes are de-enriched in the synaptic fraction relative to the homogenate. *Center*: Cytosolic ribosomes are enriched in the synaptic fraction cushion relative to the homogenate. *Right*: Cytosolic ribosomes, and to a lesser extent mitochondrial ribosomes, are enriched in the synaptic fraction cushion relative to the synaptic fraction. Data points represent the median average of log_2_ protein intensities across replicates. (**C**) GO overrepresentation analysis of the proteins in the synaptic fraction and synaptic fraction cushion. Open circles denote significant terms (FDR<0.01) that did not rank among the top 10. ‘Synaptic membrane’ is a top term in the synaptic fraction but not in the cushion, where it is weakly enriched. In contrast, postsynaptic terms including ‘postsynaptic density’ and ‘postsynaptic specialization’ persist as top terms, consistent with PSD proteins as bona fide ribosome interactors. (**D**) GO overrepresentation analysis for proteins in the ribosome IP derived from the synaptic fraction and synaptic fraction cushion. Open circles denote significant terms (FDR<0.01) that did not rank among the top 10. ‘Glutamate receptor binding’ is significantly enriched in both conditions but becomes a top-ranked term only following inclusion of the cushion step.

**Fig. S4.**
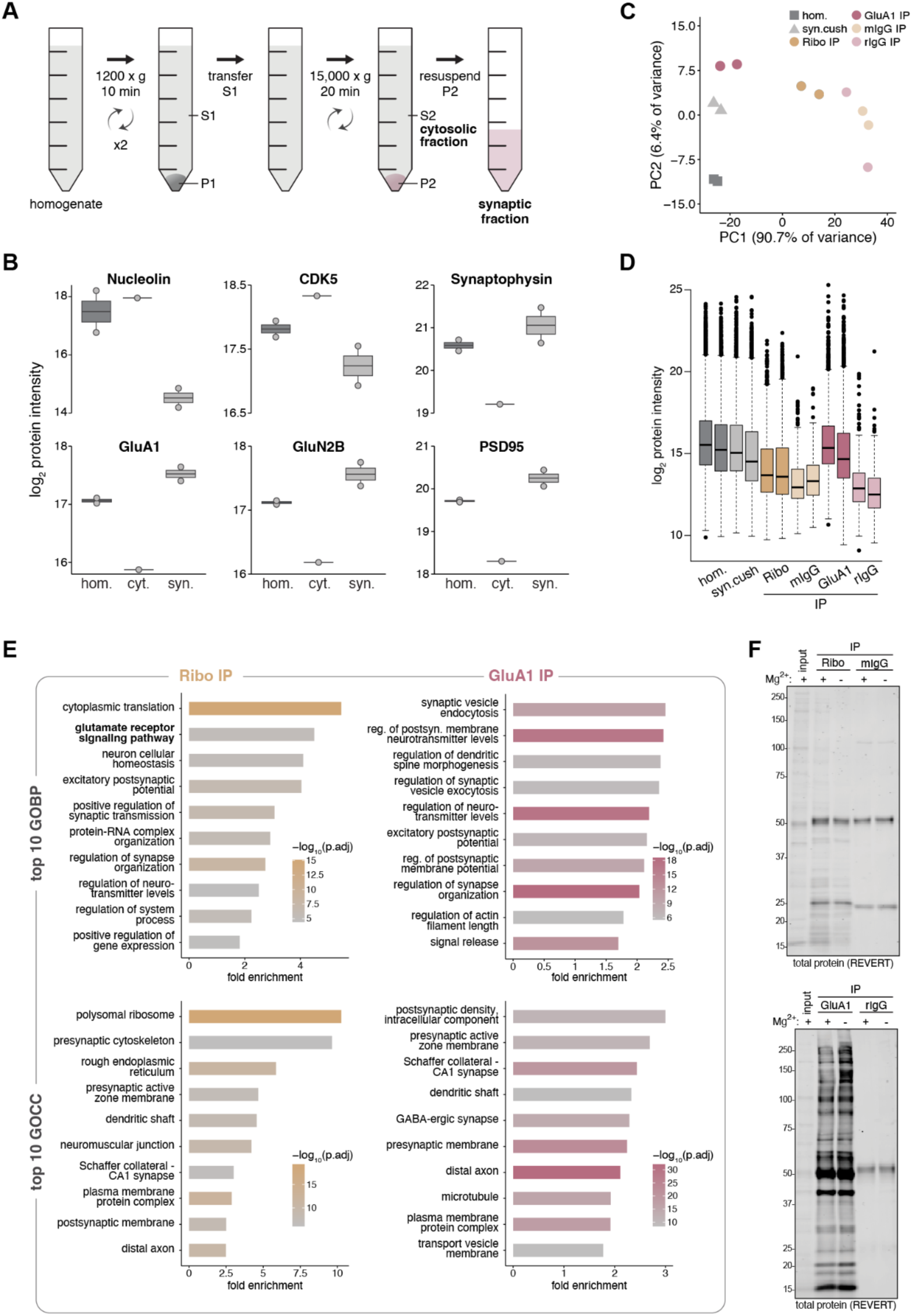
Synaptic ribosomes interact with AMPA receptor complex proteins. (**A**) Overview of synaptosome preparation with SynPER. Shown in bold are the fractions used in Fig. 4. (**B**) Synaptic proteins are enriched in SynPER synaptosomes. Shown are the relative intensities of six selected proteins in the tissue homogenate, cytosolic fraction, and synaptic fraction (before cushion ultracentrifugation). Nucleolin serves as a nuclear marker and CDK5 serves as a cytosolic marker. (**C**) Principal component analysis (PCA) of the log_2_ protein intensities in the tissue homogenate, synaptic fraction cushion and IP eluate fractions. (**D**) Overall log_2_ protein intensities per replicate from the tissue homogenate, synaptic fraction cushion and IP eluate fractions. (**E**) GO overrepresentation analysis for proteins in the ribosome IP and GluA1 IP eluate fractions. Shown are the top 10 enriched GOBP and GOCC terms. (**F**) Total protein stain images for the immunoblots in Fig. 2G.

**Fig. S5.**
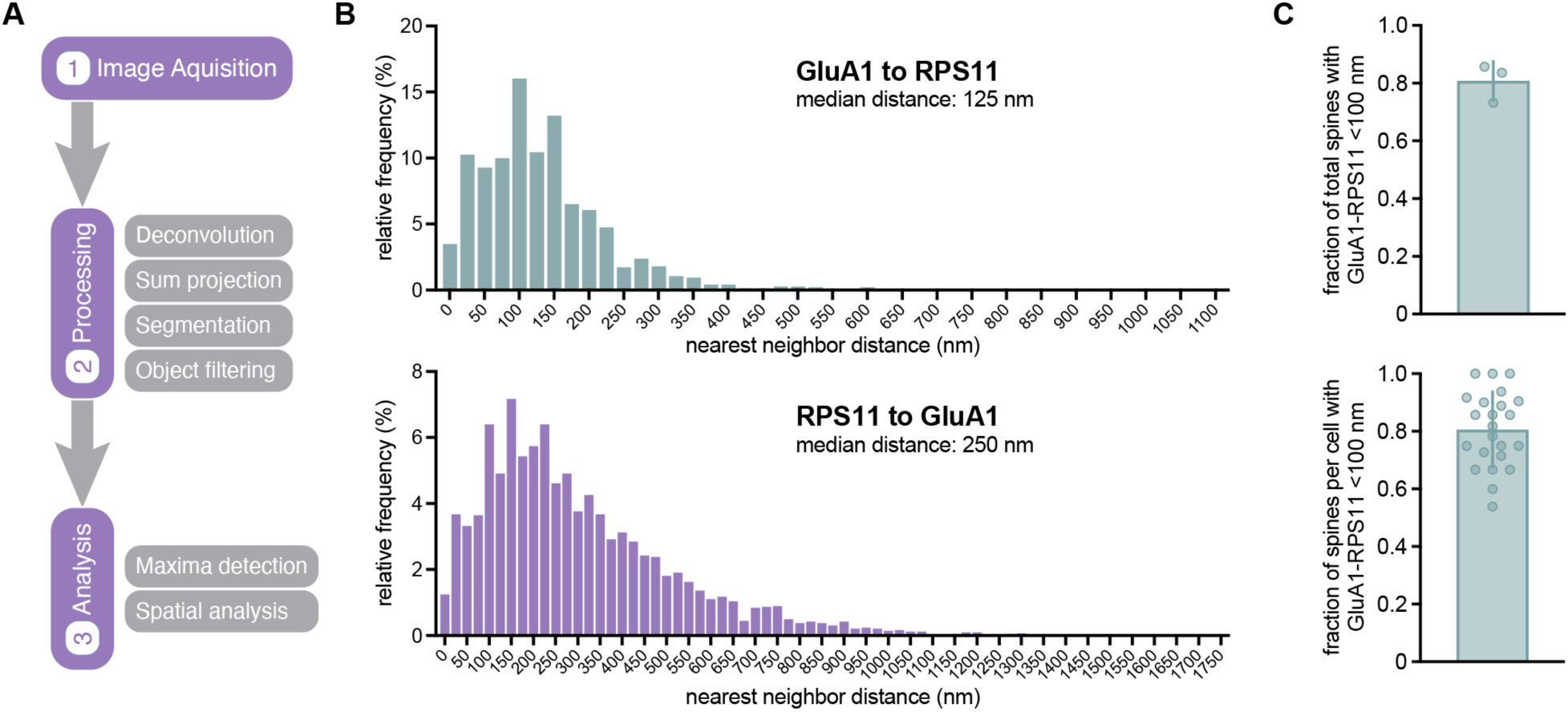
Nanoscale imaging of the AMPA receptor-ribosome association in dendritic spines. (**A**) Overview of STED imaging analysis pipeline. (**B**) Relative frequency histograms of nearest-neighbor distances. GluA1 to RPS11: n=1524 distances (puncta); RPS11 to GluA1: n=4254 distances (puncta). (**C**) Fraction of total spines per replicate (top) or spines per cell (bottom) containing at least one ribosome located within 100 nm of surface GluA1. Error bars show mean ± SEM; n=323 spines from 100 dendritic segments from 23 cells from 3 independent cultures.

**Fig. S6.**
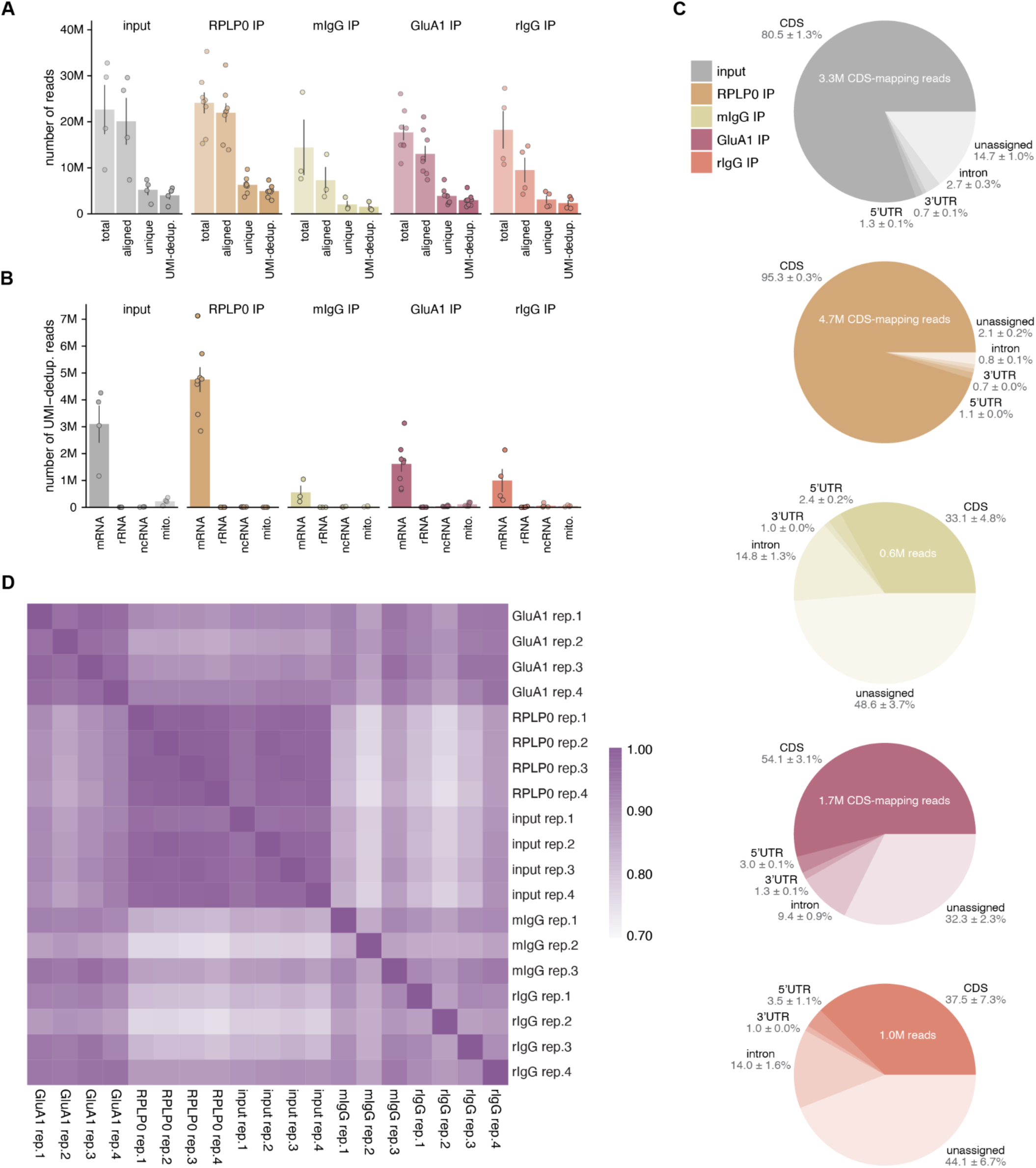
Mapping statistics of ribosome profiling data. (**A**) Footprint read counts across sequential filtering steps: total, aligned, uniquely mapped, and UMI-deduplicated reads. Error bars show mean ± SEM. (**B**) Number of footprint reads (UMI-deduplicated) assigned to mRNA, rRNA, ncRNA, and mitochondrial genome-encoded transcripts. Error bars show mean ± SEM. (**C**) Percentage of footprints (UMI-deduplicated reads; mean ± SEM) mapping to various genomic features including the 5’UTR (untranslated region), CDS (coding sequence), 3’UTR and introns. (**D**) Heatmap of Pearson’s r correlations between biological replicates of the synaptic fraction cushion (input), RPLP0 IP, GluA1 IP, and isotype control footprint libraries (log_2_CPM).

**Fig. S7.**
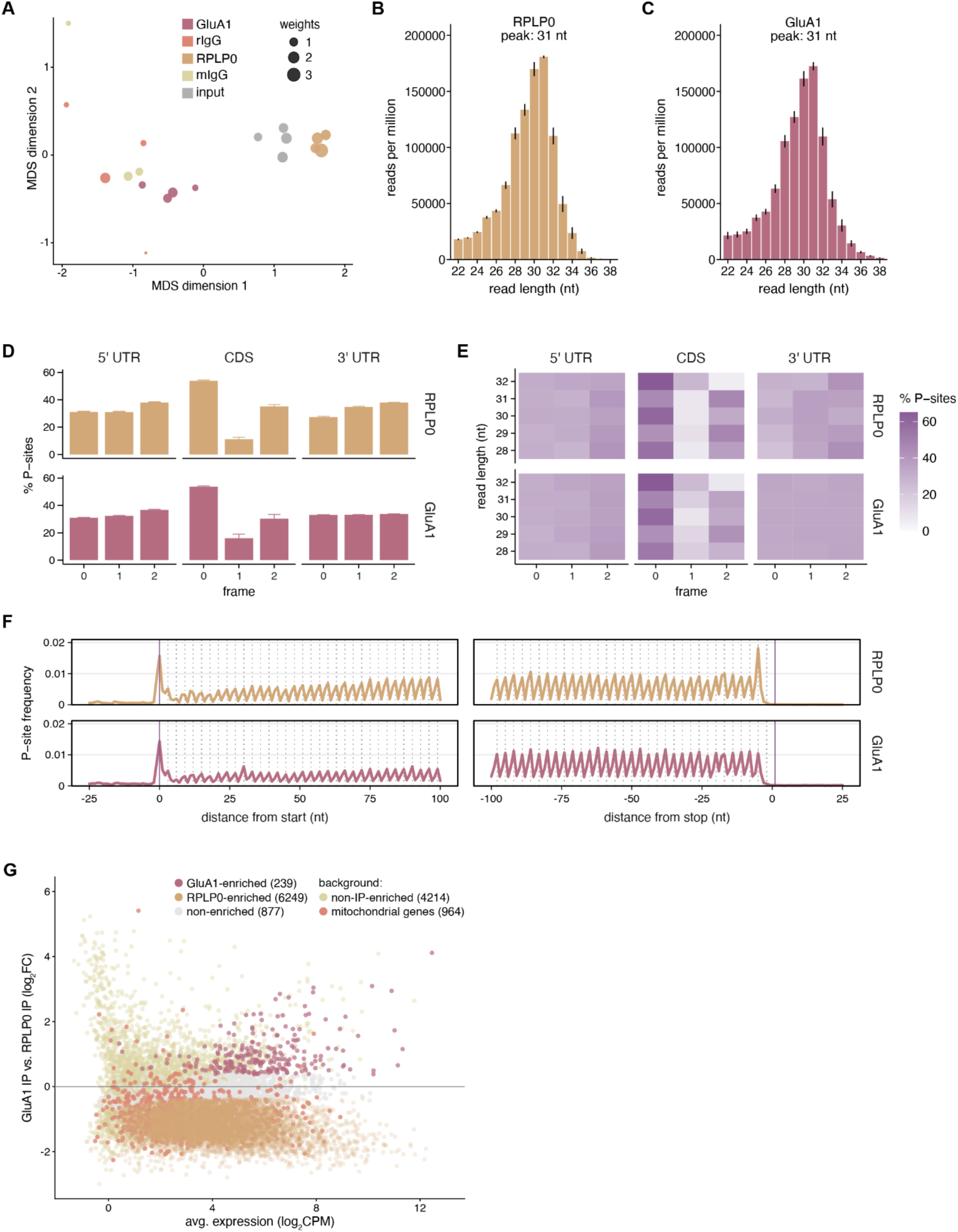
Quality metrics of ribosome profiling data. (**A**) Multidimensional scaling (MDS) plot showing sample similarity based on the 1,000 most variable genes. (**B**) Distribution of footprint read lengths obtained in Ribo IP libraries. Error bars show mean ± SEM. (**C**) Distribution of footprint read lengths obtained in GluA1 IP libraries. Error bars show mean ± SEM. (**D**) Percentage of P-sites in each reading frame (0, +1, +2) across the 5’ UTR, CDS, and 3’ UTR, using 28-32 nucleotide footprints. Error bars show mean ± SEM. (**E**) Percentage of P-sites in each reading frame (0, +1, +2) across the 5’ UTR, CDS, and 3’ UTR, stratified for read length. (**F**) Metagene plots of P-site density centered on start and stop codons, with each nucleotide position showing the summed P-site count across transcripts. (**G**) MA plot showing all nuclear-encoded protein-coding transcripts, including non-IP-enriched transcripts and nuclear-encoded mitochondrial genes (background).

**Fig. S8.**
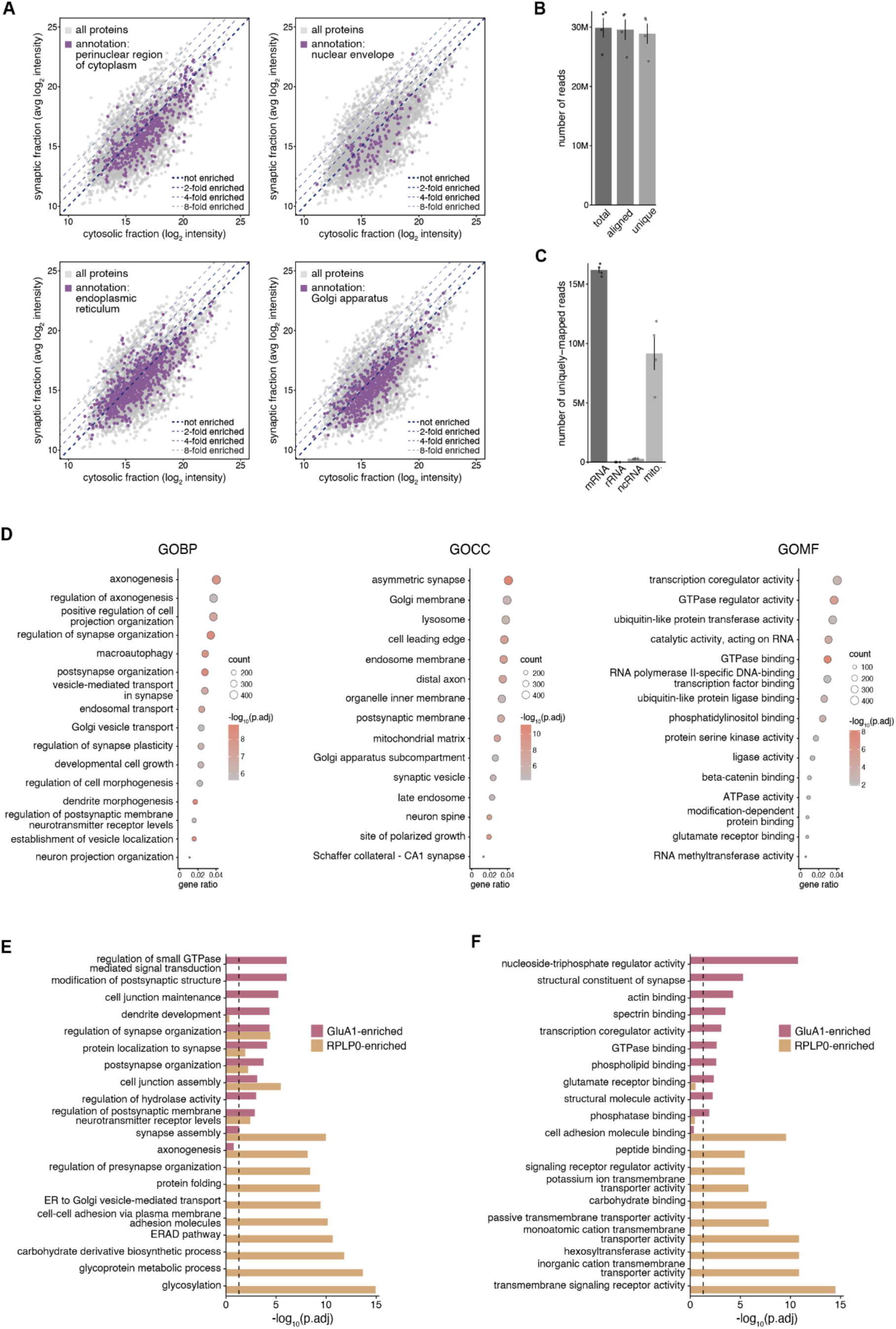
Extended analyses of synapse RNA-seq and ribosome profiling (Ribo-seq) datasets. (**A**) Proteins associated with the GO terms ‘perinuclear region of cytoplasm’, ‘nuclear envelope’, ‘endoplasmic reticulum’, and ‘Golgi apparatus’ are not enriched in the synaptic fraction. (**B**) RNA-seq transcript read counts across sequential filtering steps: total, aligned and uniquely mapped. Error bars show mean ± SEM. (**C**) Number of uniquely-mapped reads assigned to mRNA, rRNA, ncRNA, and mitochondrial genome-encoded transcripts. Error bars show mean ± SEM. (**D**) GO overrepresentation analysis of the 12,283 actively translated synaptic mRNAs shown in Fig. 4B, with all detected transcripts in the synaptic RNA-seq and Ribo-seq datasets used as background. Given the relative depletion of most ER-annotated proteins in the synaptic fraction (fig. S8A), enrichment of membrane-related terms likely reflects synaptic translation of integral membrane proteins rather than contamination by rough ER-associated ribosomes. Similarly, terms such as ‘vesicle-mediated transport in synapse’, ‘Golgi vesicle transport’, and ‘Golgi apparatus subcompartment’ likely reflect active maintenance/translation of local secretory and trafficking organelles rather than contamination with perinuclear ER and canonical Golgi membranes. (**E**) Top 10 enriched GOBP terms for GluA1- vs RPLP0-enriched transcripts. The dashed line indicates -log10(FDR). (**F**) Top 10 enriched GOMF terms for GluA1- vs RPLP0-enriched transcripts. The dashed line indicates -log10(FDR).

**Fig. S9.**
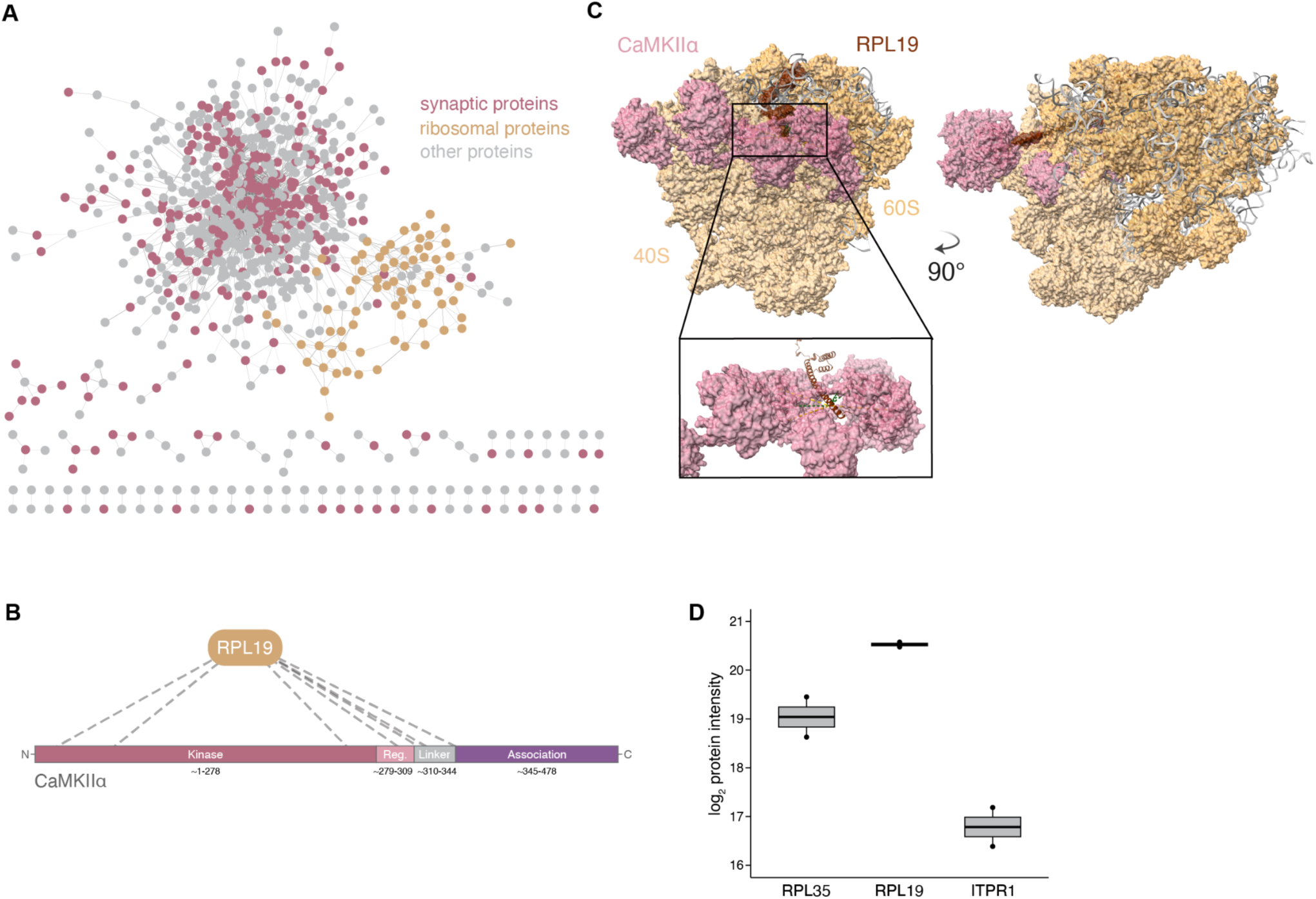
Additional XL-MS analyses. (**A**) Overview of the overall XL-based protein interaction network. Synaptic annotation is based on SynGO. (**B**) Protein domain-level crosslink map between RPL19 and CaMKIIα. Domain classification based on (*72*). (**C**) CaMKIIα-RPL19 interaction model. (**D**) Average log_2_ protein intensities of RPL35, RPL19 and ITPR1 in ribosome-enriched fractions from cortical synaptosomes.

**Fig. S10.**
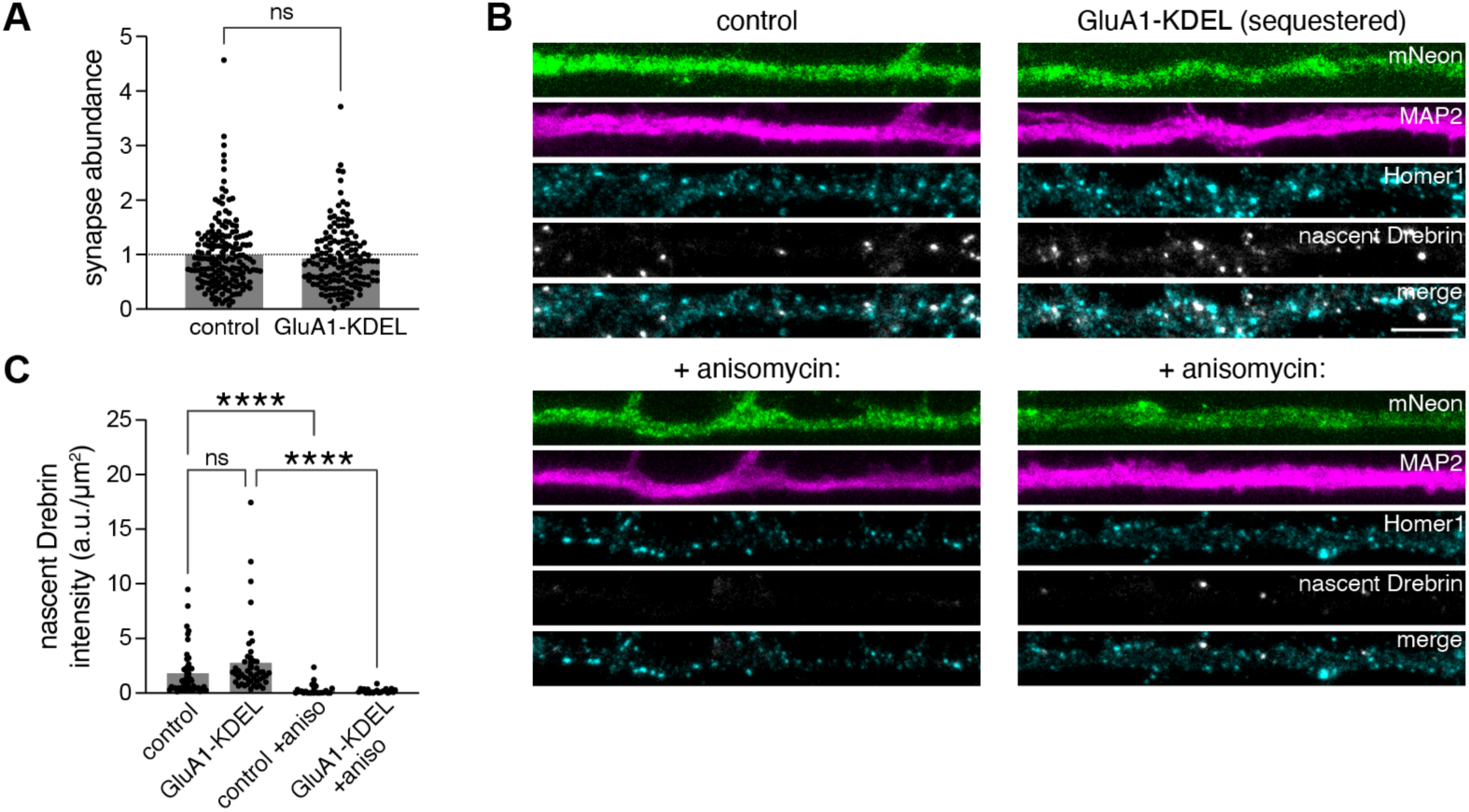
Decreased surface expression of endogenous GluA1 does not affect translation of Dbn1 mRNA within dendrites and synapses. (**A**) Estimate of synapse abundance (synaptic area normalized to the total area) for 50-80 μm dendritic segments from control and GluA1-KDEL expressing neurons. Error bars show mean ± SEM. ns=not significant, two-tailed unpaired t-test; n=146-176 dendritic segments from 146-176 neurons from 2-3 independent cultures. (**B**) Detection of nascent Drebrin in dendrites from control and GluA1-KDEL expressing neurons. Nascent Drebrin was labeled with puromycin (5 min) in the absence or presence of the protein synthesis inhibitor anisomycin (see Methods). Scale bar, 10 µm. (**C**) Quantification of nascent Drebrin in postsynaptic regions from a 50-80 μm dendritic segment from control and GluA1-KDEL expressing neurons. Error bars show mean ± SEM. ****p<0.0001, Brown-Forsythe and Welch’s ANOVA, Dunnett’s multiple comparisons test; n=22-48 dendritic segments from 22-48 neurons from 2 independent cultures.

**Table S1. (separate file)**

Differential expression analyses for determination of compartment-specific ribosome interactomes (related to Fig. 1 and fig. S2)

Sheet 1: Results of the differential expression analysis comparing APEX-NUC to the no-APEX control

Sheet 2: Results of the differential expression analysis comparing APEX-CYTO to the no-APEX control

Sheet 3: Results of the differential expression analysis comparing APEX-SYN to the no-APEX control

**Table S2. (separate file)**

Comparison of the cytosolic and nuclear ribosome interactomes (related to Fig. 1 and fig. S2)

Sheet 1: Results of the differential expression analysis comparing the cytosolic ribosome interactome to the nuclear ribosome interactome

Sheet 2: GO analysis results (filtered for redundancy) for nuclear ribosome-enriched interactors Sheet 3: GO analysis results (filtered for redundancy) for cytosolic ribosome-enriched interactors Sheet 4: Unfiltered GO analysis results for nuclear ribosome-enriched interactors

Sheet 5: Unfiltered GO analysis results for cytosolic ribosome-enriched interactors

**Table S3. (separate file)**

Comparison of the synaptic and non-synaptic ribosome interactomes (related to Fig. 1 and fig. S2)

Sheet 1: Results of the differential expression analysis comparing the synaptic ribosome interactome to the non-synaptic ribosome interactome

Sheet 2: GO analysis results (filtered for redundancy) for synaptic ribosome-enriched interactors

Sheet 3: GO analysis results (filtered for redundancy) for non-synaptic ribosome-enriched interactors

Sheet 4: Unfiltered GO analysis results for synaptic ribosome-enriched interactors Sheet 5: Unfiltered GO analysis results for non-synaptic ribosome-enriched interactors

**Table S4. (separate file)**

Validation of the AMPA receptor-ribosome interaction by reciprocal co-IP from synaptosomes (related to Fig. 2 and fig. S4)

Sheet 1: Results of the differential expression analysis comparing the Ribo IP to the isotype control

Sheet 2: Results of the differential expression analysis comparing the GluA1 IP to the isotype control

Sheet 3: List of proteins (represented as unique gene identifiers) that are unique to the Ribo IP or >3-fold enriched compared to the isotype control

Sheet 4: List of proteins (represented as unique gene identifiers) that are unique to the GluA1 IP or >3-fold enriched compared to the isotype control

Sheet 5: Log_2_ protein intensities of ribosomal proteins across IPs (related to Fig. 2F)

Sheet 6: List of proteins (represented as unique gene identifiers) that are enriched in the APEX PL-MS synaptic ribosome interactome and found in both the Ribo IP and GluA1 IP interactomes

Sheet 7: GO analysis results (filtered for redundancy) for Ribo IP interactors

Sheet 8: GO analysis results (filtered for redundancy) for GluA1 IP interactors

Sheet 9: Unfiltered GO analysis results for Ribo IP interactors

Sheet 10: Unfiltered GO analysis results for GluA1 IP interactors

**Table S5. (separate file)**

Comparison of the synaptic and cytosolic translatomes (related to Fig. 4)

Sheet 1: Results of the differential expression analysis comparing the synaptic and cytosolic translatomes

Sheet 2: Results of the differential expression analysis comparing the RPLP0 IP from the synaptic fraction and the isotype control from the synaptic fraction

Sheet 3: Results of the differential expression analysis comparing the RPLP0 IP from the cytosolic fraction and the isotype control from the cytosolic fraction

Sheet 4: GO analysis results (filtered for redundancy) for synapse-enriched transcripts

Sheet 5: GO analysis results (filtered for redundancy) for cytosol-enriched transcripts

Sheet 6: Unfiltered GO analysis results for synapse-enriched transcripts

Sheet 7: Unfiltered GO analysis results for cytosol-enriched transcripts

**Table S6. (separate file)**

Comparison of synaptic, neuropil, and somata translatomes (related to Fig. 4)

Sheet 1: Combined GO analysis results for synapse-, neuropil-, and somata-enriched transcripts

Sheet 2: GOCC analysis results for rank-increased transcripts

**Table S7. (separate file)**

Comparison of the GluA1-subsynaptic and total synaptic translatomes (related to Fig. 4 and fig. S8)

Sheet 1: Results of the differential expression analysis comparing the GluA1 IP and RPLP0 IP translatomes

Sheet 2: Results of the differential expression analysis comparing the GluA1 IP and the isotype control (rabbit IgG)

Sheet 3: Results of the differential expression analysis comparing the RPLP0 IP and the isotype control (mouse IgG)

Sheet 4: GO analysis results (filtered for redundancy) for GluA1-enriched transcripts

Sheet 5: GO analysis results (filtered for redundancy) for RPLP0-enriched transcripts

Sheet 6: Unfiltered GO analysis results for GluA1-enriched transcripts

Sheet 7: Unfiltered GO analysis results for RPLP0-enriched transcripts

**Table S8. (separate file)**

Analysis of XL-MS data (related to Fig. 5 and fig. S9)

Sheet 1: Results summary

Sheet 2: Original residue-pairs from Scout and pLink3

Sheet 3: List of identified crosslinks before resolving ambiguous peptide assignments

Sheet 4: List of identified crosslinks after resolving ambiguous peptide assignments

Sheet 5: Inter-protein crosslinks between ribosomal proteins

**Movie S1. (separate file)**

Rotating view of the 80S ribosome (PDB 7QGG) highlighting RPL35 (rose) and RPL19 (coral). The yellow segments mark the RPL35/RPL19 peptides identified as crosslinked to CaMKIIα.

## References and Notes

1. M. S. Helm, T. M. Dankovich, S. Mandad, B. Rammner, S. Jähne, V. Salimi, C. Koerbs, R. Leibrandt, H. Urlaub, T. Schikorski, S. O. Rizzoli, A large-scale nanoscopy and biochemistry analysis of postsynaptic dendritic spines. Nat Neurosci 24, 1151–1162 (2021).

2. A. R. Dörrbaum, L. Kochen, J. D. Langer, E. M. Schuman, Local and global influences on protein turnover in neurons and glia. Elife 7, e34202 (2018).

3. E. F. Fornasiero, S. Mandad, H. Wildhagen, M. Alevra, B. Rammner, S. Keihani, F. Opazo, I. Urban, T. Ischebeck, M. S. Sakib, M. K. Fard, K. Kirli, T. P. Centeno, R. O. Vidal, R.-U. Rahman, E. Benito, A. Fischer, S. Dennerlein, P. Rehling, I. Feussner, S. Bonn, M. Simons, H. Urlaub, S. O. Rizzoli, Precisely measured protein lifetimes in the mouse brain reveal differences across tissues and subcellular fractions. Nat Commun 9, 4230 (2018).

4. S. Das, M. Vera, V. Gandin, R. H. Singer, E. Tutucci, Intracellular mRNA transport and localized translation. Nat Rev Mol Cell Biol 22, 483–504 (2021).

5. M. S. Fernandopulle, J. Lippincott-Schwartz, M. E. Ward, RNA transport and local translation in neurodevelopmental and neurodegenerative disease. Nat Neurosci 24, 622–632 (2021).

6. A. M. Bourke, A. Schwarz, E. M. Schuman, De-centralizing the Central Dogma: mRNA translation in space and time. Mol Cell 83, 452–468 (2023).

7. M. M. Poon, S.-H. Choi, C. A. M. Jamieson, D. H. Geschwind, K. C. Martin, Identification of process-localized mRNAs from cultured rodent hippocampal neurons. J Neurosci 26, 13390–13399 (2006).

8. I. J. Cajigas, G. Tushev, T. J. Will, S. tom Dieck, N. Fuerst, E. M. Schuman, The local transcriptome in the synaptic neuropil revealed by deep sequencing and high-resolution imaging. Neuron 74, 453–466 (2012).

9. A. Biever, C. Glock, G. Tushev, E. Ciirdaeva, T. Dalmay, J. D. Langer, E. M. Schuman, Monosomes actively translate synaptic mRNAs in neuronal processes. Science 367, eaay4991 (2020).

10. C. Glock, A. Biever, G. Tushev, B. Nassim-Assir, A. Kao, I. Bartnik, S. Tom Dieck, E. M. Schuman, The translatome of neuronal cell bodies, dendrites, and axons. Proc Natl Acad Sci U S A 118, e2113929118 (2021).

11. A. Zappulo, D. van den Bruck, C. Ciolli Mattioli, V. Franke, K. Imami, E. McShane, M. Moreno-Estelles, L. Calviello, A. Filipchyk, E. Peguero-Sanchez, T. Müller, A. Woehler, C. Birchmeier, E. Merino, N. Rajewsky, U. Ohler, E. O. Mazzoni, M. Selbach, A. Akalin, M. Chekulaeva, RNA localization is a key determinant of neurite-enriched proteome. Nat Commun 8, 583 (2017).

12. K. H. Zivraj, Y. C. L. Tung, M. Piper, L. Gumy, J. W. Fawcett, G. S. H. Yeo, C. E. Holt, Subcellular profiling reveals distinct and developmentally regulated repertoire of growth cone mRNAs. J Neurosci 30, 15464–15478 (2010).

13. D. Bodian, A SUGGESTIVE RELATIONSHIP OF NERVE CELL RNA WITH SPECIFIC SYNAPTIC SITES. Proc Natl Acad Sci U S A 53, 418–425 (1965).

14. O. Steward, W. B. Levy, Preferential localization of polyribosomes under the base of dendritic spines in granule cells of the dentate gyrus. J Neurosci 2, 284–291 (1982).

15. O. Steward, Alterations in polyribosomes associated with dendritic spines during the reinnervation of the dentate gyrus of the adult rat. J Neurosci 3, 177–188 (1983).

16. L. E. Ostroff, J. C. Fiala, B. Allwardt, K. M. Harris, Polyribosomes redistribute from dendritic shafts into spines with enlarged synapses during LTP in developing rat hippocampal slices. Neuron 35, 535–545 (2002).

17. C. Sun, A. Nold, C. M. Fusco, V. Rangaraju, T. Tchumatchenko, M. Heilemann, E. M. Schuman, The prevalence and specificity of local protein synthesis during neuronal synaptic plasticity. Sci Adv 7, eabj0790 (2021).

18. O. Steward, C. S. Wallace, G. L. Lyford, P. F. Worley, Synaptic activation causes the mRNA for the IEG Arc to localize selectively near activated postsynaptic sites on dendrites. Neuron 21, 741–751 (1998).

19. Y. J. Yoon, B. Wu, A. R. Buxbaum, S. Das, A. Tsai, B. P. English, J. B. Grimm, L. D. Lavis, R. H. Singer, Glutamate-induced RNA localization and translation in neurons. Proc Natl Acad Sci U S A 113, E6877–E6886 (2016).

20. J. N. Bourne, K. E. Sorra, J. Hurlburt, K. M. Harris, Polyribosomes are increased in spines of CA1 dendrites 2 h after the induction of LTP in mature rat hippocampal slices. Hippocampus 17, 1–4 (2007).

21. L. E. Ostroff, D. J. Watson, G. Cao, P. H. Parker, H. Smith, K. M. Harris, Shifting patterns of polyribosome accumulation at synapses over the course of hippocampal long-term potentiation. Hippocampus 28, 416–430 (2018).

22. L. E. Ostroff, C. K. Cain, J. Bedont, M. H. Monfils, J. E. Ledoux, Fear and safety learning differentially affect synapse size and dendritic translation in the lateral amygdala. Proc Natl Acad Sci U S A 107, 9418–9423 (2010).

23. J. Tcherkezian, P. A. Brittis, F. Thomas, P. P. Roux, J. G. Flanagan, Transmembrane receptor DCC associates with protein synthesis machinery and regulates translation. Cell 141, 632–644 (2010).

24. M. Koppers, R. Cagnetta, T. Shigeoka, L. C. Wunderlich, P. Vallejo-Ramirez, J. Qiaojin Lin, S. Zhao, M. A. Jakobs, A. Dwivedy, M. S. Minett, A. Bellon, C. F. Kaminski, W. A. Harris, J. G. Flanagan, C. E. Holt, Receptor-specific interactome as a hub for rapid cue-induced selective translation in axons. Elife 8, e48718 (2019).

25. C. M. Fusco, K. Desch, A. R. Dörrbaum, M. Wang, A. Staab, I. C. W. Chan, E. Vail, V. Villeri, J. D. Langer, E. M. Schuman, Neuronal ribosomes exhibit dynamic and context-dependent exchange of ribosomal proteins. Nat Commun 12, 6127 (2021).

26. H. Imai, D. Utsumi, H. Torihara, K. Takahashi, H. Kuroyanagi, A. Yamashita, Simultaneous measurement of nascent transcriptome and translatome using 4-thiouridine metabolic RNA labeling and translating ribosome affinity purification. Nucleic Acids Res 51, e76 (2023).

27. J. Schwenk, N. Harmel, A. Brechet, G. Zolles, H. Berkefeld, C. S. Müller, W. Bildl, D. Baehrens, B. Hüber, A. Kulik, N. Klöcker, U. Schulte, B. Fakler, High-resolution proteomics unravel architecture and molecular diversity of native AMPA receptor complexes. Neuron 74, 621–633 (2012).

28. R. F. Gesteland, Unfolding of Escherichia coli ribosomes by removal of magnesium. J Mol Biol 18, 356–371 (1966).

29. H. Tiedge, J. Brosius, Translational machinery in dendrites of hippocampal neurons in culture. J Neurosci 16, 7171–7181 (1996).

30. A. Gardiol, C. Racca, A. Triller, Dendritic and postsynaptic protein synthetic machinery. J Neurosci 19, 168–179 (1999).

31. G. Aakalu, W. B. Smith, N. Nguyen, C. Jiang, E. M. Schuman, Dynamic visualization of local protein synthesis in hippocampal neurons. Neuron 30, 489–502 (2001).

32. Y. Fukata, A. Dimitrov, G. Boncompain, O. Vielemeyer, F. Perez, M. Fukata, Local palmitoylation cycles define activity-regulated postsynaptic subdomains. J Cell Biol 202, 145–161 (2013).

33. D. Nair, E. Hosy, J. D. Petersen, A. Constals, G. Giannone, D. Choquet, J.-B. Sibarita, Super-resolution imaging reveals that AMPA receptors inside synapses are dynamically organized in nanodomains regulated by PSD95. J Neurosci 33, 13204–13224 (2013).

34. H. D. MacGillavry, Y. Song, S. Raghavachari, T. A. Blanpied, Nanoscale scaffolding domains within the postsynaptic density concentrate synaptic AMPA receptors. Neuron 78, 615–622 (2013).

35. L. E. Ostroff, B. Botsford, S. Gindina, K. K. Cowansage, J. E. LeDoux, E. Klann, C. Hoeffer, Accumulation of Polyribosomes in Dendritic Spine Heads, But Not Bases and Necks, during Memory Consolidation Depends on Cap-Dependent Translation Initiation. J Neurosci 37, 1862–1872 (2017).

36. K. Simbriger, I. S. Amorim, K. Chalkiadaki, G. Lach, S. M. Jafarnejad, A. Khoutorsky, C. G. Gkogkas, Monitoring translation in synaptic fractions using a ribosome profiling strategy. J Neurosci Methods 329, 108456 (2020).

37. C. Bernard, D. Exposito-Alonso, M. Selten, S. Sanalidou, A. Hanusz-Godoy, A. Aguilera, F. Hamid, F. Oozeer, P. Maeso, L. Allison, M. Russell, R. A. Fleck, B. Rico, O. Marín, Cortical wiring by synapse type-specific control of local protein synthesis. Science 378, eabm7466 (2022).

38. N. T. Ingolia, S. Ghaemmaghami, J. R. S. Newman, J. S. Weissman, Genome-wide analysis in vivo of translation with nucleotide resolution using ribosome profiling. Science 324, 218–223 (2009).

39. S. C. Harward, N. G. Hedrick, C. E. Hall, P. Parra-Bueno, T. A. Milner, E. Pan, T. Laviv, B. L. Hempstead, R. Yasuda, J. O. McNamara, Autocrine BDNF-TrkB signalling within a single dendritic spine. Nature 538, 99–103 (2016).

40. W.-H. Lu, T.-T. Chang, Y.-M. Chang, Y.-H. Liu, C.-H. Lin, C.-S. Suen, M.-J. Hwang, Y.- S. Huang, CPEB2-activated axonal translation of VGLUT2 mRNA promotes glutamatergic transmission and presynaptic plasticity. J Biomed Sci 31, 69 (2024).

41. M. A. Gonzalez-Lozano, F. Koopmans, P. F. Sullivan, J. Protze, G. Krause, M. Verhage, K. W. Li, F. Liu, A. B. Smit, Stitching the synapse: Cross-linking mass spectrometry into resolving synaptic protein interactions. Sci Adv 6, eaax5783 (2020).

42. K. U. Bayer, H. Schulman, CaM Kinase: Still Inspiring at 40. Neuron 103, 380–394 (2019).

43. J. B. Myers, V. Zaegel, S. J. Coultrap, A. P. Miller, K. U. Bayer, S. L. Reichow, The CaMKII holoenzyme structure in activation-competent conformations. Nat Commun 8, 15742 (2017).

44. P. Opazo, S. Labrecque, C. M. Tigaret, A. Frouin, P. W. Wiseman, P. De Koninck, D. Choquet, CaMKII triggers the diffusional trapping of surface AMPARs through phosphorylation of stargazin. Neuron 67, 239–252 (2010).

45. A. S. Leonard, I. A. Lim, D. E. Hemsworth, M. C. Horne, J. W. Hell, Calcium/calmodulin-dependent protein kinase II is associated with the N-methyl-D-aspartate receptor. Proc Natl Acad Sci U S A 96, 3239–3244 (1999).

46. A. S. Kristensen, M. A. Jenkins, T. G. Banke, A. Schousboe, Y. Makino, R. C. Johnson, R. Huganir, S. F. Traynelis, Mechanism of Ca2+/calmodulin-dependent kinase II regulation of AMPA receptor gating. Nat Neurosci 14, 727–735 (2011).

47. A. David, B. P. Dolan, H. D. Hickman, J. J. Knowlton, G. Clavarino, P. Pierre, J. R. Bennink, J. W. Yewdell, Nuclear translation visualized by ribosome-bound nascent chain puromycylation. J Cell Biol 197, 45–57 (2012).

48. M. Sumi, K. Kiuchi, T. Ishikawa, A. Ishii, M. Hagiwara, T. Nagatsu, H. Hidaka, The newly synthesized selective Ca2+/calmodulin dependent protein kinase II inhibitor KN-93 reduces dopamine contents in PC12h cells. Biochem Biophys Res Commun 181, 968–975 (1991).

49. A. Ishida, I. Kameshita, S. Okuno, T. Kitani, H. Fujisawa, A novel highly specific and potent inhibitor of calmodulin-dependent protein kinase II. Biochem Biophys Res Commun 212, 806–812 (1995).

50. C. N. Brown, K. U. Bayer, Studying CaMKII: Tools and standards. Cell Rep 43, 113982 (2024).

51. K. U. Bayer, E. LeBel, G. L. McDonald, H. O’Leary, H. Schulman, P. De Koninck, Transition from reversible to persistent binding of CaMKII to postsynaptic sites and NR2B. J Neurosci 26, 1164–1174 (2006).

52. P. Mullasseril, A. Dosemeci, J. E. Lisman, L. C. Griffith, A structural mechanism for maintaining the “on-state” of the CaMKII memory switch in the post-synaptic density. J Neurochem 103, 357–364 (2007).

53. R. S. Vest, H. O’Leary, S. J. Coultrap, M. S. Kindy, K. U. Bayer, Effective post-insult neuroprotection by a novel Ca(2+)/ calmodulin-dependent protein kinase II (CaMKII) inhibitor. J Biol Chem 285, 20675–20682 (2010).

54. D. J. Kareemo, C. S. Winborn, S. S. Olah, C. N. Miller, J. Kim, C. A. Kadgien, H. S. Actor-Engel, H. J. Ramsay, A. M. Ramsey, J. Aoto, M. J. Kennedy, Genetically encoded intrabody probes for labeling and manipulating AMPA-type glutamate receptors. Nat Commun 15, 10374 (2024).

55. S. tom Dieck, L. Kochen, C. Hanus, M. Heumüller, I. Bartnik, B. Nassim-Assir, K. Merk, T. Mosler, S. Garg, S. Bunse, D. A. Tirrell, E. M. Schuman, Direct visualization of newly synthesized target proteins in situ. Nat Methods 12, 411–414 (2015).

56. M. Juengling, J. Mosbacher, J. Perez, M. Van Oostrum, E. Kaulich, S. Tom Dieck, N. Fuerst, G. Tushev, M. Korelidou, E. M. Schuman, Brain-wide synaptosome profiling reveals localized mRNAs that diversify synapses. Neuroscience [Preprint] (2025). 10.64898/2025.12.07.692806.

57. R. Cagnetta, C. K. Frese, T. Shigeoka, J. Krijgsveld, C. E. Holt, Rapid Cue-Specific Remodeling of the Nascent Axonal Proteome. Neuron 99, 29–46.e4 (2018).

58. T. Shigeoka, H. Jung, J. Jung, B. Turner-Bridger, J. Ohk, J. Q. Lin, P. S. Amieux, C. E. Holt, Dynamic Axonal Translation in Developing and Mature Visual Circuits. Cell 166, 181–192 (2016).

59. B. D. Hobson, L. Kong, M. F. Angelo, O. J. Lieberman, E. V. Mosharov, E. Herzog, D. Sulzer, P. A. Sims, Subcellular and regional localization of mRNA translation in midbrain dopamine neurons. Cell Rep 38, 110208 (2022).

60. E. Hacisuleyman, C. R. Hale, N. Noble, J.-D. Luo, J. J. Fak, M. Saito, J. Chen, J. S. Weissman, R. B. Darnell, Neuronal activity rapidly reprograms dendritic translation via eIF4G2:uORF binding. Nat Neurosci 27, 822–835 (2024).

61. K. Shen, T. Meyer, Dynamic control of CaMKII translocation and localization in hippocampal neurons by NMDA receptor stimulation. Science 284, 162–166 (1999).

62. J. E. Lisman, A. M. Zhabotinsky, A model of synaptic memory: a CaMKII/PP1 switch that potentiates transmission by organizing an AMPA receptor anchoring assembly. Neuron 31, 191–201 (2001).

63. S. Tomita, H. Adesnik, M. Sekiguchi, W. Zhang, K. Wada, J. R. Howe, R. A. Nicoll, D. S. Bredt, Stargazin modulates AMPA receptor gating and trafficking by distinct domains. Nature 435, 1052–1058 (2005).

64. J. E. Tullis, M. E. Larsen, N. L. Rumian, R. K. Freund, E. E. Boxer, C. N. Brown, S. J. Coultrap, H. Schulman, J. Aoto, M. L. Dell’Acqua, K. U. Bayer, LTP induction by structural rather than enzymatic functions of CaMKII. Nature 621, 146–153 (2023).

65. N. L. Rumian, C. M. Barker, M. E. Larsen, J. E. Tullis, R. K. Freund, A. Taslimi, S. J. Coultrap, C. L. Tucker, M. L. Dell’Acqua, K. U. Bayer, LTP expression mediated by autonomous activity of GluN2B-bound CaMKII. Cell Rep 43, 114866 (2024).

66. W. J. Oh, C. Wu, S. J. Kim, V. Facchinetti, L.-A. Julien, M. Finlan, P. P. Roux, B. Su, E. Jacinto, mTORC2 can associate with ribosomes to promote cotranslational phosphorylation and stability of nascent Akt polypeptide. EMBO J 29, 3939–3951 (2010).

67. B. Bingol, C.-F. Wang, D. Arnott, D. Cheng, J. Peng, M. Sheng, Autophosphorylated CaMKIIalpha acts as a scaffold to recruit proteasomes to dendritic spines. Cell 140, 567–578 (2010).

68. D. Meyer, T. Bonhoeffer, V. Scheuss, Balance and stability of synaptic structures during synaptic plasticity. Neuron 82, 430–443 (2014).

69. M. Bosch, J. Castro, T. Saneyoshi, H. Matsuno, M. Sur, Y. Hayashi, Structural and molecular remodeling of dendritic spine substructures during long-term potentiation. Neuron 82, 444–459 (2014).

70. A.-H. Tang, H. Chen, T. P. Li, S. R. Metzbower, H. D. MacGillavry, T. A. Blanpied, A trans-synaptic nanocolumn aligns neurotransmitter release to receptors. Nature 536, 210–214 (2016).

71. Y. Perez-Riverol, C. Bandla, D. J. Kundu, S. Kamatchinathan, J. Bai, S. Hewapathirana, N. S. John, A. Prakash, M. Walzer, S. Wang, J. A. Vizcaíno, The PRIDE database at 20 years: 2025 update. Nucleic Acids Res 53, D543–D553 (2025).

72. K. U. Bayer, K. P. Giese, A revised view of the role of CaMKII in learning and memory. Nat Neurosci 28, 24–34 (2025).

73. K. Desch, E. M. Schuman, J. D. Langer, Quantifying phosphorylation dynamics in primary neuronal cultures using LC-MS/MS. STAR Protoc 3, 101063 (2022).

74. C. T. Rueden, J. Schindelin, M. C. Hiner, B. E. DeZonia, A. E. Walter, E. T. Arena, K. W. Eliceiri, ImageJ2: ImageJ for the next generation of scientific image data. BMC Bioinformatics 18, 529 (2017).

75. K. Desch, J. D. Langer, E. M. Schuman, Dynamic bi-directional phosphorylation events associated with the reciprocal regulation of synapses during homeostatic up- and down-scaling. Cell Rep 36, 109583 (2021).

76. J. Rappsilber, M. Mann, Y. Ishihama, Protocol for micro-purification, enrichment, pre-fractionation and storage of peptides for proteomics using StageTips. Nat Protoc 2, 1896–1906 (2007).

77. J. Muntel, J. Kirkpatrick, R. Bruderer, T. Huang, O. Vitek, A. Ori, L. Reiter, Comparison of Protein Quantification in a Complex Background by DIA and TMT Workflows with Fixed Instrument Time. J Proteome Res 18, 1340–1351 (2019).

78. V. Demichev, C. B. Messner, S. I. Vernardis, K. S. Lilley, M. Ralser, DIA-NN: neural networks and interference correction enable deep proteome coverage in high throughput. Nat Methods 17, 41–44 (2020).

79. F. Koopmans, K. W. Li, R. V. Klaassen, A. B. Smit, MS-DAP Platform for Downstream Data Analysis of Label-Free Proteomics Uncovers Optimal Workflows in Benchmark Data Sets and Increased Sensitivity in Analysis of Alzheimer’s Biomarker Data. J Proteome Res 22, 374–386 (2023).

80. Laurent Gatto, Christophe Vanderaa, QFeatures, Bioconductor; 10.18129/B9.BIOC.QFEATURES.

81. V. O. Martin Morgan, SummarizedExperiment, Bioconductor (2017); 10.18129/B9.BIOC.SUMMARIZEDEXPERIMENT.

82. S. Hediyeh-Zadeh, A. I. Webb, M. J. Davis, MsImpute: Estimation of Missing Peptide Intensity Data in Label-Free Quantitative Mass Spectrometry. Mol Cell Proteomics 22, 100558 (2023).

83. A. Sticker, L. Goeminne, L. Martens, L. Clement, Robust Summarization and Inference in Proteome-wide Label-free Quantification. Mol Cell Proteomics 19, 1209–1219 (2020).

84. T. Wu, E. Hu, S. Xu, M. Chen, P. Guo, Z. Dai, T. Feng, L. Zhou, W. Tang, L. Zhan, X. Fu, S. Liu, X. Bo, G. Yu, clusterProfiler 4.0: A universal enrichment tool for interpreting omics data. Innovation (Camb) 2, 100141 (2021).

85. S. Sayols, rrvgo: a Bioconductor package for interpreting lists of Gene Ontology terms. MicroPubl Biol 2023 (2023).

86. G. Yu, F. Li, Y. Qin, X. Bo, Y. Wu, S. Wang, GOSemSim: an R package for measuring semantic similarity among GO terms and gene products. Bioinformatics 26, 976–978 (2010).

87. Y. Liao, G. K. Smyth, W. Shi, featureCounts: an efficient general purpose program for assigning sequence reads to genomic features. Bioinformatics 30, 923–930 (2014).

88. L. Ferguson, H. E. Upton, S. C. Pimentel, A. Mok, L. F. Lareau, K. Collins, N. T. Ingolia, Streamlined and sensitive mono- and di-ribosome profiling in yeast and human cells. Nat Methods 20, 1704–1715 (2023).

89. F. Lauria, T. Tebaldi, P. Bernabò, E. J. N. Groen, T. H. Gillingwater, G. Viero, riboWaltz: Optimization of ribosome P-site positioning in ribosome profiling data. PLoS Comput Biol 14, e1006169 (2018).

90. D. Karolchik, A. S. Hinrichs, T. S. Furey, K. M. Roskin, C. W. Sugnet, D. Haussler, W. J. Kent, The UCSC Table Browser data retrieval tool. Nucleic Acids Res 32, D493–496 (2004).

91. A. R. Quinlan, I. M. Hall, BEDTools: a flexible suite of utilities for comparing genomic features. Bioinformatics 26, 841–842 (2010).

92. J. M. Rodriguez, J. Rodriguez-Rivas, T. Di Domenico, J. Vázquez, A. Valencia, M. L. Tress, APPRIS 2017: principal isoforms for multiple gene sets. Nucleic Acids Res 46, D213–D217 (2018).

93. C. W. Law, Y. Chen, W. Shi, G. K. Smyth, voom: Precision weights unlock linear model analysis tools for RNA-seq read counts. Genome Biol 15, R29 (2014).

94. M. E. Ritchie, B. Phipson, D. Wu, Y. Hu, C. W. Law, W. Shi, G. K. Smyth, limma powers differential expression analyses for RNA-sequencing and microarray studies. Nucleic Acids Res 43, e47 (2015).

95. M. D. Robinson, D. J. McCarthy, G. K. Smyth, edgeR: a Bioconductor package for differential expression analysis of digital gene expression data. Bioinformatics 26, 139–140 (2010).

96. C. S. Hughes, S. Moggridge, T. Müller, P. H. Sorensen, G. B. Morin, J. Krijgsveld, Single-pot, solid-phase-enhanced sample preparation for proteomics experiments. Nat Protoc 14, 68–85 (2019).

97. P.-L. Jiang, Y. Zhu, J. Cai, C. Wang, M. Wu, K. Pu, F. Liu, Optimized In-Solution and Gas-Phase Chemistry Enables High-Efficiency Interactome Mapping by DSBSO-Based Cross-Linking Mass Spectrometry. Angew Chem Int Ed Engl 65, e18355 (2026).

98. R. V. Honorato, M. E. Trellet, B. Jiménez-García, J. J. Schaarschmidt, M. Giulini, V. Reys, P. I. Koukos, J. P. G. L. M. Rodrigues, E. Karaca, G. C. P. van Zundert, J. Roel-Touris, C. W. van Noort, Z. Jandová, A. S. J. Melquiond, A. M. J. J. Bonvin, The HADDOCK2.4 web server for integrative modeling of biomolecular complexes. Nat Protoc 19, 3219–3241 (2024).

